# Back-propagation learning in deep Spike-By-Spike networks

**DOI:** 10.1101/569236

**Authors:** David Rotermund, Klaus R. Pawelzik

**Author notes:** Correspondence: David Rotermund.

## Abstract

Neural networks are important building blocks in technical applications. These artificial neural networks (ANNs) rely on noiseless continuous signals in stark contrast to the discrete action potentials stochastically exchanged among the neurons in real brains. A promising approach towards bridging this gap are the Spike-by-Spike (SbS) networks which represent a compromise between non-spiking and spiking versions of generative models that perform inference on their inputs. What is still missing are algorithms for finding weight sets that would optimize the output performances of deep SbS networks with many layers.

Here, a learning rule for hierarchically organized SbS networks is derived. The properties of this approach are investigated and its functionality demonstrated by simulations. In particular, a Deep Convolutional SbS network for classifying handwritten digits (MNIST) is presented. When applied together with an optimizer this learning method achieves a classification performance of roughly 99.3% on the MNIST test data. Thereby it approaches the benchmark results of ANNs without extensive parameter optimization. We envision that with this learning rule SBS networks will provide a new basis for research in neuroscience and for technical applications, especially when they become implemented on specialized computational hardware.

## 1 INTRODUCTION

Fueled by the huge improvements in computational power by CPUs, GPUs, and special hardware (Sze et al., 2017; Jouppi et al., 2018; Lacey et al., 2016), deep neuronal networks (Schmidhuber, 2015) brought a massive improvement for the field of expert systems as well as artificial intelligence (Guo et al., 2016; Azkarate Saiz, 2015; Silver et al., 2016; Mnih et al., 2015; Gatys et al., 2016). These networks started out as simple perceptrons (Rosenblatt, 1958) which where extended into multi-layer networks by a learning rule that utilizes the chain rule to propagate the error, between the actual and the desired output of the network, back from the output layer to the input. This allows to train all weights in such a network based on this back-propagated error. This learning rule is called backprop (Rumelhart et al., 1986).

In real biological neuronal networks, however, information typically is exchanged between neurons by discrete stereotyped signals, the action potentials. Combining deep networks with spikes can open the door for new opportunities (Tavanaei et al., 2018; Lee et al., 2016; Anwani and Rajendran, 2018; Wu et al., 2018), among which are biologically more realistic neuronal networks for studying and describing the information processing in the brain as well as interesting technical approaches for improving the operation of such networks (e.g. low power consumption, fast inference, event-driven information processing, and massive parallelization) (Pfeiffer and Pfeil, 2018).

Ernst et al. (2007) presented a type of shallow neuronal network that is based on non-negative generative models (Lee and Seung, 1999, 2001). Since here time progresses only from one spike to the next in this Spike-By-Spike (SbS) framework, it comes with relatively low additional computational requirements for using spikes as a mean for transmitting information to other neurons.

This makes SbS akin to event-based neuron models (Brette, 2006, 2007; Serrano-Gotarredona et al., 2015; Lagorce et al., 2015) but with populations of stochastically firing neurons that perform inference. While the goal of these inference populations (IPs) is to represent the input as best as possible by their latent variables also their sparseness becomes optimized: An IP has one latent variable *h*(*i*) for each of its *N* neurons (*i* = 1, …, *N*) and the update of the latent variables (here termed h-dynamic) finds solutions where the *h*(*i*)’s are sparse over the populations of neurons. The level of sparseness is influenced by a parameter *ϵ* (Ernst et al., 2007) that represents the temporal integration rate and benefits from using non-negative elements. Using non-negative elements is connected to compressed sensing (CS, (Bruckstein et al., 2008; Candes et al., 2006; Lustig et al., 2008; Wiedemann et al., 2018; Ganguli and Sompolinsky, 2012, 2010)), a method used in technical applications to reconstruct underlying causes from data if these causes are sparse.

Furthermore, SbS networks allow for massive parallelization. In particular, in layered architectures each layer consists of many IPs which can be simulated independently while the communication between the IPs is organized by a low bandwidth signal – the spikes –. While this natural parallelization into populations or single neurons is a property of most neuronal networks with spiking neurons, the required bandwidth for using a typical spiking neuron model is higher when for integration time needs to be segmented into small time steps. In the case of the SbS model, where time progresses from one spike to the next, only the identity of the spiking neuron and information about the corresponding IP is transferred. Technically speaking, this allows to build special hardware (Rotermund and Pawelzik, 2018) dealing with the SbS IPs where one application specific integrated circuit (ASIC) can host a sub-network of IPs which can be arranged into larger networks by connecting such ASICs via exchanging the low bandwidth spike information. This is furthered by the ability to run the internal h-dynamics of an IP asynchronously from the spike exchange process.

In summary, a SbS network is fundamentally different from usual artificial neuronal networks since a.) the building block of the network are SbS inference populations which realize an optimized generative representation of the IP’s input with only non-negative elements, b.) time progresses from one spike to the next, while conserving the property of stochastically firing neurons, and c.) a SbS network has only a small number of parameters, making it easy to use. With respect to biological realism and computational effort to simulate them these properties place a SbS network in between non-spiking neural networks and networks of stochastically spiking neurons.

However, until now constructing deep SbS networks was not possible since only local learning rules for the weights existed.

For convenience we here recapitulate the inner workings of one SbS inference population (Ernst et al., 2007) by taking a look at the most simple network, constructed from one input population and one hidden inference population (see figure 1a): For setting up the framework we initially assume that the input neurons fire independently according a Poisson distribution with a firing rate *ρ*_*µ*_(*s*), where *s* denoted one of the *N*_*S*_ input neurons and *µ* the momentary input pattern from a set of *M* available input patterns.

**Figure 1.**
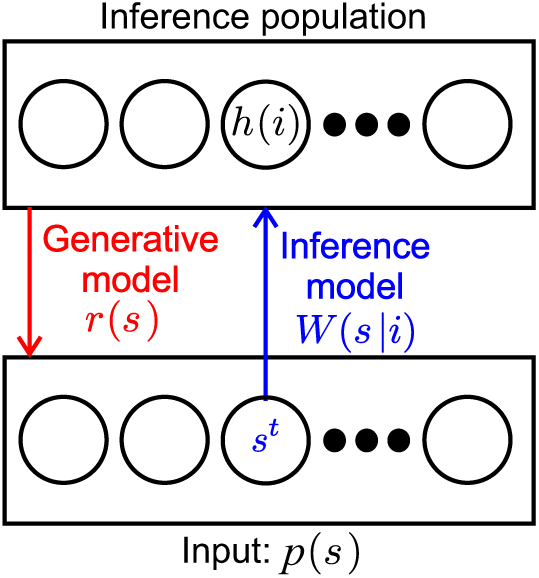
SbS inference population (IPs). SbS IP that processes incoming stochastic spikes *s*^*t*^ generated from an input population *p*(*s*), where *s* denotes the input neuron. The non-negative weights *W* (*s*|*i*) and the latent variables *h*(*i*) of the neurons (*i* enumerates the *N*_*H*_ neurons in that IP) are used in a generative model to optimally represent the observed input.

The input pattern can be e.g. pixel images, waveforms or other types of rate coded information. The normalized external input 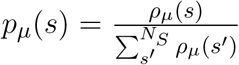 is the probability of input neuron *s*_*t*_ to be the next to fire a spike where *t* denotes the time step when this happens.

The inference population has one latent variable *h*_*µ*_(*i*) for each of its neurons, where *i* denotes the identity of the neuron. The latent variables of a IP can form a probability distribution with 0 ≤ *h*(*i*) ≤ 1 since they are normalized according to 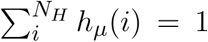. The purpose of representing its input as a generative model with its latent variables is laid upon the inference population. The internal representation *r*_*µ*_(*s*) of the normalized external input 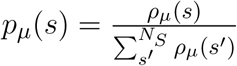 is defined by

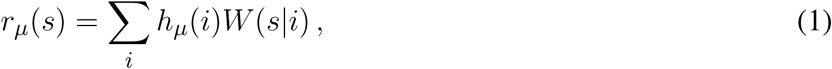

where *W* (*s*|*i*) are the corresponding weights between the input population and the hidden population, which are also normalized and non-negative numbers (0 ≤ *W* (*s*|*i*) ≤ 1 with ∑_*s*_ *W* (*s*|*i*) = 1).

*p*_*µ*_(*s*) can not be observed directly by the inference population. It can only see the spikes *s*^*t*^ emitted by the input population. By counting the spikes 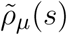 can be estimated through

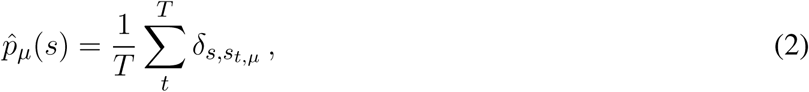

after observing *T* spikes. The goal of the IP is to minimize the difference between 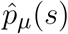 and its own representation *r*_*µ*_(*s*). As measure the cross-entropy can be used:

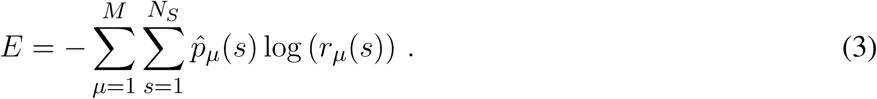

*E* can now be derived for *h*_*µ*_(*i*) for producing an iterative algorithm for finding the optimal *h*_*µ*_(*i*) for given weights *W* (*s*|*i*) and observed spike pattern 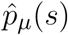.

The iterative update algorithm for the latent variables *h*_*µ*_(*i*) is simplified by processing only one spike 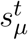 at a time. This results in the so called h-dynamic (Ernst et al., 2007):

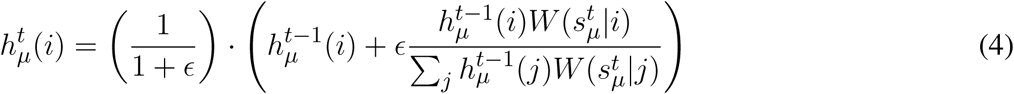

with *ϵ* > 0 as smoothing constant (i.e. a rate constant in a low-pass filter that regulates how much impact the contribution for one spike has onto the latent variables) which also results in different level of sparseness on *h*_*µ*_(*i*) depending of its value (the larger *ϵ* the sparser the representation).

The transition to processing only the momentary spike in the h-dynamic also allows to replace the original Poisson rate coded input populations by Bernoulli processes based on the probability distribution *p*_*µ*_(*s*). This is possible because now the only important information is which one of the input neuron fires next.

This simple example with one input population and one SbS inference population can be extended to all kind of network structures (Rotermund and Pawelzik, 2019b) by exploiting the fact that the latent variables of an IP themselves form a probability distribution 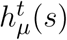. This distribution is then used in a Bernoulli process to also produce spikes and act as an input population for other IPs. While the *p*_*µ*_(*s*) of the input is assumed to be constant over time for a given pattern, 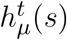 will change with every spike that IP processes. This allows to transport information through the whole network.

While Ernst et al. (2007) provides suitable learning rules for networks with one hidden layer, it doesn’t offer learning rules for networks with arbitrary many layers. In this paper a learning rule capable of training the weights for much deeper networks is derived in analogy to error back-propagation (Rumelhart et al., 1986) but takes into account the update rule for the latent variables presented above. After that, the functionality of this new learning rule is demonstrated through several examples with increasing complexity. Finally a deep convolutional network used for classifying handwritten digits is examined where we also introduce an optimizer which significantly improves the network’s performance as compared to using only a simple gradient descent method.

## 2 RESULTS

The source code used for simulating the presented networks can be found in the supplemental materials.

### 2.1 SbS backprop learning rule

In the following a short summary of the learning rule is given. A detailed derivation of the learning rule can be found in the supplemental material. Since the equations are heavily laden with indices, figure 2 gives an overview of the equations for an example segment of a network with an output layer and three hidden layers.

**Figure 2.**
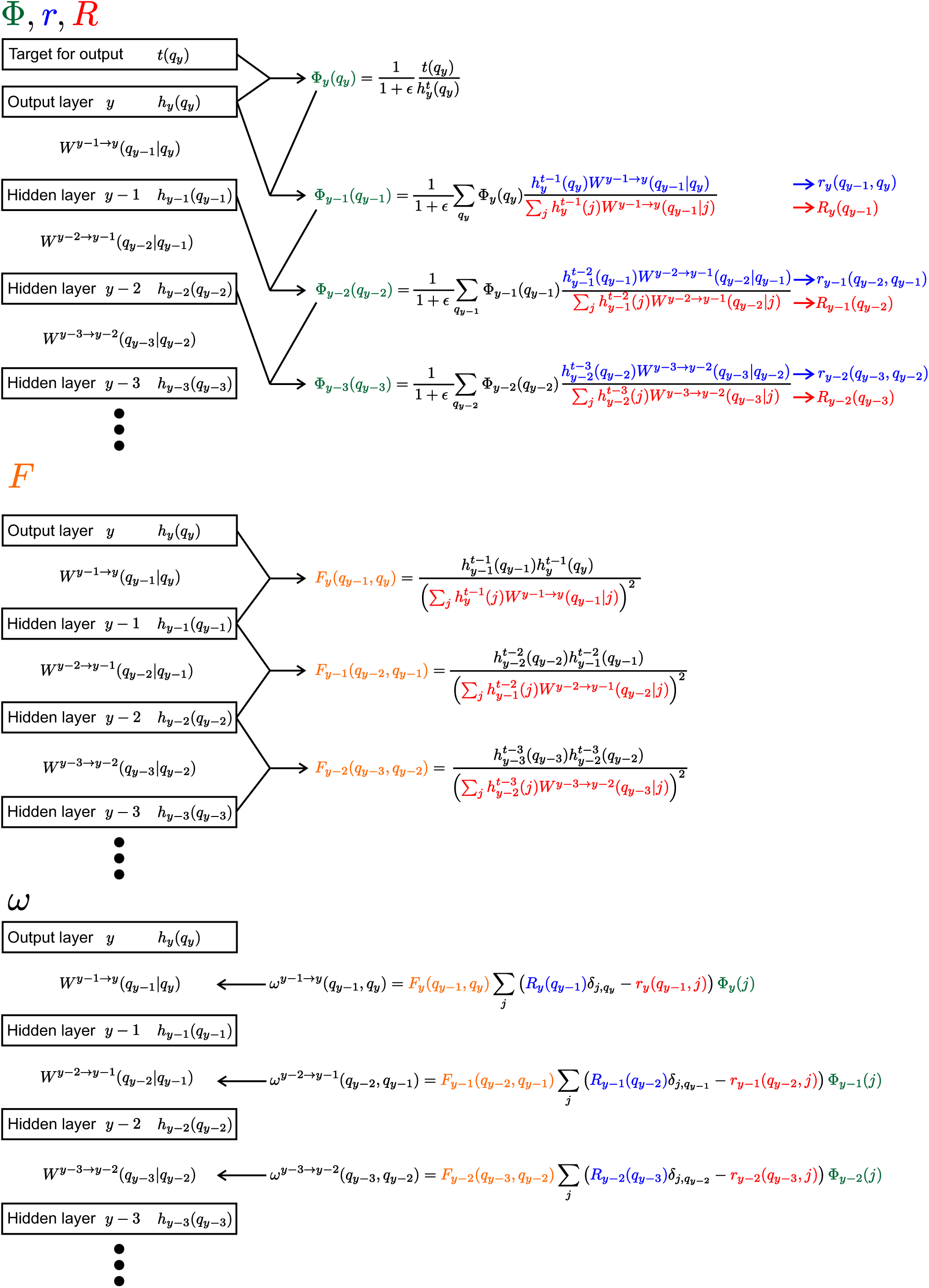
Visualization of the learning rule and its parts. The equation 8 and its sub-equations are shown for the output layer and three hidden layers. The pattern index *µ* isn’t shown to keep it uncluttered. Everything, except the weights *W*, has the same index *µ*.

The goal is to update the old weights *W* to get new weights *W*^*new*^, using the gradient 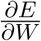 on the objective function *E*. This update for the weights between layer *y* − *m* − 1 and layer *y* − *m* (the output layer *y* is used as reference from which all the other layers are counted backwards) can be calculated via

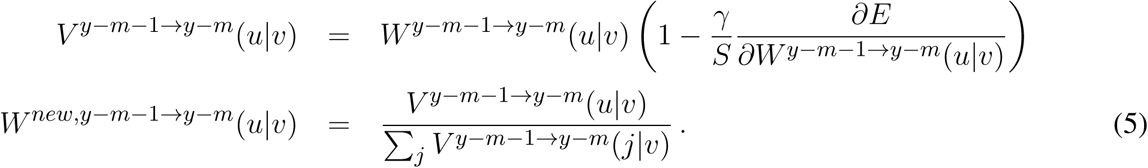

*γ* is a learning rate (0 < *γ* < 1) and *S* is a scaling factor for ensuring the non-negativity of the weight values. *S* needs to ensure that 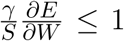 for all components of the weight matrix. This can be done by using

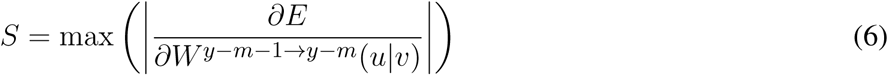

with calculating the maximum over all components of 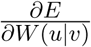.

The gradient itself is a sum over all contributions from a set of several pattern *µ* (i.e. a batch or mini-batch of input pattern) and, in the case of a convolutional layer, a sum over different spatial positions:

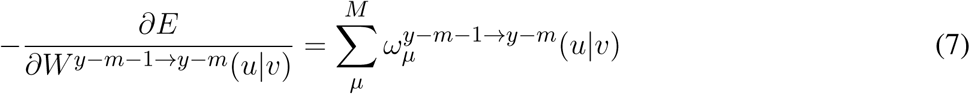

Each of the contribution *ω* is calculated from four sub-equations Φ, *r, R* and *F* (see figure 2):

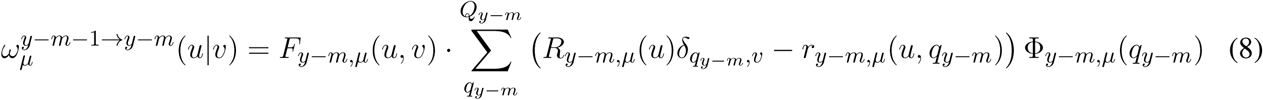

Φ is akin to the back propagating error in non-spiking neural networks. At the output layer *y*, the distribution over the latent variables 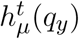 is compared to the desired target *t*_*µ*_(*p*_*y*_):

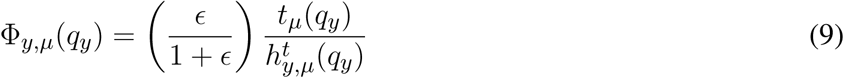

From there Φ is propagated backward in direction on the input (with *m* ≥ 1 as the distance measured to the output layer *y*):

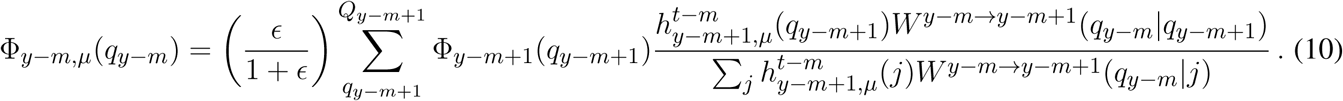

However, unlike in the typical back-prop rule Φ is not only propagating from the output layer to the input layer, it also travels back in time. This is due to the fact that Φ is updated with latent variables from further in the past the further one gets from the output layer.

For the other three components *r, R*, and *F* the equations are:

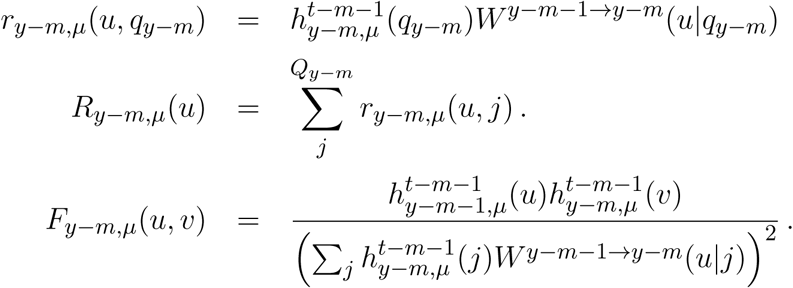

### 2.2 Learning the XOR function

The first example is a two layer network (see figure 3a) with four neurons in the input layer, four neurons in the hidden layer and two neurons in the output layer. The task of the network is to realize the XOR function, which receives two bits of input and outputs zero if the values of both bits are the same or one if both bits have different values. The input layer consists of two bits while every bit is represented by a population of two neurons. The first neuron in a population is only active if the input bit has a value of zero. The second input neuron of that population is only active if the input bit has a value of one. In every time step one spike is drawn from the input pattern distribution which is represented by the input neurons and send to the hidden layer. This is done using a Bernoulli processes.

**Figure 3.**
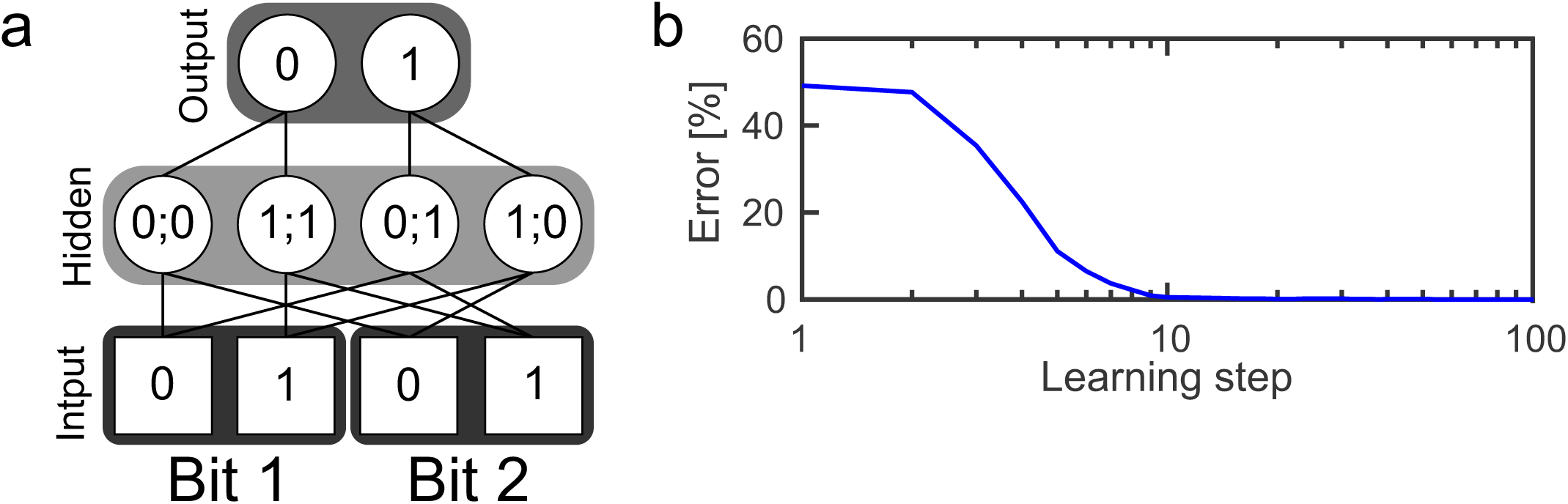
Learning the XOR function. a.) Structure of a SbS network for solving the XOR function. b.) Error of the network’s output during learning the weights, averaged over 250 initial conditions.

The hidden layer consists of one inference population (for which ∑_*i*_ *h*(*i*) = 1) with four neurons. In the figure, the non-zero weights between the input and the hidden layer are shown and have a value of 0.5 each. Given these weights and the SbS dynamics for *h*(*i*) (also called h-dynamic), after processing enough input spikes, only one hidden neuron will remain active due to the competition within an inference group. The corresponding input patterns, which lead to an activation of the neurons is listed in the hidden neurons in figure 3a.

The probability distribution formed by the latent variables in the hidden neurons is also used to draw one spike in every time step according a Bernoulli process. The spikes from the hidden layer are send to the output layer. The output layer processes incoming spikes according to the h-dynamic using the weight values between the hidden and output layer as shown in the figure. For decoding the result of the information processing, the output neuron with the higher value in its latent variable is selected. The first output neuron represents an output of zero and the second output neuron an output of one.

For the first test, the weights in this network were randomly initialized according to

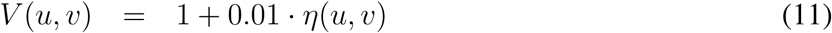

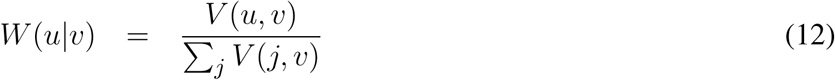

with *η*(*u, v*) as random numbers drawn from a uniform distribution [0, 1]. Before presentation of a new input pattern *p*_*X*_(*i*), the latent variables of the hidden neurons and the output neurons are always set to 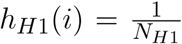 and 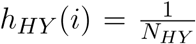, respectively. Then for every time step of the simulation, one spike each (i.e. the index of the neuron which fires next) is drawn from *p*_*X*_ and *h*_*H*1_ each. This is done by using these two probability distribution for a Bernoulli process.

These spikes are then used to update *h*_*H*1_(*i*) and *h*_*HY*_ (*i*) (using *ϵ* = 0.1 in the update process). The cycle of drawing spikes and updating the latent variables of the hidden and output layer is performed with the same input pattern until a given number of spikes has been processed. Using equation 8 the gradient for this pattern is calculated and stored. After collecting these contributions for all four input patterns (equation 7), equation 5 is applied to update the weights.

However, before normalizing *V* ^*y*−*m*−1→*y*−*m*^(*u*|*v*), it is ensured that the smallest value in *V* is Θ = 0.0001:

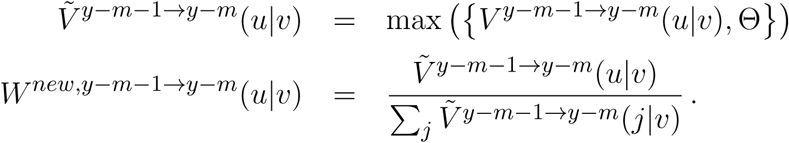

This prevents the multiplicative update equation 5 from getting stuck in zero values. Figure 3b shows the quality of the output during learning, averaged over 250 initial weights. Using 1024 spikes per input pattern & learning step as well as *γ* = 0.025, an error of 0 is reached after 32 learning steps.

Analyzing the magnitude of the learned weight values, reveals that the weights have only changed a small amount from the randomly initialized weights. This is a result of the competition inside of the hidden as well as the output layer. Already small asymmetries can be used to solve the task correctly. In the supplemental materials we show the weights and present a procedure how to force the learning rule such that it produces weights more like the ideal weights shown in figure 3a.

Furthermore, we show in the supplemental materials that the learning rule is able to ignore non task relevant inputs that are not correlated to the goal of the training procedure.

### 2.3 Learning the 4 bit parity function

The 4 bit parity function can be understood as an extension of the 2 bit XOR function. This function counts the number of its input bits with value one. Then it outputs one if the count is odd or zero if the count is even. A SbS network able to realize this function has four layers (see figure 4a): Input layer *X* with 8 neurons, which encodes the input in a similar fashion to the XOR network but via four groups with two neurons each. Hidden layer *H*1 with also 8 neurons and hidden layer *H*2 with four neurons as well as the output layer with two neurons. Besides the network structure, the procedure for simulating this network is as described in the example for the XOR network. Again, learning started with randomly initialized weights (see XOR example for how the weights were initialized). 250 simulations with different initial seeds were performed.

**Figure 4.**
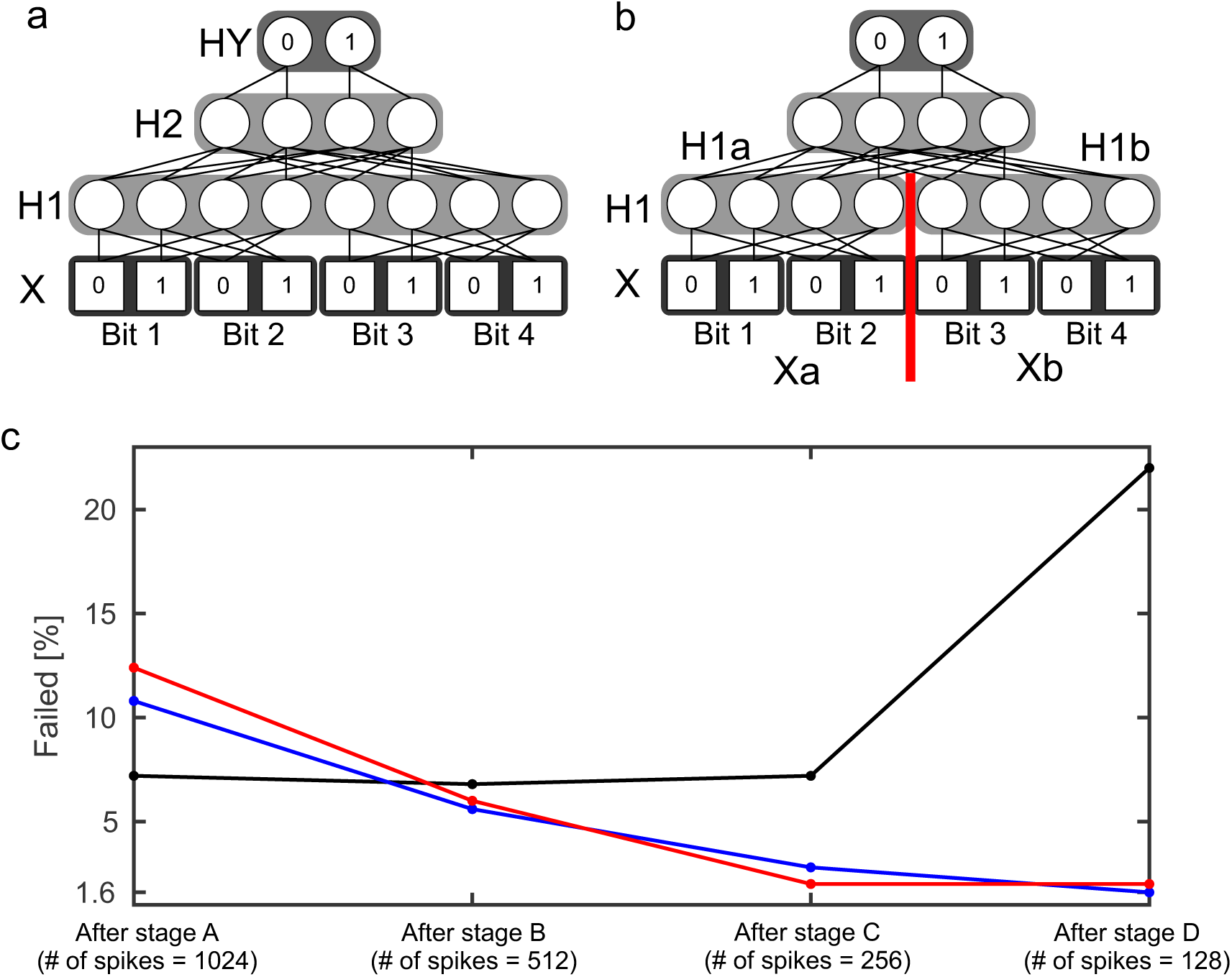
4 bit parity function. a.) Structure of a SbS network able to solve the 4 bit parity task. b.) Variation of the structure of the network. The hidden layer *H*1 was split into two normalization groups *H*1*a* and *H*1*b* with 4 neurons each. c.) Failure rate of learning the 4 bit parity function with the network structure shown in a.) (black) and using the network structure b.) (blue and red). For the red curve, the temporal back-propagation of the latent variables was replaced by the last value. The failure rate is shown for the weights after the four stages of learning.

In the following, if two performances are called significantly different then an one-sided Fisher’s exact test with p-level of 1% was used to determine this statement.

In summary, learning the 4 bit parity function shows that the learning algorithm has a problem with local minima.

We tried to minimize the amount of failed attempts (i.e. only if all the possible outputs of the network are correct then it is not a failed attempt) by using different combinations of numbers of spikes per pattern and *γ* learning rates during learning. Such a procedure showed fruitful results in the XOR example where it lead to weights that looked more like the ideal weights (see supplemental material for the details). Thus we also applied it to the 4 bit parity function.

Learning went through four stages with every stage had 7000 learning steps each. Furthermore, the subsequent stages started with the final weights from the stage before, while the first stage started with random weights. For the stage A, 1024 spikes per pattern were used. The following stages reduced the amount of spikes by a factor of 2 (512 spikes for stage B, 256 spikes for stage C, and 128 spikes for stage D) for learning. However, the performance was tested with 1024 spikes per pattern, to keep the results comparable.

In addition to reducing the number of spikes from one stage to the next, the learning rate gamma is reduced during a stage. Every stage starts with *γ* = 0.03. After every 1000 learning steps, *γ* is divided by 2 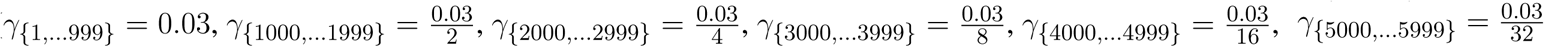, and 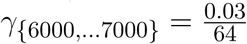. The idea behind this procedure is that in the beginning are allowed to change strongly and then they are supposed to settle at their correct values. In the supplemental materials, detailed learning curves for the four stages are shown. The black line in figure 4c shows the amount of initial conditions that failed to learn after the stages. Figure 4c reveals that the first three stages don’t improve the failure rate. Furthermore, stage D even significantly increases the number of failed learning attempts.

A putative source for this problem could be that for a successfully operating network two neurons in *H*1 need to be active simultaneously. However, learning a SbS IP tends to favor sparse solutions. Thus, the failed learning attempts could be a result of over-sparsification. Hence, we modified the network structure (see figure 4b) and split the hidden layer *H*1 into two normalization groups *H*1*a* and *H*1*b*. Now a successful network needs only one active neuron per SbS IP. *H*1*a* gets only input from the first two bits (*Xa*) of the Input *X* and *H*1*b* sees the spikes from the latter two input bits (*Xb*).

With the new structure five spikes are drawn in every time step: One spike each from *Xa, Xb, H*1*a, H*1*b* and *H*2. The weights between *Xa* and *H*1*a* as well as the weights between *Xb* and *H*1*b* are the same, like it would be in a convolutional neuronal network. During learning, the two contributions from the SbS backprop learning rule are averaged. Figure 4c (blue line) shows that there is now a significant difference between the failure rate after stage A and after stage D. Also a significant difference of the failure rate after stage D between the network with the split *H*1 layer (1.6%) vs the original network (22.0%) was found. Splitting layer *H*1 into two IPs reduces the failure rate to.

As final test with the 4 bit parity function, we used the network with the split in *H*1 and investigated how important the retardation of the latent variables during learning is. Instead of using the h-values from the earlier spikes – like it is required by the derived SbS backprop rule –, we only used the h-values after processing the last spike for every input pattern. Figure 4c (red line) shows here no significant difference. Thus it might be an interesting alternative (e.g. for saving memory) to neglect the retardation on the latent variables.

Examples for successfully working weights sets for these three tests are shown in the supplemental materials.

### 2.4 Deep convolutional network (MNIST)

The MNIST database is a benchmark for machine learning (see http://yann.lecun.com/exdb/mnist/). It consists of handwritten digits with 28 × 28 pixels and 256 gray values per pixel. The database contains 60,000 examples for training the network and 10,000 examples for testing the performance of the network. As part of the tutorial for the Google TensorFlow machine learning software, a convolutional neuron network for classifying these handwritten digits is presented https://www.tensorflow.org/tutorials/estimators/cnn. In their example they present a simple network learned via back-propagation based on a simple gradient descent optimization method. The performance for this network is listed with 97.3% classifications correct. Replacing the simple gradient decent by Adam (Kingma and Ba, 2014) and dropout learning (Srivastava et al., 2014), allows this network to reach a performance of 99.2% classifications correct (both source codes can be found in the supplemental materials).

The network in the TensorFlow tutorial is structured as follows: The input layer *X* consists of 28 × 28 elements representing the picture of a handwritten digit. The input layer is followed by a first hidden layer *H*1 which performs a convolution (with stride 1 and zero padding for keeping the size at 28 × 28 pixel after the convolution) through a kernel with the size of 5 × 5 with 32 filters. *H*1 is followed by a max pooling layer *H*2, which calculated the maximum over a 2 × 2 segment with stride of 2 from *H*1’s output. Layer *H*3 is again a convolution layer like *H*1 but looking at the output of *H*2 and with 64 filters instead. *H*4 is a 2 × 2 max pooling layer to *H*3. After *H*4, a fully connected layer *H*5 with 1024 neurons is positioned. And finally the output of the network can be read out from the output layer *HY*. *HY* consists of 10 neurons, where each neuron represents one class of the 10 digits. The classification result is decoded by calculating the argmax from *HY*’s neurons.

This network structure is mimicked by a SbS network. Since the computational complexity of the SbS network is orders of magnitude bigger, we simplified the TensorFlow example network. We removed the zero padding in both convolutional layers of the network. Thus layer *X* still has 28 × 28 pixels but layer *H*1 decreases to 24 × 24 with 32 filters. *H*2 halves the size to 12 × 12. Convolution layer *H*3 compressed the output of *H*2 to 8 × 8 with 64 filters. Its max pooling layer halves it again to 4 × 4. We keep the 1024 neurons for the fully connected layer *H*5 as well as the 10 neurons for the output layer. In the supplemental materials, the structure of the SbS network is discussed in detail, especially that all layers – including the pooling layer – realize the same algorithm.

We used TensorFlow to train this reduced network with a simple gradient descent optimization and got a classification correct performance of 97.1%. Furthermore, we got still 99.2% for a version using Adam and dropout learning. In Rotermund and Pawelzik (2019b) we showed that – with 97.8% classification correct – this SbS network structure using a only local learning rule (based on simple gradient descent and bi-directional flow of spikes) can match the performance TensorFlow example with the simple gradient descent optimization.

In the following we will present the SbS back-prop learning rule combined with a new optimizer applied to the MNIST benchmark.

For the input to the SbS network, a so called on/off split was made (Ernst et al. (2007)) which results in two channels per pixel. This is very similar to the representation of a bit by two neurons in the XOR example. This transformation is defined by *I*_*ON*_ (*x, y*) = *f* (2*P* (*x, y*) − 1) and *I*_*Off*_ (*x, y*) = *f* (1 − 2*P* (*x, y*)) with *f* (·) as a threshold linear function which sets all negative values to 0 and passes on all positive values without change.

Furthermore, instead of using max functions, the pooling layers *H*2 & *H*4 use only the inherent competition implemented by the SbS update rule. For the pooling IPs, the weights are fixed and not learned. E.g. a 2 × 2 spatial patch from convolution layer *H*1 (with its 2 × 2 × 32 neurons) delivers input to one pooling IP in *H*2 which has also 32 neurons. The structure of the weights ensures that the input from different features (i.e. the 32 features that are represented by the 32 neurons in the IPs) do not mix. Thus the combined spatial inputs from the 32 features compete against each other. Only the features with strong inputs are represented in the corresponding *H*2 normalization group. The same happens for pooling layer *H*4 but with 64 neurons per IP.

A widely known experience from learning neural networks is that using a simple gradient descent is prone to end in a low performance. Thus optimizers like Adam (Kingma and Ba, 2014) are used. Transferring optimizers like Adam to the SbS world is problematic due to the normalization and non-negativity boundary conditions on the latent variables and the weights. As alternative we developed an optimizer for SbS networks inspired by L4 (Rolinek and Martius, 2018). First of all, we used random mini-batches with 10% of the whole training data set. We smoothed the gradients from our SbS back-prop learning rule as well as the Kullback-Leibler divergence *KL* (measured between the desired and the actual distribution of the latent variables of the output layer *HY*) over the mini-batch according the equation from (Rolinek and Martius, 2018):

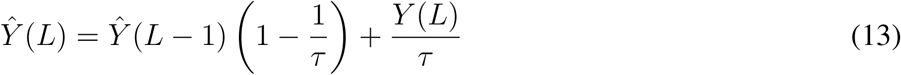

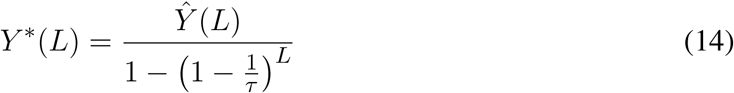

where *Y* (*L*) is the entity which needs smoothing and with *L* as learning step (i.e. number of used mini-batches) (*L* ∈ {1, … *L*_*Max*_}). Since we use mini-batches with 10% of the pattern randomly selected from the whole training data set, we selected *τ* = 10. In the following we will denote low-pass filtered variables with a ∗.

Furthermore, we modulated the learning rate *γ* with the smoothed Kullback-Leibler divergence *KL*^∗^ via

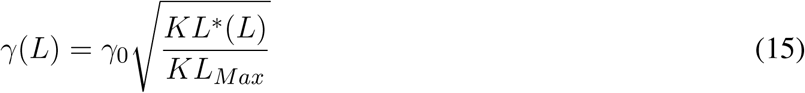

with *γ*_0_ = 0.05 (which was our guess for a good initial learning rate),

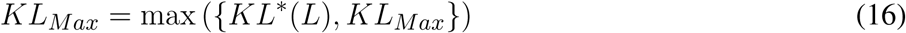

and

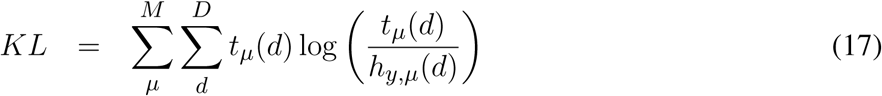

where *M* is the size of the mini-batch. In equation 15 we selected to use the square root, compared to a linear function in (Rolinek and Martius, 2018). Our reasoning was that this keeps the learning rate higher in the beginning. We didn’t compared the performances that both choices would result in.

The updates of the weights are done according to

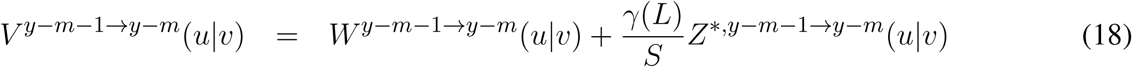

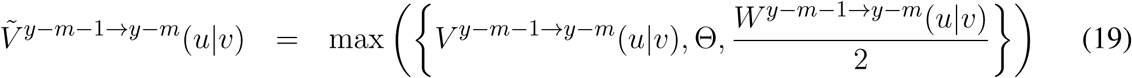

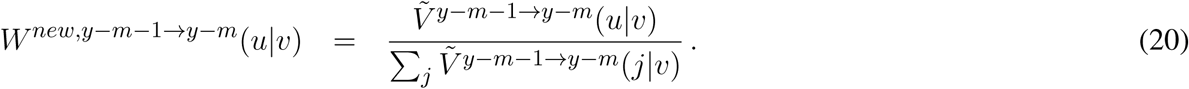

using Θ = 0.0001 and

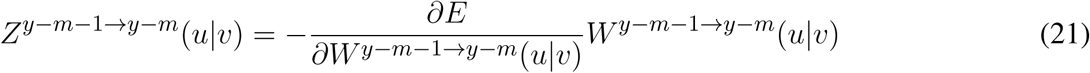

as well as

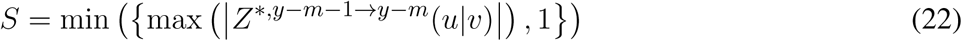

with calculating the maximum over all components of *Z*^∗^(*u*|*v*).

For the SbS network with the back-prop learning rule, a three stage learning procedure is applied. This sequence of stages is shown in figure 6a. In figure 6b the development of the classification error over the these stages is summarized. In the supplemental materials detailed plots are shown that present how the training error (measured with the Kullback-Leibler divergence), learning rate, and the classification error on the test data set develops.

**Figure 5.**
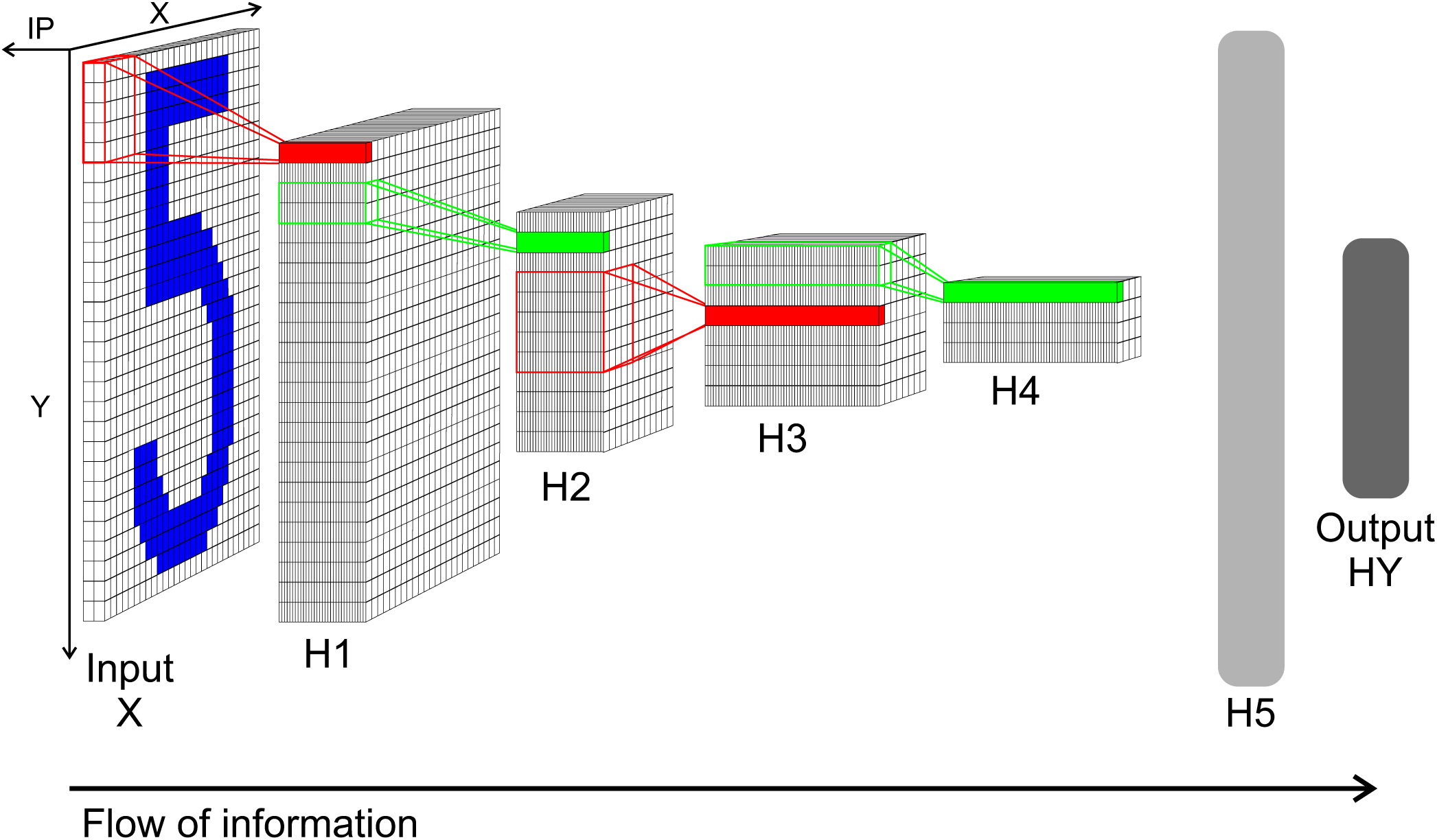
Network structure of the convolution network for the MNIST data. Input *X*: Input layer with 28 × 28 normalization modules for 28 × 28 input pixel. Each module has two neurons realizing a simplified version of on/off cells for enforcing positive activity also for low pixel values. From this layer spikes are send to layer *H*1. *H*1: Convolution layer *H*1 with 24 × 24 IPs with 32 neurons each. Every IP processes the spikes from 5 × 5 spatial patches of the input pattern (x and y stride is 1). *H*2: 2 × 2 pooling layer *H*2 (x and y stride is 2) with 12 × 12 IPs with 32 neurons each. The weights between *H*1 and *H*2 are not learned but set to a fixed weight matrix that creates a competition between the 32 features of *H*1. *H*3: 5 × 5 convolution layer *H*3 (x and y stride is 1) with 8 × 8 IPs. Similar to *H*1 but with 64 neuron for each IP. *H*4: 2 × 2 pooling layer *H*4 (x and y stride is 2) with 4 × 4 IPs with 64 neurons each. This layer is similar to layer *H*2. *H*5: Fully connected layer *H*5. E.g. 1024 neurons in one big IP which are fully connected to layer *H*4 and output layer *HY*. *HY* : Output layer *HY* with 10 neurons for the 10 types of digits. For decoding the identity of the neuron with the highest activity is selected.

**Figure 6.**
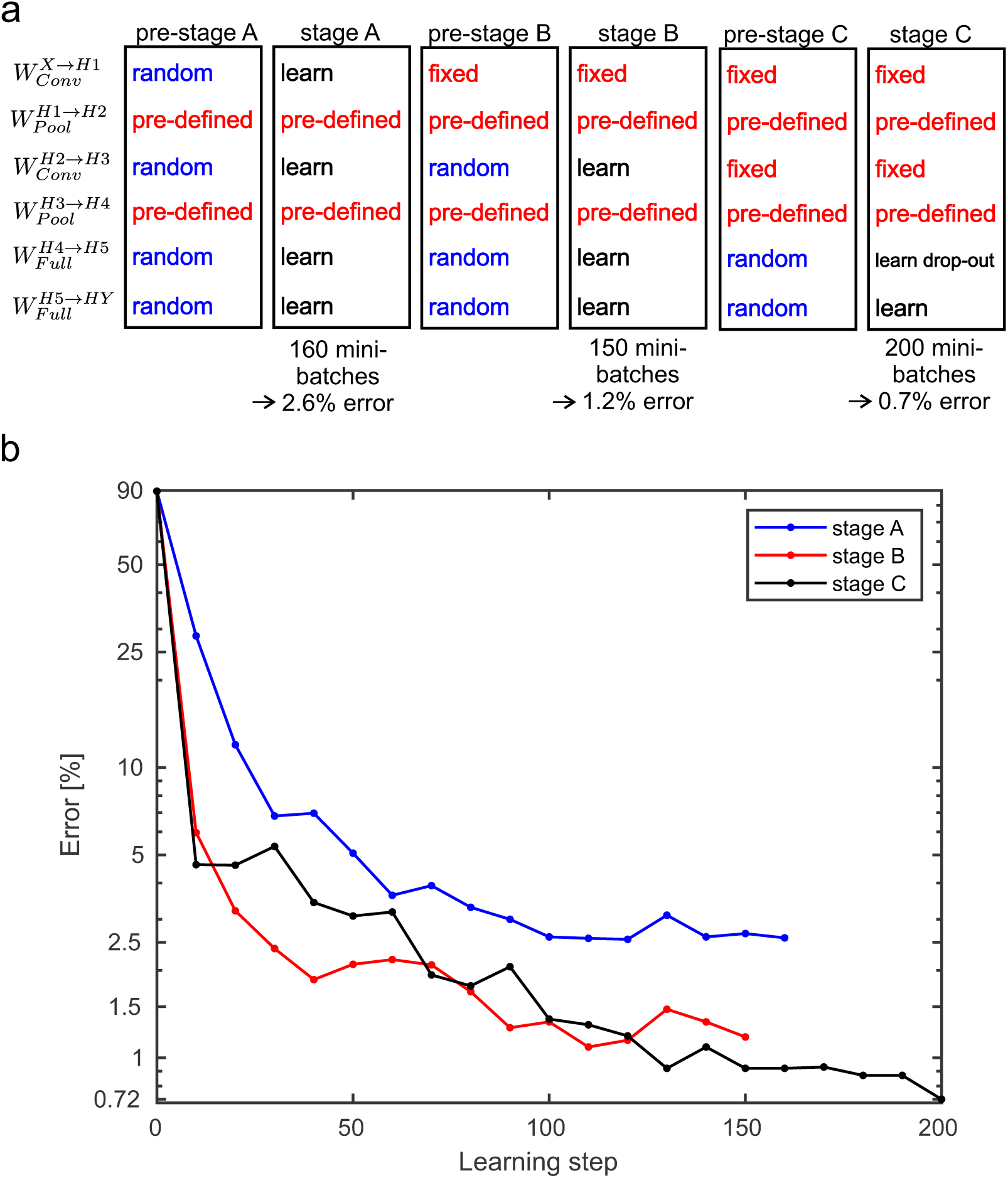
Performance values for the MNIST benchmark. a.) A three stage learning process was used. First, in pre-stage A all weights are set to random values except the weights for the pooling layers which are pre-defined and not learned at all during any of the stage. In stage A the weights are trained for 160 mini-batches. This results in a classification error on the test data of 2.6%. Then in pre-stage B, the weights for the second convolutinal layer and the fully connected layer are reset to random values. In stage B these random weights are trained again for 150 mini-batches which yields an error of 1.2%. In pre-stage C, the weights for the fully connected layers are replaced by random weights and in stage C these weights are learned again. However, this time a drop-out procedure for the layer *H*5 is applied. After processing 200 mini-batches an error of 0.72% is reached. Alternatively, stage C can be done without drop-out and then yields in an error of 0.83% (see supplemental materials for the learning curves). b.) Development of the error on the MNIST test data set for the three stages over the performed learning steps.

During learning weights are set to random values, this is done by

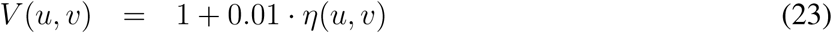

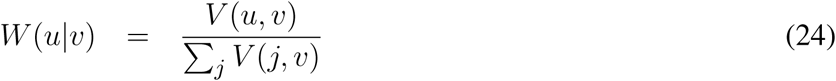

with *η*(*u, v*) as random numbers drawn from a uniform distribution [0, 1].

For every input pattern, every input or SbS inference population generates 1200 spikes over the course of simulating the SbS network for this pattern.

For the first stage of learning, we selected *ϵ*_0_ = 0.1 which is our standard value that typically works. For the latter two stages we reduced it to 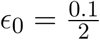. The reasoning behind this decision was that we feared that otherwise the higher layer could might get too sparse in the distribution of latent variables in the SbS IPs. From *ϵ*_0_ we derive *ϵ* values for the SbS IPs in the different layers. The underlying idea is that while IPs in different layers get a different amount of spikes in every time step of the simulations, we wanted to equalize the change on the latent variables of all IPs. Thus we divided *ϵ*_0_ by the amount of spikes an IP receives in one time step. This results in: 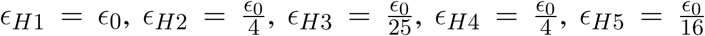, and *ϵ*_*HY*_ = *ϵ*_0_. After 1000 spikes *ϵ*_0_ is divided by 25. Since *ϵ* acts like a low-pass filter parameter that is controlling of the impact one spike can have on the latent variables, reducing *ϵ* results in a reduction on the fluctuations of the latent variables. First, we want to allow the network to reach some ‘good’ state with the first 1000 spikes, then we use the reduction on *ϵ* and the next 200 spikes to implement an implicit averaging of the latent variables over the incoming spikes.

It is important to note, that we didn’t optimize these *ϵ* values or *ϵ*_0_ due to missing computational power. The parameters stem from a mere guess which parameters might work. The same *ϵ* values were used for learning the weights and testing the classification performance.

The first stage of learning starts with all the weights set to random values, except the pooling layer which are pre-set and not changed or learned at all. Learning the weights for 160 mini-batches, we reached a classification error on the test data set of 2.6%. Then the weights *W*^*H*2→*H*3^, *W*^*H*4→*H*5^, and *W*^*H*5→*HY*^ are set to random values again. Only the weights *W*^*X*→*H*1^ are kept and not learned during stage B. After 150 mini-batches the error goes down to 1.2%. Then again, the weights *W*^*H*4→*H*5^ and *W*^*H*5→*HY*^ are set to random values again. *W*^*X*→*H*1^ and *W*^*H*2→*H*3^ are kept constant.

When learning is performed again, the error goes down to 0.83% (see supplemental materials for detailed learning curves). However, sparseness on the latent variables can get a problem in the layer *H*5 with its 1024 neurons. Selected neurons can show very strong values in their latent variables which results – due to the competition in the SbS IP – in suppressing the other neurons. During learning this can get a self-reinforcing process. For compensation this behavior, we devised a strategy inspired from dropout learning (Srivastava et al., 2014).

Normally, in the beginning of the simulation all the latent variables start with the value 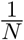, were *N* is the number of neurons in a SbS IP. A simple way to include dropout in the SbS model is to initialize selected neuron’s latent variables with the value zero. Since the update rule for the latent variables is multiplicative, such a neuron is disabled for that simulation. We use this aspect of the h-dynamic to implement dropout for layer *H*5 and to control how any neurons in *H*5 are active at the same time during learning.

In the beginning of the learning process we want only a few neurons to be active in *H*5 during learning (We started with 16 neurons of 1024). Thus we calculated for every input pattern 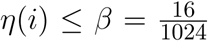 with *η*(*i*) as random numbers drawn from a uniform distribution [0, 1]. For every *H*5 neuron *i* that fulfilled this condition, its latent variable 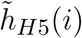 was set to one. Otherwise the latent variable 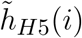 was set to zero. Afterwards, the latent variables of the IPs are normalized 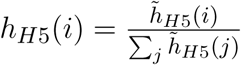 and used for the simulation as initial values. After every 30 mini-batches the number of active neurons is double (i.e. *β* is doubled), until *β* reaches a value of one.

Applying dropout learning over a duration of 200 mini-batches yields an classification error of 0.72% (see supplemental materials for detailed learning curves).

For the 10% mini-batches used in our simulations, we didn’t noticed any over-fitting. Thus the Kullback-Leibler divergence on the training data was a reliable estimator for the expected classification performance on the test data. However, it needs to be noted that this might not the case if the mini-batches get bigger. Then it might be necessary to separate a part of the training data as validation data set.

## 3 DISCUSSION

The present work is based on a framework where the basic computational units are local populations rather than individual neurons. These so called inference populations iteratively perform inference on the potential causes of their inputs from each stochastic spike impinging on the population (Ernst et al., 2007). While this dynamics might appear artificial, it captures the essence of models with more biologically realistic neurons (Rozell et al., 2008; Zhu and Rozell, 2015; Moreno-Bote and Drugowitsch, 2015) that perform sparse efficient coding, which is a leading hypothesis for understanding coding in the brain (Olshausen and Field, 2006; Spanne and Jörntell, 2015; Zhu and Rozell, 2015; Capparelli et al., 2019).

Hierarchical generative networks built from inference modules are used in technical approaches as e.g. for image generation (Ghosh et al., 2019). There are many papers concerned with learning such deep generative models with error back-propagation (e.g. Lee and Seung (1999, 2001); Oh and Seung (1998); Guo and Zhang (2017); Rezende et al. (2014); Bengio et al. (2014); Salakhutdinov (2015)) as well as publications on training similar networks that use non-negative matrix factorization (e.g. Ahn et al. (2004); Zeng et al. (2016)). In contrast to these approaches, the present framework conserves the specific spike driven update dynamics from Ernst et al. (2007) which is of special interest because it results in a very simple yet biologically plausible neuronal network that uses only spikes as signals. It allows to build special computational hardware that can be massively parallelized (Rotermund and Pawelzik, 2018) and exhibits sparse representations that are know from compressed sensing (Ganguli and Sompolinsky, 2012, 2010).

While deep convolutional networks were successful in predicting responses of neurons in primate visual cortex investigations into the potential of deep generative models for explaining natural computation in the brain are just beginning (Bengio et al., 2014, 2015). But even if formal approaches along these lines were phenomenologically successful, realistic models are still required to elucidate how the striking performances of brains in terms of speed and precision are realized with the stochastic spikes that in reality are available as sole signals.

For exploring the potential of SbS networks as alternative to deep convolutional networks with respect to performance in technical applications as well as as models for real neuronal networks suitable learning rules are missing. As a first step towards this goal we here present an error back-propagation based learning rule for training multi-layer feed-forward SbS networks.

Having a supervised method for optimizing the weights in deep SbS networks allows not only to explore the potential of the rather unusual SbS framework but also to compare it to other approaches towards learning in these networks as e.g. a local rule for unsupervised learning in recurrent SbS networks (Rotermund and Pawelzik, 2019b).

Using simple examples with Boolean functions we show that the backpropagation algorithm presented here can learn the weights in a given forward network architecture from scratch (i.e. randomly initialized weights) as well as to ignore non task relevant information. We found that the algorithm can be used to train weights several layers away from the output. We also proposed architectures with convolutional and pooling layers composed of the same basic SbS elements. In particular, no special functionality was required for setting up the pooling layers, in contrast to usual deep convolutional networks.

We used the MNIST benchmark to investigate deeper convolutional SbS networks. Beside the input and the output layer, the network has five internal layers where two are pooling layers and two convolutional layers as well as one fully connected layer. Overall, the network contains 1378 populations with 57994 neurons. During the simulation of one input pattern, a total of 1200·1378 spikes are generated (28.5 spikes per neuron). On the MNIST benchmark test data, our SbS network achieved up to 99.3% classification correct performance. Using the same the architecture for a non-spiking convolutional neural network we measured 99.2%. However, comparing performance values with other networks (e.g. see (Tavanaei et al., 2018) and for a list of MNIST networks http://yann.lecun.com/exdb/mnist) is not as simple as comparing the performance values. In our case we didn’t optimized the network structure for the use of SbS inference populations. We rather decided to re-use the network structure associated with the Tensor Flow Tutorial because this gave us a base-line for a network design which our computer cluster was just been able to simulate. Furthermore, we didn’t use any input distortion methods (e.g. shifting, scaling, or rotating the input pictures) for increasing the size of the training data set, methods that are known to lead to performance values of up to 99.8% (Wan et al., 2013). The reason was simply that this would have been too much for our computer cluster, like it would have been to optimize the parameters used in the SbS MNIST network. Or in other words: The performances shown for the MNIST SbS network certainly do not reflect what a fully optimized SbS network might be capable to deliver.

The SbS-approach avoids the real time dynamics required for simulation of noisy leaky integrate-and-fire (IaF) neurons where the membrane potential needs to become updated every time step *dt* which is often in the range of sub-milliseconds. The number of updates between two spikes depends on the firing rate of that neuron. If for example *dt* = 0.1*ms* and the firing rate is 10Hz then this would translate roughly into 1000 updates of the membrane potential between two spikes. Compared to that, a SbS neuron would perform one update. While the different types of spiking neuron models (Izhikevich, 2004) have varying number of computations for one update, in a SbS population with *N* neurons 3*N* multiplications, 2*N* summations, and one division are used for one update of the whole population. This reduction in computational requirement is payed by a decrease in biological realism.

An approach akin to the SbS’s removal of real time is also known for integrate-and-fire neurons. This so called event-based neuronal networks (e.g. Brette (2006, 2007); Serrano-Gotarredona et al. (2015); Lagorce et al. (2015)) use analytic solutions of the neuron’s dynamics to bridge the time between to spikes. However, with stochastic neurons this approach gets problematic (Brette, 2007). This is similar to the problem of finding an analytically solution for the first passage time (Burkitt, 2006a,b) of neurons in a network of neurons with stochastic inputs, which is a hard problem too. In the case of the SbS network, the stochasticity is an important aspect because it is a type of importance sampling of the input as well as the latent variables. This acts as a filter for the capturing the more dominant information in the network and suppress noise.

Concerning the parallelization of non-spiking neural networks compared to the SbS model: A traditional deep convolutional neuronal network (DNN) implements several different types of layers (e.g. convolutional layer, pooling layers, and dense layers) for which it requires a variety of optimized hardware elements (Sze et al., 2017). In the case of the SbS model, all these different type of layers are represented by the same dynamics of the latent variables. A hardware solution for SbS networks (Rotermund and Pawelzik, 2018) can be understood more like a pool of SbS inference populations that are shaped into the desired network structure by just organizing the flow of the spikes. Furthermore, it is less easy to extend one layer of a non-spiking network over several ASICs. This is a consequence from the high amount of information that needs to be exchanged in a typical non-spiking DNN. In the SbS case, the computation is already compartmentalized into IPs that can be operatied also asynchronously having a low bandwidth communication among each other.

In summary, we presented an error back-propagation based learning rule for training multi-layer feed-foreward SbS networks. This significantly extends earlier work (Ernst et al. (2007)) such that for the first time it makes supervised training of deep Spike-By-Spike based networks possible. Combined with optimized and massively parallel computational hardware (Rotermund and Pawelzik (2018)) this will open the door for future investigations of this conceptually simple spike based neuronal network which we believe has interesting properties as a generative model with in-built sparseness for both, technical applications as well as as model for natural computations in real brains.

## Supporting information

Supplemental information and pictures

## ACKNOWLEDGMENTS

This manuscript has been released as a Pre-Print at www.biorxiv.org (Rotermund and Pawelzik (2019a)).

