## Supplementary material for "Back-propagation learning in deep Spike-By-Spike networks": Source Code: Calc_ExactFischerTest.m

```
% It is assumes that N is the same for both tests (N = number of total samples i.e. trials)
% a_I and b_I are the number of correct training results 
% P is the probability that both a_I and b_I are drawn form the same distribution
function [P] = Calc_ExactFischerTest(a_I,b_I,N)

a = min([a_I b_I]);
b = max([a_I b_I]);

[Dummy P] = fishertest([[a;N-a],[b;N-b]],'Tail','left');
```
