## Supplementary material for "Back-propagation learning in deep Spike-By-Spike networks": Source Code: DiffFactor.m

```
function [DiffFa] = DiffFactor(Factor, ReconstructionMatrix, SumReconstruction, FF)

DiffFa = FF .* (repmat(SumReconstruction,[1 size(ReconstructionMatrix,2)]) .* repmat(Factor(:)',[size(ReconstructionMatrix,1) 1])  - repmat(ReconstructionMatrix * Factor,[1 size(ReconstructionMatrix,2)]));
```
