## Supplementary material for "Back-propagation learning in deep Spike-By-Spike networks": Source Code: FrontFactor.m

```
function [FF, ReconstructionMatrix, SumReconstruction] = FrontFactor(H_B, H_A, W, Epsilon)

FF = repmat(H_A(:)', [length(H_B(:))  1] ) .* repmat(H_B(:), [1 length(H_A)]);

ReconstructionMatrix = W .* repmat(H_A(:)', [size(W,1) 1] );

SumReconstruction = sum(ReconstructionMatrix,2);

Idx = find(FF == 0);
FF = (Epsilon / (1 + Epsilon)) * FF ./ repmat(SumReconstruction.^2,[1 length(H_A)]);
FF(Idx) = 0;
```
