## Supplementary material for "Back-propagation learning in deep Spike-By-Spike networks": Source Code: GiveMeASpike.m

```
function [Spike] = GiveMeASpike(Distribution)

  Distribution = Distribution(:);
  Distribution = Distribution / sum(Distribution);
  
  RandomNumber = rand;

  for i = 1:1:length(Distribution),

    RandomNumber = RandomNumber - Distribution(i);

    if (RandomNumber <= 0)
      Spike = i;
      return;
    end

  end

  Spike = length(Distribution);
```
