## Supplementary material for "Back-propagation learning in deep Spike-By-Spike networks": Source Code: plotit_2_OK.m

```
SeedID = 2;
eval(['load Data_C/Data_' num2str(SeedID) '.mat W_H1_H2 W_X_H1 W_H2_HY Correct_Full']);

Handle = figure(2);

subplot(3, 1, 1)
imagesc(W_X_H1)
colorbar
axis off
axis image


subplot(3, 1, 2)
imagesc(W_H1_H2)
colorbar
axis off
axis image


subplot(3, 1, 3)
imagesc(W_H2_HY)
colorbar
axis off
axis image


colormap('hot')
drawnow

print(Handle,'SubPlot_3_2_OK.pdf','-dpdf')
```
