## Supplementary material for "Back-propagation learning in deep Spike-By-Spike networks": Source Code: plotit_a.m

```
clear
First = true;
LStep_Max = 7000;
Smooth = [1:9 10:2:99 100:5:999 1000:10:LStep_Max];

for SeedID = 1:1:250
  try
    eval(['load Data_A/Data_' num2str(SeedID) '.mat KL W_Range Correct Correct_Full']);
    if First == true
      KL_Av = KL;
      W_Range_Av = W_Range;
      Correct_Av = Correct;
      Correct_Full_Av = Correct_Full;
      First = false;
      Counter = 1;
    else
      KL_Av = KL_Av + KL;
      W_Range_Av = W_Range_Av + W_Range;
      Correct_Av = Correct_Av + Correct;
      Correct_Full_Av = Correct_Full_Av + Correct_Full;
      Counter =  Counter + 1;
    end
  catch
  end
end

KL = KL_Av / Counter;
W_Range = W_Range_Av / Counter;
Correct = Correct_Av / (Counter*16);
Correct_Full = Correct_Full_Av / (Counter*16);

Handle = figure(10);

subplot(3, 1, 1)
hold off
loglog([1000 1000],[10^0  60],'k--')
hold on
for i = 1:1:6
  loglog([1000 1000]+i*1000,[10^0 60],'k--')
end
loglog(Smooth,(1.0-Correct)*100.0)
loglog(Smooth,(1.0-Correct_Full)*100.0)
axis tight
yticks([1 2.5 5 10 25 50])
ylim([0 60])
xlabel('Learning step')
ylabel('Error [%]')


subplot(3, 1, 2)
hold off
loglog([1000 1000],[10^-1  15],'k--')
hold on
for i = 1:1:6
  semilogx([1000 1000]+i*1000,[10^-1 15],'k--')
end
loglog(Smooth,KL);
axis tight
yticks([0.15 0.25 0.5 1 2.5 5 10])
ylim([0.15 13])
xlabel('Learning step')
ylabel('KL')

subplot(3, 1, 3)
hold off
semilogx([1000 1000],[0 0.775],'k--')
hold on
for i = 1:1:6
  semilogx([1000 1000]+i*1000,[0 0.775],'k--')
end
semilogx(Smooth,W_Range(:,1:2),'r');
semilogx(Smooth,W_Range(:,3:4),'b');
semilogx(Smooth,W_Range(:,5:6),'k');
axis tight
ylim([0 0.775])
yticks([0 0.125 0.25 0.375 0.5 0.625 0.75])

xlabel('Learning step')
ylabel('Weight range')
drawnow

print(Handle,'SubPlot_3_A.pdf','-dpdf')
```
