## Supplementary material for "Back-propagation learning in deep Spike-By-Spike networks": Source Code: README.txt

The simulation with the 250 different seeds are run as follows: 
for TheSeed = 1:1:250,
  stage_all(TheSeed);
end

The back-prop learning rule used the mini sub-functions DiffFactor.m, ErrorFactor.m, FrontFactor.m, and PropagateFactor.m.

Spikes are drawn from distributions by the function GiveMeASpike.m

With plotit_a.m, plotit_b.m, plotit_c.m, and plotit_d.m the results are plotted. 
