## Supplementary material for "Back-propagation learning in deep Spike-By-Spike networks": Source Code: stage_a.m

```
function paper(SeedID)

rand('seed',SeedID);

NumberIterations_0 = 1024;
PlotEveryX_0 = 50;

NumberIterations = NumberIterations_0;
PlotEveryX = PlotEveryX_0;

NumberOfBits = 4;
NumberOfPatterns = 2.^NumberOfBits;
NoN_X  = NumberOfBits * 2 ;
NoN_H1 = 4;
NoN_H2 = 4;
NoN_HY = 2;

Epsilon = 0.1;
Gamma = 0.03;

% Lernschritte:
LStep_Max = 7000;
WindowSize = 250;

Smooth = [1:9 10:2:99 100:5:999 1000:10:LStep_Max];

Done = false;

ExistTest = 0;
try
  eval(['load Data_A/Data_' num2str(SeedID) '.mat W_X_H1']);
catch
  ExistTest = 1;
end

if ExistTest == 0
  return;
end


%%%%%%%%%%%%%

W_X_H1 = ones(NoN_X/2,NoN_H1) + 1.0/100.0 * rand(NoN_X/2,NoN_H1);
W_X_H1 = W_X_H1 ./ repmat(sum(W_X_H1,1), [size(W_X_H1,1) 1]);

W_H1_H2 = ones(NoN_H1*2,NoN_H2) + 1.0/100.0 * rand(NoN_H1*2,NoN_H2);
W_H1_H2 = W_H1_H2 ./ repmat(sum(W_H1_H2,1), [size(W_H1_H2,1) 1]);

W_H2_HY = ones(NoN_H2,NoN_HY) + 1.0/100.0 * rand(NoN_H2,NoN_HY);
W_H2_HY = W_H2_HY ./ repmat(sum(W_H2_HY,1), [size(W_H2_HY,1) 1]);

Patterns_Input = zeros(NoN_X,NumberOfPatterns);
Patterns_Output = zeros(NoN_HY,NumberOfPatterns);

for i = 1:1:NumberOfPatterns,
    BinString = dec2bin(i-1,NumberOfBits);
    BinString = double(BinString)-48;
    Temp = zeros(NoN_X,1);
    Temp([1:2:NoN_X]+BinString) = 1;
    Patterns_Input(:,i) = Temp / sum(Temp);
end


Temp = mod(sum([Patterns_Input(1:2:end,:) > 0],1),2)+1;
for i = 1:1:NumberOfPatterns,
  Patterns_Output(Temp(i),i) = 1; 
end

OneDivOnePlusEps = 1 / (1 + Epsilon);

KL = zeros(length(Smooth),1);
W_Range = zeros(length(Smooth),6);
Correct = zeros(length(Smooth),1);
Correct_Full = zeros(length(Smooth),1);

for LStep = 1:1:LStep_Max,
  fprintf([num2str(LStep) ' of ' num2str(LStep_Max) ', Number of Spikes:' num2str(NumberIterations) ' \n']);

  Smooth_ID = find(Smooth == LStep);

  H1_A_Collect = zeros(NoN_H1,NumberOfPatterns,4);
  H1_B_Collect = zeros(NoN_H1,NumberOfPatterns,4);
  H2_Collect = zeros(NoN_H2,NumberOfPatterns,4);
  HY_Collect = zeros(NoN_HY,NumberOfPatterns,4);

  dW_X_H1 = zeros(size(W_X_H1));
  dW_H1_H2 = zeros(size(W_H1_H2));
  dW_H2_HY = zeros(size(W_H2_HY));

  if (LStep == 1000)
    Gamma = Gamma / 2.0;
  elseif (LStep == 2000)
    Gamma = Gamma / 2.0;
  elseif (LStep == 3000)
    Gamma = Gamma / 2.0;
  elseif (LStep == 4000)
    Gamma = Gamma / 2.0;
  elseif (LStep == 5000)
    Gamma = Gamma / 2.0;
  elseif (LStep == 6000)
    Gamma = Gamma / 2.0;
  end
 
  for PatternID = 1:1:NumberOfPatterns,
    H1_A = ones(NoN_H1,1) / NoN_H1;
    H1_B = ones(NoN_H1,1) / NoN_H1;
    H2 = ones(NoN_H2,1) / NoN_H2;
    HY = ones(NoN_HY,1) / NoN_HY;

    P_X = Patterns_Input(:,PatternID);
    P_Y = Patterns_Output(:,PatternID);

    for t = 1:1:NumberIterations,

      Spike_X_A     = GiveMeASpike(P_X(1:4)*2);
      Spike_X_B     = GiveMeASpike(P_X(5:8)*2);
      Spike_H1_A    = GiveMeASpike(H1_A);
      Spike_H1_B    = GiveMeASpike(H1_B);
      Spike_H2    = GiveMeASpike(H2);

      Temp = W_X_H1(Spike_X_A,:)' .* H1_A;
      ST = sum(Temp);
      if ST > 10^-100
        H1_A = OneDivOnePlusEps * (H1_A + Epsilon * Temp/ST);
      end

      Temp = W_X_H1(Spike_X_B,:)' .* H1_B;
      ST = sum(Temp);
      if ST > 10^-100
        H1_B = OneDivOnePlusEps * (H1_B + Epsilon * Temp/ST);
      end

      Temp = W_H1_H2(Spike_H1_A,:)' .* H2;
      ST = sum(Temp);
      if ST > 10^-100
        H2 = OneDivOnePlusEps * (H2 + Epsilon * Temp/ST);
      end

      Temp = W_H1_H2(Spike_H1_B+4,:)' .* H2;
      ST = sum(Temp);
      if ST > 10^-100
        H2 = OneDivOnePlusEps * (H2 + Epsilon * Temp/ST);
      end

      Temp = W_H2_HY(Spike_H2,:)' .* HY;
      ST = sum(Temp);
      if ST > 10^-100
        HY = OneDivOnePlusEps * (HY + Epsilon * Temp/ST);
      end

      if t > (NumberIterations-6 )
        H1_A_Collect(:,PatternID,4) = H1_A_Collect(:,PatternID,3);
        H1_A_Collect(:,PatternID,3) = H1_A_Collect(:,PatternID,2);
        H1_A_Collect(:,PatternID,2) = H1_A_Collect(:,PatternID,1);
        H1_A_Collect(:,PatternID,1) = H1_A;

        H1_B_Collect(:,PatternID,4) = H1_B_Collect(:,PatternID,3);
        H1_B_Collect(:,PatternID,3) = H1_B_Collect(:,PatternID,2);
        H1_B_Collect(:,PatternID,2) = H1_B_Collect(:,PatternID,1);
        H1_B_Collect(:,PatternID,1) = H1_B;

        H2_Collect(:,PatternID,4) = H2_Collect(:,PatternID,3);
        H2_Collect(:,PatternID,3) = H2_Collect(:,PatternID,2);
        H2_Collect(:,PatternID,2) = H2_Collect(:,PatternID,1);
        H2_Collect(:,PatternID,1) = H2;

        HY_Collect(:,PatternID,4) = HY_Collect(:,PatternID,3);
        HY_Collect(:,PatternID,3) = HY_Collect(:,PatternID,2);
        HY_Collect(:,PatternID,2) = HY_Collect(:,PatternID,1);
        HY_Collect(:,PatternID,1) = HY;
      end

    end

    if length(Smooth_ID) == 1
      [Dummy MaxPos_Correct] = max(P_Y);
      [Dummy MaxPos] = max(HY);

      Correct(Smooth_ID,1) = Correct(Smooth_ID,1) + double([MaxPos_Correct == MaxPos]);

      H1_A = ones(NoN_H1,1) / NoN_H1;
      H1_B = ones(NoN_H1,1) / NoN_H1;
      H2 = ones(NoN_H2,1) / NoN_H2;
      HY = ones(NoN_HY,1) / NoN_HY;


      for t = 1:1:1024,


        Spike_X_A     = GiveMeASpike(P_X(1:4)*2);
        Spike_X_B     = GiveMeASpike(P_X(5:8)*2);
        Spike_H1_A    = GiveMeASpike(H1_A);
        Spike_H1_B    = GiveMeASpike(H1_B);
        Spike_H2      = GiveMeASpike(H2);
      
        Temp = W_X_H1(Spike_X_A,:)' .* H1_A;
        ST = sum(Temp);
        if ST > 10^-100
          H1_A = OneDivOnePlusEps * (H1_A + Epsilon * Temp/ST);
        end

        Temp = W_X_H1(Spike_X_B,:)' .* H1_B;
        ST = sum(Temp);
        if ST > 10^-100
          H1_B = OneDivOnePlusEps * (H1_B + Epsilon * Temp/ST);
        end

        Temp = W_H1_H2(Spike_H1_A,:)' .* H2;
        ST = sum(Temp);
        if ST > 10^-100
          H2 = OneDivOnePlusEps * (H2 + Epsilon * Temp/ST);
        end

        Temp = W_H1_H2(Spike_H1_B+4,:)' .* H2;
        ST = sum(Temp);
        if ST > 10^-100
          H2 = OneDivOnePlusEps * (H2 + Epsilon * Temp/ST);
        end

        Temp = W_H2_HY(Spike_H2,:)' .* HY;
        ST = sum(Temp);
        if ST > 10^-100
          HY = OneDivOnePlusEps * (HY + Epsilon * Temp/ST);
        end

      end

      [Dummy MaxPos_Correct] = max(P_Y);
      [Dummy MaxPos] = max(HY);
    
      Correct_Full(Smooth_ID,1) = Correct_Full(Smooth_ID,1) + double([MaxPos_Correct == MaxPos]);

    end

    %% Updating the weights
    [F_H1_A, T_H1_A, SUM_T_H1_A]    = FrontFactor(P_X(1:4)*2, squeeze(H1_A_Collect(:,PatternID,4)), W_X_H1, Epsilon);
    [F_H1_B, T_H1_B, SUM_T_H1_B]    = FrontFactor(P_X(5:8)*2, squeeze(H1_B_Collect(:,PatternID,4)), W_X_H1, Epsilon);
    [F_H2, T_H2, SUM_T_H2]          = FrontFactor([squeeze(H1_A_Collect(:,PatternID,3)); squeeze(H1_B_Collect(:,PatternID,3))], squeeze(H2_Collect(:,PatternID,3)), W_H1_H2, Epsilon);
    [F_HY, T_HY, SUM_T_HY]          = FrontFactor(squeeze(H2_Collect(:,PatternID,2)), squeeze(HY_Collect(:,PatternID,2)), W_H2_HY, Epsilon);
    [ES]                            = ErrorFactor(squeeze(HY_Collect(:,PatternID,1)), P_Y);

    PropFactor = ES;
    dW_H2_HY = dW_H2_HY + DiffFactor(PropFactor, T_HY, SUM_T_HY, F_HY);

    [PropFactor] = PropagateFactor(T_HY, SUM_T_HY, PropFactor);
    dW_H1_H2  = dW_H1_H2  + DiffFactor(PropFactor, T_H2, SUM_T_H2, F_H2); 

    [PropFactor] = PropagateFactor(T_H2, SUM_T_H2, PropFactor);

    dW_X_H1  = dW_X_H1  + DiffFactor(PropFactor(1:4)*2, T_H1_A, SUM_T_H1_A, F_H1_A); 
    dW_X_H1  = dW_X_H1  + DiffFactor(PropFactor(5:8)*2, T_H1_B, SUM_T_H1_B, F_H1_B); 
    
  end

  if length(Smooth_ID) == 1
    Temp = Patterns_Output .* log(Patterns_Output./squeeze(HY_Collect(:,:,1)));
    Temp(find(Patterns_Output == 0)) = 0;
    KL(Smooth_ID) = sum(Temp(:));
  end

  Scale = max(abs(dW_X_H1(:)));
%  W_X_H1 = W_X_H1   + (Gamma/Scale) * dW_X_H1;
  W_X_H1_N = W_X_H1   + (Gamma/Scale) * W_X_H1 .* dW_X_H1 ;
  W_X_H1_N(find(W_X_H1_N < 0.0001)) = 0.0001;
  W_X_H1_N = W_X_H1_N ./ repmat(sum(W_X_H1_N,1), [size(W_X_H1,1) 1]);

  if sum(isnan(W_X_H1_N(:))) == 0
    W_X_H1 = W_X_H1_N;
  end


  Scale = max(abs(dW_H1_H2(:)));
%  W_H1_H2 = W_H1_H2   + (Gamma/Scale) * dW_H1_H2;
  W_H1_H2_N = W_H1_H2   + (Gamma/Scale) * W_H1_H2 .* dW_H1_H2;
  W_H1_H2_N(find(W_H1_H2_N < 0.0001)) = 0.0001;
  W_H1_H2_N = W_H1_H2_N ./ repmat(sum(W_H1_H2_N,1), [size(W_H1_H2,1) 1]);

  if sum(isnan(W_H1_H2_N(:))) == 0
    W_H1_H2 = W_H1_H2_N;
  end


  Scale = max(abs(dW_H2_HY(:)));
%  W_H2_HY = W_H2_HY + (Gamma/Scale) * dW_H2_HY;
  W_H2_HY_N = W_H2_HY   + (Gamma/Scale) * W_H2_HY .* dW_H2_HY;
  W_H2_HY_N(find(W_H2_HY_N < 0.0001)) = 0.0001;
  W_H2_HY_N = W_H2_HY_N ./ repmat(sum(W_H2_HY_N,1), [size(W_H2_HY,1) 1]);

  if sum(isnan(W_H2_HY_N(:))) == 0
    W_H2_HY = W_H2_HY_N;
  end

  if length(Smooth_ID) == 1
    W_Range(Smooth_ID,1) = min(W_X_H1(:));
    W_Range(Smooth_ID,2) = max(W_X_H1(:));

    W_Range(Smooth_ID,3) = min(W_H1_H2(:));
    W_Range(Smooth_ID,4) = max(W_H1_H2(:));

    W_Range(Smooth_ID,5) = min(W_H2_HY(:));
    W_Range(Smooth_ID,6) = max(W_H2_HY(:));
  end

end

eval(['save Data_A/Data_' num2str(SeedID) '.mat KL W_Range W_X_H1 W_H1_H2 W_H2_HY Smooth_ID Correct Correct_Full']);
```
