## Supplementary material for "Back-propagation learning in deep Spike-By-Spike networks": Source Code: doit.m

```
function paper(SeedID)

rand('seed',SeedID);

NumberIterations_0 = 1024;
NumberIterations = NumberIterations_0;

NumberOfBits = 2;
NumberOfPatterns = 2.^NumberOfBits;
NoN_X  = (NumberOfBits+1) * 2;
NoN_H1 = 4;
NoN_HY = 2;

Epsilon = 0.1;
Gamma = 0.025;

% Lernschritte:
LStep_Max = 10000;
WindowSize = 1000;

Smooth = [1:9 10:2:99 100:5:999 1000:10:LStep_Max];

%%%%%%%%%%%%%

W_X_H1 = ones(NoN_X,NoN_H1) + 1.0/100.0 * rand(NoN_X,NoN_H1);
W_X_H1 = W_X_H1 ./ repmat(sum(W_X_H1,1), [size(W_X_H1,1) 1]);

W_H1_HY = ones(NoN_H1,NoN_HY) + 1.0/100.0 * rand(NoN_H1,NoN_HY);
W_H1_HY = W_H1_HY ./ repmat(sum(W_H1_HY,1), [size(W_H1_HY,1) 1]);

Done = false;

Patterns_Input = zeros(NoN_X-2,NumberOfPatterns);
Patterns_Output = zeros(NoN_HY,NumberOfPatterns);

for i = 1:1:NumberOfPatterns,
    BinString = dec2bin(i-1,NumberOfBits);
    BinString = double(BinString)-48;
    Temp = zeros(NoN_X-2,1);
    Temp([1:2:NoN_X-2]+BinString) = 1;
    Patterns_Input(:,i) = Temp / sum(Temp);
end

Temp = mod(sum([Patterns_Input(1:2:end,:) > 0],1),2)+1;
for i = 1:1:NumberOfPatterns,
  Patterns_Output(Temp(i),i) = 1; 
end

Patterns_Input = Patterns_Input * 2.0 / 3.0;
 
OneDivOnePlusEps = 1 / (1 + Epsilon);

KL = zeros(length(Smooth),1);
W_Range = zeros(length(Smooth),6);
Correct = zeros(length(Smooth),1);
Correct_Full = zeros(length(Smooth),1);

for LStep = 1:1:LStep_Max,
  if mod(LStep,100) == 0
    fprintf([num2str(LStep) ' of ' num2str(LStep_Max) ', Number of Spikes:' num2str(NumberIterations) ' \n']);
  end

  Smooth_ID = find(Smooth == LStep);

  H1_Collect = zeros(NoN_H1,NumberOfPatterns,3);
  HY_Collect = zeros(NoN_HY,NumberOfPatterns,3);

  dW_X_H1 = zeros(size(W_X_H1));
  dW_H1_HY = zeros(size(W_H1_HY));

  if (LStep == 100)
    NumberIterations = fix(NumberIterations / 2);
    Gamma = Gamma / 2.0;
  elseif (LStep == (100+250))
    NumberIterations = fix(NumberIterations / 2);
    Gamma = Gamma / 2.0;
  elseif (LStep == (100+500))
    NumberIterations = fix(NumberIterations / 2);
    Gamma = Gamma / 2.0;
  end

  if Done == true
    NumberIterations = NumberIterations_0;
  end

  AdditionalBits = zeros(2,NumberOfPatterns);
  AdditionalBits(1,:) = rand(NumberOfPatterns,1) > 0.5;
  AdditionalBits(2,:) = AdditionalBits(1,:) == 0;
  Patterns_Input_Mock = [Patterns_Input; AdditionalBits * (1.0/3.0)];

  for PatternID = 1:1:NumberOfPatterns,

    H1 = ones(NoN_H1,1) / NoN_H1;
    HY = ones(NoN_HY,1) / NoN_HY;

    P_X = Patterns_Input_Mock(:,PatternID);
    P_Y = Patterns_Output(:,PatternID);

    %% For learning
    for t = 1:1:NumberIterations,

      Spike_X  = GiveMeASpike(P_X);
      Spike_H1 = GiveMeASpike(H1);
      
      Temp = W_X_H1(Spike_X,:)' .* H1;
      ST = sum(Temp);
      if ST > 10^-100
        H1 = OneDivOnePlusEps * (H1 + Epsilon * Temp/ST);
      end

      Temp = W_H1_HY(Spike_H1,:)' .* HY;
      ST = sum(Temp);
      if ST > 10^-100
        HY = OneDivOnePlusEps * (HY + Epsilon * Temp/ST);
      end

      if t > (NumberIterations-5 )
        H1_Collect(:,PatternID,3) = H1_Collect(:,PatternID,2);
        H1_Collect(:,PatternID,2) = H1_Collect(:,PatternID,1);
        H1_Collect(:,PatternID,1) = H1;

        HY_Collect(:,PatternID,3) = HY_Collect(:,PatternID,2);
        HY_Collect(:,PatternID,2) = HY_Collect(:,PatternID,1);
        HY_Collect(:,PatternID,1) = HY;
      end
    end

    if length(Smooth_ID) == 1
      [Dummy MaxPos_Correct] = max(P_Y);
      [Dummy MaxPos] = max(HY);
    
      Correct(Smooth_ID,1) = Correct(Smooth_ID,1) + double([MaxPos_Correct == MaxPos]);

      H1 = ones(NoN_H1,1) / NoN_H1;
      HY = ones(NoN_HY,1) / NoN_HY;

      %% For testing the performance 
      for t = 1:1:1024,

        Spike_X  = GiveMeASpike(P_X);
        Spike_H1 = GiveMeASpike(H1);
      
        Temp = W_X_H1(Spike_X,:)' .* H1;
        ST = sum(Temp);
        if ST > 10^-100
          H1 = OneDivOnePlusEps * (H1 + Epsilon * Temp/ST);
        end

        Temp = W_H1_HY(Spike_H1,:)' .* HY;
        ST = sum(Temp);
        if ST > 10^-100
          HY = OneDivOnePlusEps * (HY + Epsilon * Temp/ST);
        end

      end

      [Dummy MaxPos_Correct] = max(P_Y);
      [Dummy MaxPos] = max(HY);

      Correct_Full(Smooth_ID,1) = Correct_Full(Smooth_ID,1) + double([MaxPos_Correct == MaxPos]);
    end

    %% Updating the weights

    [F_H1, T_H1, SUM_T_H1]          = FrontFactor(P_X, squeeze(H1_Collect(:,PatternID,3)), W_X_H1, Epsilon);
    [F_HY, T_HY, SUM_T_HY]          = FrontFactor(squeeze(H1_Collect(:,PatternID,2)), squeeze(HY_Collect(:,PatternID,2)), W_H1_HY, Epsilon);
    [ES]                            = ErrorFactor(squeeze(HY_Collect(:,PatternID,1)), P_Y);

    dW_H1_HY = dW_H1_HY + DiffFactor(ES, T_HY, SUM_T_HY, F_HY);

    [PropFactor] = PropagateFactor(T_HY, SUM_T_HY, ES);
    dW_X_H1  = dW_X_H1  + DiffFactor(PropFactor, T_H1, SUM_T_H1, F_H1); 
    
  end

  if length(Smooth_ID) == 1
    Temp = Patterns_Output .* log(Patterns_Output./squeeze(HY_Collect(:,:,1)));
    Temp(find(Patterns_Output == 0)) = 0;
    KL(Smooth_ID) = sum(Temp(:));
  end

  Scale = max(abs(dW_X_H1(:)));
  W_X_H1_N = W_X_H1  + (Gamma/Scale) * W_X_H1 .* dW_X_H1;
  W_X_H1_N(find(W_X_H1_N < 0.0001)) = 0.0001;
  W_X_H1_N = W_X_H1_N ./ repmat(sum(W_X_H1_N,1), [size(W_X_H1,1) 1]);

  if sum(isnan(W_X_H1_N(:))) == 0
    W_X_H1 = W_X_H1_N;
  end

  Scale = max(abs(dW_H1_HY(:)));
  W_H1_HY_N = W_H1_HY + (Gamma/Scale) * W_H1_HY .* dW_H1_HY;
  W_H1_HY_N(find(W_H1_HY_N < 0.0001)) = 0.0001;
  W_H1_HY_N = W_H1_HY_N ./ repmat(sum(W_H1_HY_N,1), [size(W_H1_HY,1) 1]);

  if sum(isnan(W_H1_HY_N(:))) == 0
    W_H1_HY = W_H1_HY_N;
  end

  if length(Smooth_ID) == 1

    W_Range(Smooth_ID,1) = min(min(W_X_H1(1:4,:)));
    W_Range(Smooth_ID,2) = max(max(W_X_H1(1:4,:)));

    W_Range(Smooth_ID,3) = min(W_H1_HY(:));
    W_Range(Smooth_ID,4) = max(W_H1_HY(:));

    W_Range(Smooth_ID,5) = min(min(W_X_H1(5:6,:)));
    W_Range(Smooth_ID,6) = max(max(W_X_H1(5:6,:)));

  end

end

eval(['save Data/Data_' num2str(SeedID) '.mat KL W_Range W_H1_HY W_X_H1 Correct Correct_Full Smooth']);
```
