## Supplementary material for "Back-propagation learning in deep Spike-By-Spike networks": Source Code: plotit.m

```
clear

eval(['load Data/Data_' num2str(1) '.mat Smooth']);

First = true;
Counter = 0;
for SeedID = 1:1:250,

  try 
    eval(['load Data/Data_' num2str(SeedID) '.mat KL W_Range W_H1_HY W_X_H1 Correct Correct_Full']);

    if First == true
      First = false;
      KL_Collect = zeros(size(KL));
      W_Range_Collect = zeros(size(W_Range));
      Correct_Collect = zeros(size(Correct));
      Correct_Full_Collect = zeros(size(Correct_Full));
    end
    KL_Collect = KL_Collect + KL;
    W_Range_Collect = W_Range_Collect + W_Range;
    Correct_Collect = Correct_Collect + Correct;
    Correct_Full_Collect = Correct_Full_Collect + Correct_Full;

    Counter = Counter + 1;
  catch

  end

end
Counter 

Correct = Correct_Collect/(Counter*4);
Correct_Full = Correct_Full_Collect/(Counter*4);
KL = KL_Collect/(Counter);
W_Range = W_Range_Collect/(Counter);

Handle = figure(1);

subplot(3, 1, 1)
hold off
semilogx([100 100],[0 100],'k--')
hold on
semilogx([100 100]+250,[0 100],'k--')
semilogx([100 100]+500,[0 100],'k--')
plot(Smooth,(1.0-Correct)*100.0)
plot(Smooth,(1.0-Correct_Full)*100.0)
ylim([0 60])

xlabel('Learning step')
ylabel('Error [%]')

subplot(3, 1, 2)
hold off
loglog([100 100],[10^-15 10],'k--')
hold on
loglog([100 100]+250,[10^-15 10],'k--')
loglog([100 100]+500,[10^-15 10],'k--')
loglog(Smooth,KL);
axis tight
yticks([10^-15 10^-10 10^-5 1])
ylim([10^-15 10])
xlabel('Learning step')
ylabel('KL')

subplot(3, 1, 3)
hold off
semilogx([100 100],[0 max(W_Range(:))],'k--')
hold on
semilogx([100 100]+250,[0 max(W_Range(:))],'k--')
semilogx([100 100]+500,[0 max(W_Range(:))],'k--')
semilogx(Smooth,W_Range(:,1:2),'r');
semilogx(Smooth,W_Range(:,3:4),'b');
semilogx(Smooth,W_Range(:,5:6),'k');
axis tight
ylim([0 0.55])
yticks([0 0.125 0.25 0.375 0.5])

xlabel('Learning step')
ylabel('Weight range')

print(Handle,'SubPlot_2_1.pdf','-dpdf')

Handle = figure(2);

SeedID = 1;
eval(['load Data/Data_' num2str(SeedID) '.mat W_H1_HY W_X_H1']);

subplot(2, 1, 1)
imagesc(W_X_H1)
colorbar
axis off
axis image

subplot(2, 1, 2)
imagesc(W_H1_HY)
colorbar
axis off
axis image

colormap('hot')

print(Handle,'SubPlot_2_2.pdf','-dpdf')
```
