## Supplementary material for "Back-propagation learning in deep Spike-By-Spike networks": Source Code: Convert_Train.m

```
clear
Images = loadMNISTImages('train-images-idx3-ubyte');
Label = loadMNISTLabels('train-labels-idx1-ubyte');

Processed = zeros(size(Images,1)*2,size(Images,2));
Images = 2 * Images - 1.0;

A = Images;
A(find( A < 0 )) = 0;
B = -Images;
B(find( B < 0 )) = 0;
Processed(1:2:end,:) = B;
Processed(2:2:end,:) = A;
Data_Train = Processed;
Label_Train = Label';

save MINST_Train.mat Data_Train Label_Train
```
