## Supplementary material for "Back-propagation learning in deep Spike-By-Spike networks": Source Code: CookieCutter_Test.m

```
clear
load MINST_Test.mat
BlockSize=100;

Selector = zeros(24,24,50);
LineID = [1:1:10];
LineOffset = 28*2;
for x = 1:1:24,
  for y = 1:1:24,
    Sel = [ (LineID+2*(x-1)+(y-1)*LineOffset) ...
            (LineID+2*(x-1)+(y+0)*LineOffset) ...
            (LineID+2*(x-1)+(y+1)*LineOffset) ...
            (LineID+2*(x-1)+(y+2)*LineOffset) ...
            (LineID+2*(x-1)+(y+3)*LineOffset)];
    Selector(x,y,:) = Sel;
  end
end

i = 1;
j = 1;
for BlockID = 1:1:ceil(size(Data_Test,2)/BlockSize),

  Data = zeros(50,24,24,BlockSize);
  Label = zeros(1,BlockSize);

  for g = 1:1:BlockSize
    fprintf([num2str(g) ' ' num2str(j) '\n'])
    Label(1,i)  = Label_Test(j) + 1;

    for x = 1:1:24,
      for y = 1:1:24,
        Data(:,x,y,i) = Data_Test(squeeze(Selector(x,y,:)),j);
      end
    end

    i = i + 1;
    j = j + 1;
  end

  eval(['save ../Data/Data_Test_Cut_' num2str(BlockID) '.mat Data Label']);
  i = 1;
  Data = zeros(size(Data));
  Label = zeros(1,BlockSize);

end
```
