## Supplementary material for "Back-propagation learning in deep Spike-By-Spike networks": Source Code: README.txt

This set of tools converts the MNIST data into 24 x 24 input probabilites with 50 neurons each. 

Or in other words:
The 28 x 28 pixel images are "pre-convolved" with a 5 x 5 spatial window plus On/Off transformation.

0.) Four files are required from http://yann.lecun.com/exdb/mnist/
Download the files 
http://yann.lecun.com/exdb/mnist/train-images-idx3-ubyte.gz
http://yann.lecun.com/exdb/mnist/train-labels-idx1-ubyte.gz
http://yann.lecun.com/exdb/mnist/t10k-images-idx3-ubyte.gz
http://yann.lecun.com/exdb/mnist/t10k-labels-idx1-ubyte.gz
and extract them.

Two third party Matlab functions are required:loadMNISTImages.m and loadMNISTLabels.m
These can be found under
http://ufldl.stanford.edu/wiki/resources/mnistHelper.zip 

1.) Run Convert_Test.m and Convert_Train.m for doing the On/Off transformation 

2.) Run CookieCutter_Test.m and CookieCutter_Train.m for cutting out the 5 x 5 windows and splitting the whole data set in:
Training data: 600 files with 100 pattern each
Test data: 100 files with 100 pattern each
We did this for parallelization purposes. Make sure that the folder ../Data exisits because the scripts will try to write there the output files.
