## Supplementary material for "Back-propagation learning in deep Spike-By-Spike networks": Source Code: Create_W_H1_H2.m

```
% Pool Weights 1

Number_Of_H1 = 32;
Number_Of_H2 = 32;

W_H1_H2 = zeros(4*Number_Of_H1,Number_Of_H2);
for g = 1:1:Number_Of_H2
  W_H1_H2([0 1 2 3] * Number_Of_H1 + g,g) = 1;
end
W_H1_H2 = W_H1_H2 ./ repmat(sum(W_H1_H2,1), [size(W_H1_H2,1) 1]);

W_H1_H2 = single(W_H1_H2);

save W_H1_H2.mat W_H1_H2;
```
