## Supplementary material for "Back-propagation learning in deep Spike-By-Spike networks": Source Code: Create_W_H3_H4.m

```
% Pool Weights 2

Number_Of_H3 = 64;
Number_Of_H4 = 64;

W_H3_H4 = zeros(4*Number_Of_H3,Number_Of_H4);
for g = 1:1:Number_Of_H4
  W_H3_H4([0 1 2 3] * Number_Of_H3 + g,g) = 1;
end
W_H3_H4 = W_H3_H4 ./ repmat(sum(W_H3_H4,1), [size(W_H3_H4,1) 1]);

W_H3_H4 = single(W_H3_H4);

save W_H3_H4.mat W_H3_H4;
```
