## Supplementary material for "Back-propagation learning in deep Spike-By-Spike networks": Source Code: CreateInitW_H2_H3.m

```
clear
NumberOfH2 = 32*25;
NumberOfH3 = 64;

W_H2_H3 = ones(NumberOfH2, NumberOfH3);
W_H2_H3 = W_H2_H3 + 1.0/100 * rand(size(W_H2_H3));
W_H2_H3 = W_H2_H3 ./ repmat(sum(W_H2_H3,1), [size(W_H2_H3,1) 1]);

W_H2_H3 = single(W_H2_H3);
save W_H2_H3_Iter0.mat W_H2_H3 
```
