## Supplementary material for "Back-propagation learning in deep Spike-By-Spike networks": Source Code: CreateInitW_H4_H5.m

```
clear
NumberOfH4 = 64*4*4;
NumberOfH5 = 1024;

W_H4_H5 = ones(NumberOfH4, NumberOfH5);
W_H4_H5 = W_H4_H5 +1.0/100 * rand(size(W_H4_H5));
W_H4_H5 = W_H4_H5 ./ repmat(sum(W_H4_H5,1), [size(W_H4_H5,1) 1]);

W_H4_H5 = single(W_H4_H5);
save W_H4_H5_Iter0.mat W_H4_H5 
```
