## Supplementary material for "Back-propagation learning in deep Spike-By-Spike networks": Source Code: CreateInitW_H5_HY.m

```
clear
NumberOfHY = 10;
NumberOfH5 = 1024;

W_H5_HY = ones(NumberOfH5,NumberOfHY);
W_H5_HY = W_H5_HY + 1.0/100 * rand(size(W_H5_HY));
W_H5_HY = W_H5_HY ./ repmat(sum(W_H5_HY,1), [size(W_H5_HY,1) 1]);

W_H5_HY = single(W_H5_HY);
save W_H5_HY_Iter0.mat W_H5_HY 
```
