## Supplementary material for "Back-propagation learning in deep Spike-By-Spike networks": Source Code: CreateInitW_X_H1.m

```
clear
NumberOfH = 32;
NumberOfInput = 50;

W_X_H1 = ones(NumberOfInput,NumberOfH);
W_X_H1 = W_X_H1 + 1.0/100 * rand(size(W_X_H1));
W_X_H1 = W_X_H1 ./ repmat(sum(W_X_H1,1), [size(W_X_H1,1) 1]);

W_X_H1 = single(W_X_H1);
save W_X_H1_Iter0.mat W_X_H1
```
