## Supplementary material for "Back-propagation learning in deep Spike-By-Spike networks": Source Code: doit_1_weight_minibatch.m

```
function doit_1a(BlockID, WeightIter)


  rand('seed', BlockID  + WeightIter * 1000)

  Epsilon_F = 0.1;  

  ExistTest = 0;
  try
    eval(['load TempData/Temp_Train_Iter' num2str(WeightIter) '_BlockID' num2str(BlockID) '.mat Pattern_Output ']);
  catch
    ExistTest = 1;
  end

  if ExistTest == 0
    fprintf(['Already done\n']);
    return;
  end

  Number_Of_Spikes = 1200;
  Smooth_After_Num_Of_Spikes = 1000;

  R_X_H1 = 1;
  R_H1_H2 = 4;
  R_H2_H3 = 25;
  R_H3_H4 = 4;
  R_H4_H5 = 16;
  R_H5_HY = 1;

  fprintf([num2str(BlockID)  ' ' num2str(WeightIter) '\n'])
  system('hostname');
  system('pwd');
  mfilename

  NumberOfH_1 = 32;
  NumberOfH_2 = 32;
  NumberOfH_3 = 64;
  NumberOfH_4 = 64;
  NumberOfH_5 = 1024;
  NumberOfOutput = 10;

  Sp_Offset_H2 = zeros(5,5);
  Order_H2 = zeros(25,2);
  for x = 1:1:5,
    for y = 1:1:5,
      Sp_Offset_H2(x,y) = ((x-1) + (y-1) * 5) * NumberOfH_2;
    end
  end
  Temp = (Sp_Offset_H2/NumberOfH_2)+1;
  for i = 1:1:25
    [A,B] = find(Temp == i);
    Order_H2(i,1) = A;
    Order_H2(i,2) = B;
  end

  Sp_Offset_H4 = zeros(4,4);
  w = 0;
  for x = 1:1:4,
    for y = 1:1:4,
      Sp_Offset_H4(x,y) = w * NumberOfH_4;
      w = w + 1;
    end
  end

  Order_H4 = zeros(16,2);
  Temp = (Sp_Offset_H4/NumberOfH_4)+1;
  for i = 1:1:16
    [A,B] = find(Temp == i);
    Order_H4(i,1) = A;
    Order_H4(i,2) = B;
  end

  eval(['load Data/Data_Train_Cut_' num2str(BlockID) '.mat Data Label']);

  RandomPatterns = randperm(size(Data,4));
  RandomPatterns = RandomPatterns(1:10);
  Label = Label(RandomPatterns);

  % Weights
  eval(['load W_X_H1_Iter' num2str(WeightIter) '.mat W_X_H1']);
  W = double(W_X_H1);
  W(find(W < 0.0001)) = 0.0001;
  W = W ./ repmat(sum(W,1),[size(W,1) 1 ]);
  W_X_H1 = single(W);

  eval(['load W_H1_H2.mat W_H1_H2']);
  W = double(W_H1_H2);
  W(find(W < 0.0001)) = 0.0001;
  W = W ./ repmat(sum(W,1),[size(W,1) 1 ]);
  W_H1_H2 = single(W);

  eval(['load W_H2_H3_Iter' num2str(WeightIter) '.mat W_H2_H3']);
  W = double(W_H2_H3);
  W(find(W < 0.0001)) = 0.0001;
  W = W ./ repmat(sum(W,1),[size(W,1) 1 ]);
  W_H2_H3 = single(W);

  eval(['load W_H3_H4.mat W_H3_H4']);
  W = double(W_H3_H4);
  W(find(W < 0.0001)) = 0.0001;
  W = W ./ repmat(sum(W,1),[size(W,1) 1 ]);
  W_H3_H4 = single(W);

  eval(['load W_H4_H5_Iter' num2str(WeightIter) '.mat W_H4_H5']);
  W = double(W_H4_H5);
  W(find(W < 0.0001)) = 0.0001;
  W = W ./ repmat(sum(W,1),[size(W,1) 1 ]);
  W_H4_H5 = single(W);

  eval(['load W_H5_HY_Iter' num2str(WeightIter) '.mat W_H5_HY']);
  W = double(W_H5_HY);
  W(find(W < 0.0001)) = 0.0001;
  W = W ./ repmat(sum(W,1),[size(W,1) 1 ]);
  W_H5_HY = single(W);

  TX = Epsilon_F / R_X_H1;
  Temp = (TX/(1.0-TX))  * ones(Number_Of_Spikes,1);
  Temp(Smooth_After_Num_Of_Spikes:Number_Of_Spikes) = Temp(Smooth_After_Num_Of_Spikes:Number_Of_Spikes)/25;
  Temp = single(Temp);
  Epsilon_X_H1 = Temp;

  TX = Epsilon_F / R_H1_H2;
  Temp = (TX/(1.0-TX))  * ones(Number_Of_Spikes,1);
  Temp(Smooth_After_Num_Of_Spikes:Number_Of_Spikes) = Temp(Smooth_After_Num_Of_Spikes:Number_Of_Spikes)/25;
  Temp = single(Temp);
  Epsilon_H1_H2 = Temp;

  TX = Epsilon_F / R_H2_H3;
  Temp = (TX/(1.0-TX))  * ones(Number_Of_Spikes,1);
  Temp(Smooth_After_Num_Of_Spikes:Number_Of_Spikes) = Temp(Smooth_After_Num_Of_Spikes:Number_Of_Spikes)/25;
  Temp = single(Temp);
  Epsilon_H2_H3 = Temp;

  TX = Epsilon_F / R_H3_H4;
  Temp = (TX/(1.0-TX))  * ones(Number_Of_Spikes,1);
  Temp(Smooth_After_Num_Of_Spikes:Number_Of_Spikes) = Temp(Smooth_After_Num_Of_Spikes:Number_Of_Spikes)/25;
  Temp = single(Temp);
  Epsilon_H3_H4 = Temp;

  TX = Epsilon_F / R_H4_H5;
  Temp = (TX/(1.0-TX))  * ones(Number_Of_Spikes,1);
  Temp(Smooth_After_Num_Of_Spikes:Number_Of_Spikes) = Temp(Smooth_After_Num_Of_Spikes:Number_Of_Spikes)/25;
  Temp = single(Temp);
  Epsilon_H4_H5 = Temp;

  TX = Epsilon_F / R_H5_HY;
  Temp = (TX/(1.0-TX))  * ones(Number_Of_Spikes,1);
  Temp(Smooth_After_Num_Of_Spikes:Number_Of_Spikes) = Temp(Smooth_After_Num_Of_Spikes:Number_Of_Spikes)/25;
  Temp = single(Temp);
  Epsilon_H5_HY = Temp;

  clear Epsilon_F;

  
  for TrialID = 1:1:length(RandomPatterns),%size(Data,4)

    ExistTest = 0;
    try
      eval(['load TempData/Temp_Train_Iter' num2str(WeightIter) '_BlockID' num2str(BlockID) '.mat Pattern_Output ']);
    catch
      ExistTest = 1;
    end

    if ExistTest == 0
      fprintf(['Already done\n']);
      return;
    end

    fprintf(['T:' num2str(TrialID) ' ->' ])
    tic

    Pattern = squeeze(Data(:,:,:,RandomPatterns(TrialID)));
    for x = 1:1:size(Pattern,2)
      for y = 1:1:size(Pattern,3)
        Pattern(:,x,y) = single(Pattern(:,x,y) ./ sum(Pattern(:,x,y)));
      end
    end

%    Pattern = Convert2Pattern(squeeze(Data(:,:,:,TrialID)));

    H1 = ones(NumberOfH_1,24,24);
    H1 = H1 ./ repmat(sum(H1,1),[size(H1,1) 1 1]);
    H1 = single(H1);

    H2 = ones(NumberOfH_2,12,12);
    H2 = H2 ./ repmat(sum(H2,1),[size(H2,1) 1 1]);
    H2 = single(H2);

    H3 = ones(NumberOfH_3,8,8);
    H3 = H3 ./ repmat(sum(H3,1),[size(H3,1) 1 1]);
    H3 = single(H3);

    H4 = ones(NumberOfH_4,4,4);
    H4 = H4 ./ repmat(sum(H4,1),[size(H4,1) 1 1]);
    H4 = single(H4);

    % 1: 1024
    H5 = ones(NumberOfH_5,1);
    H5 = H5 ./ repmat(sum(H5,1),[size(H5,1) 1] );
    H5 = single(H5);

    % 1: 10
    HY = ones(NumberOfOutput,1);
    HY = HY ./ repmat(sum(HY,1),[size(HY,1) 1]);
    HY = single(HY);

    if TrialID == 1

      %Pattern_Input = single(zeros(size(Pattern,1),size(Pattern,2),size(Pattern,3),size(Data,4)));
      %H1_All        = single(zeros(NumberOfH_1,28,28,size(Data,4)));
      %H2_All        = single(zeros(NumberOfH_2,14,14,size(Data,4)));
      %H3_All        = single(zeros(NumberOfH_3,10,10,size(Data,4)));
      %H4_All        = single(zeros(NumberOfH_4,5,5,size(Data,4)));
      %H5_All        = single(zeros(NumberOfH_5,size(Data,4)));
      %HY_All        = single(zeros(NumberOfOutput,size(Data,4)));
      %Pattern_Output= single(zeros(NumberOfOutput,size(Data,4)));

      Pattern_Input = single(zeros(size(Pattern,1),size(Pattern,2),size(Pattern,3),length(RandomPatterns)));
      H1_All        = single(zeros(NumberOfH_1,24,24,length(RandomPatterns)));
      H2_All        = single(zeros(NumberOfH_2,12,12,length(RandomPatterns)));
      H3_All        = single(zeros(NumberOfH_3,8,8,length(RandomPatterns)));
      H4_All        = single(zeros(NumberOfH_4,4,4,length(RandomPatterns)));
      H5_All        = single(zeros(NumberOfH_5,length(RandomPatterns)));
      HY_All        = single(zeros(NumberOfOutput,length(RandomPatterns)));
      Pattern_Output= single(zeros(NumberOfOutput,length(RandomPatterns)));

    end

    for t = 1:1:Number_Of_Spikes,
      if mod(t,100) == 0
        fprintf([num2str(t) ' ']);
      end

      %%% Let's get the spikes for this time step
      Sp_X = zeros(size(Pattern,2),size(Pattern,3));
      for x = 1:1:size(Pattern,2),
        for y = 1:1:size(Pattern,3),
          Sp_X(x,y) = GiveMeASpike(squeeze(Pattern(:,x,y)));
        end
      end

      Sp_H1 = zeros(size(H1,2),size(H1,3));
      for x = 1:1:size(H1,2),
        for y = 1:1:size(H1,3),
          Sp_H1(x,y) = GiveMeASpike(squeeze(H1(:,x,y)));
        end
      end

      Sp_H2 = zeros(size(H2,2),size(H2,3));
      for x = 1:1:size(H2,2),
        for y = 1:1:size(H2,3),
          Sp_H2(x,y) = GiveMeASpike(squeeze(H2(:,x,y)));
        end
      end

      Sp_H3 = zeros(size(H3,2),size(H3,3));
      for x = 1:1:size(H3,2),
        for y = 1:1:size(H3,3),
          Sp_H3(x,y) = GiveMeASpike(squeeze(H3(:,x,y)));
        end
      end

      Sp_H4 = zeros(size(H4,2),size(H4,3));
      for x = 1:1:size(H4,2),
        for y = 1:1:size(H4,3),
          Sp_H4(x,y) = GiveMeASpike(squeeze(H4(:,x,y)));
        end
      end

      Sp_H5 = GiveMeASpike(H5);

      Sp_HY = GiveMeASpike(HY);


      %% ---------------------------------------------------------------

      %% H1 changes ->
      % Input -> H1

      OneDivOnePlusEpsilon = single(1.0) ./ (single(1.0) + Epsilon_X_H1(t));

      for x = 1:1:size(H1,2),
        for y = 1:1:size(H1,3),

          Temp = H1(:,x,y) .* W_X_H1(Sp_X(x,y),:)';
          U = sum(Temp);
          if (U > 10^-30)
            Norm = Epsilon_X_H1(t) / U;
            H1(:,x,y) = (H1(:,x,y) + Norm * Temp) * OneDivOnePlusEpsilon;
          end

        end
      end

      %% <- H1 changes

      %% ---------------------------------------------------------------

      %% H2 changes ->

      % H1 -> H2
      % Pool forward
      OneDivOnePlusEpsilon = single(1.0) ./ (single(1.0) + Epsilon_H1_H2(t));

      for x = 1:1:size(H2,2),
        for y = 1:1:size(H2,3),
          for x_s = 1:1:2,
            for y_s = 1:1:2,

              Temp = H2(:,x,y) .* squeeze(W_H1_H2(Sp_H1((x-1)*2+x_s,(y-1)*2+y_s)+((x_s-1)*2+(y_s-1))*NumberOfH_1,:))';
              U = sum(Temp);
              if (U > 10^-30)
                Norm = Epsilon_H1_H2(t) / U;
                H2(:,x,y) = (H2(:,x,y) + Norm * Temp) * OneDivOnePlusEpsilon;
              end

            end
          end
        end
      end
      %% <- H2 changes

      %% ---------------------------------------------------------------

      %% H3 changes ->

      % H2 -> H3

      OneDivOnePlusEpsilon = single(1.0) ./ (single(1.0) + Epsilon_H2_H3(t));

      for x = 1:1:size(H3,2),
        for y = 1:1:size(H3,3),

          Sp_H2_Sub = Sp_H2(x:x+4,y:y+4) + Sp_Offset_H2;
          for i = 1:1:5,
            for j = 1:1:5,

              Temp = H3(:,x,y) .* W_H2_H3(Sp_H2_Sub(i,j),:)';
              U = sum(Temp);
              if (U > 10^-30)
                Norm = Epsilon_H2_H3(t) / U;
                H3(:,x,y) = (H3(:,x,y) + Norm * Temp) * OneDivOnePlusEpsilon;
              end

            end
          end

        end
      end

      %% <- H3 changes

      %% ---------------------------------------------------------------

      %% H4 changes ->

      % H3 -> H4
      OneDivOnePlusEpsilon = single(1.0) ./ (single(1.0) + Epsilon_H3_H4(t));

      for x = 1:1:size(H4,2),
        for y = 1:1:size(H4,3),
          for x_s = 1:1:2,
            for y_s = 1:1:2,

              Temp = H4(:,x,y) .* squeeze(W_H3_H4(Sp_H3((x-1)*2+x_s,(y-1)*2+y_s)+((x_s-1)*2+(y_s-1))*NumberOfH_3,:))';
              U = sum(Temp);
              if (U > 10^-30)
                Norm = Epsilon_H3_H4(t) / U;
                H4(:,x,y) = (H4(:,x,y) + Norm * Temp) * OneDivOnePlusEpsilon;
              end

            end
          end
        end
      end
      %% <- H4 changes

      %% ---------------------------------------------------------------

      %% H5 changes ->

      % H4 -> H5
      OneDivOnePlusEpsilon = single(1.0) ./ (single(1.0) + Epsilon_H4_H5(t));

      for i = 1:1:4,
        for j = 1:1:4,

          Temp = H5 .* squeeze(W_H4_H5(Sp_H4(i,j)+Sp_Offset_H4(i,j),:))';
          U = sum(Temp);
          if (U > 10^-30)
            Norm = Epsilon_H4_H5(t) / U;
            H5 = (H5 + Norm * Temp) * OneDivOnePlusEpsilon;
          end
        end
      end
      %% <- H5 changes

      %% ---------------------------------------------------------------

      %% HY changes ->

      % H5 (1 Spike) -> HY
      OneDivOnePlusEpsilon = single(1.0) ./ (single(1.0) + Epsilon_H5_HY(t));
      Temp = HY .* W_H5_HY(Sp_H5,:)';
      U = sum(Temp);
      if (U > 10^-30)
        Norm = Epsilon_H5_HY(t) / U;
        HY = (HY + Norm * Temp) * OneDivOnePlusEpsilon;
      end
      

      %% <- HY changes

      %% ---------------------------------------------------------------

    end

    Pattern_Input(:,:,:,TrialID)            = Pattern; 
    H1_All(:,:,:,TrialID)                   = H1;
    H2_All(:,:,:,TrialID)                   = H2; 
    H3_All(:,:,:,TrialID)                   = H3; 
    H4_All(:,:,:,TrialID)                   = H4; 
    H5_All(:,TrialID)                       = H5; 
    HY_All(:,TrialID)                       = HY;
    Pattern_Output(Label(TrialID),TrialID)  = 1;
    
    fprintf([num2str(toc) ' \n']);


  end

  % For the save:
  eval(['save TempData/Temp_Train_Iter' num2str(WeightIter) '_BlockID' num2str(BlockID) '.mat  Pattern_Input H1_All H2_All H3_All H4_All H5_All HY_All Pattern_Output ']);


  %%%%%%%%%%%%%%%%%%%%%%%%%%%%%%%%%%%%%%%%%%%%%%%%%%%%%%%%%%%%%%%%%%%%%%%%%%%%%%%%%%%%%%%%%%%%%%%%%%%%%%%%%%%%%%%
  %%%%%%%%%%%%%%%%%%%%%%%%%%%%%%%%%%%%%%%%%%%%%%%%%%%%%%%%%%%%%%%%%%%%%%%%%%%%%%%%%%%%%%%%%%%%%%%%%%%%%%%%%%%%%%%
  %%%%%%%%%%%%%%%%%%%%%%%%%%%%%%%%%%%%%%%%%%%%%%%%%%%%%%%%%%%%%%%%%%%%%%%%%%%%%%%%%%%%%%%%%%%%%%%%%%%%%%%%%%%%%%%
  %%%%%%%%%%%%%%%%%%%%%%%%%%%%%%%%%%%%%%%%%%%%%%%%%%%%%%%%%%%%%%%%%%%%%%%%%%%%%%%%%%%%%%%%%%%%%%%%%%%%%%%%%%%%%%%
  %%%%%%%%%%%%%%%%%%%%%%%%%%%%%%%%%%%%%%%%%%%%%%%%%%%%%%%%%%%%%%%%%%%%%%%%%%%%%%%%%%%%%%%%%%%%%%%%%%%%%%%%%%%%%%%
```
