## Supplementary material for "Back-propagation learning in deep Spike-By-Spike networks": Source Code: GiveMeASpike.m

```
function [Spike] = GiveMeASpike(Distribution)
  
  RandomNumber = single(rand);

  for i = 1:1:length(Distribution),

    RandomNumber = RandomNumber - Distribution(i);

    if (RandomNumber <= 0)
      Spike = i;
      return;
    end

  end

  Spike = length(Distribution);
```
