## Supplementary material for "Back-propagation learning in deep Spike-By-Spike networks": Source Code: joinplot.m

```
function [M] = joinplot(Data,DimA,DimB)

A = size(Data,1);
B = size(Data,2);

M = zeros(DimA*(A+1)+1,DimB*(B+1)+1,1);% * max(Data(:));

Counter_Sub = 1;
for x = 1:1:DimB
  for y = 1:1:DimA
    D = squeeze(Data(:,:,:,Counter_Sub));
    D = D - min(D(:));
    D = D / max(D(:));
%    D = (D * 2) - 1;
    M(1+(y-1)*(A+1)+[1:A], 1+(x-1)*(B+1)+[1:B],:) = D;
    Counter_Sub = Counter_Sub + 1;
  end
end
```
