## Supplementary material for "Back-propagation learning in deep Spike-By-Spike networks": Source Code: plotit_L.m

```
function plot_development(NoE)

figure(1)

try
  load Stat_Plot_Last Last;
  W_Start = size(Last,1);
  Results = zeros(max([NoE W_Start]),3);
  Results(1:W_Start,:) = Last;
catch
  W_Start = 1;
  Results = zeros(NoE,3);
end

for WeightIter = W_Start:1:NoE,
  if WeightIter <= 0
    eval(['load Stat/Stat_Iter' num2str(WeightIter) '.mat L_New L LearnRate']);
  else
    eval(['load Stat_Iter' num2str(WeightIter) '.mat L_New L LearnRate']);
  end
  Results(WeightIter,1) = L_New;
  Results(WeightIter,2) = L;
  Results(WeightIter,3) = LearnRate;
end

Last = Results;
save Stat_Plot_Last Last;
subplot(3,1,1)
semilogy([1:1:NoE],Results(:,1:2),'.-')
xlabel('Learning step')
ylabel('KL of Mini-Batch')

subplot(3,1,2)
semilogy([1:1:NoE],Results(:,3),'.-')
xlabel('Learning step')
ylabel('Learning rate')

NoE_10 = fix(NoE / 10);

Results = zeros(NoE_10+1,1);
for WeightIter = 0:1:NoE_10,
  try
    eval(['load test_Iter' num2str(WeightIter*10) '.mat Result']);
    Results(WeightIter+1) = Result;
  catch
    Results(WeightIter+1) = 10;
  end
end

subplot(3,1,3)
semilogy(([0:1:NoE_10])*10,100.0-Results,'*-')
xlabel('Learning step')
ylabel('Error[%]')
```
