## Supplementary material for "Back-propagation learning in deep Spike-By-Spike networks": Source Code: plotit_Layer1.m

```
function plotit(WeightIter)

eval(['load W_X_H1_Iter' num2str(WeightIter) '.mat ']);

A = W_X_H1(2:2:50,:) - W_X_H1(1:2:50,:);

%for i = 1:1:32
%  subplot(4,8,i)
%  B = A(:,i);
%  B = B - min(B(:));
%  B = B / max(B(:));
%  B = (2 * B) - 1;
%
%end

figure(WeightIter+1)

Image = zeros(5,5,1,32);
for i = 1:1:32
  Image(:,:,1,i) = reshape(A(:,i),[5 5]);
end

[M] = joinplot(Image,4,8);
imagesc(M);
axis image
axis off
title([num2str(WeightIter)])

colormap('hot')
```
