## Supplementary material for "Back-propagation learning in deep Spike-By-Spike networks": Source Code: README.txt

0.) Perform the data preparation procedure.

1.) Create the inital random weight matrices
CreateInitW_X_H1
Create_W_H1_H2
CreateInitW_H2_H3
Create_W_H3_H4
CreateInitW_H4_H5
CreateInitW_H5_HY

2.) Learn the weights:
for WeightIteration = 0:1:149,
  for BlockID = 1:1:600,
    doit_1_weight_minibatch(BlockID,WeightIteration);
    Update_Weights_Sub(BlockID,WeightIteration);
  end
  Update_Weights(WeightIteration);
end
The inner loop can be distributed over many computers. 


Update_Weights_Sub.m uses the following mini sub-functions DiffFactor.m, ErrorFactor.m, FrontFactor.m, and PropagateFactor.m for the back-propation. 

GiveMeASpike.m is used for producing spikes. 

3.) Performance on the test data set can be calculated for weights from learning step 'WeightIteration' via:
for BlockID = 1:1:100,
  doit_1_test(BlockID,WeightIteration);
end
testit_loader(WeightIteration)

4.) The weight matrix between the input layer and the first hidden layer H1 can be plotted via the function plotit_Layer1.m (which use joinplot.m as a sub-function).
plotit_L.m plots the progress of the learning. Test errors are plotted in steps of 10 mini batches. 
If a performance value is not present it is substituted by a 10% performance value (=> chance level).
