## Supplementary material for "Back-propagation learning in deep Spike-By-Spike networks": Source Code: testit_loader.m

```
function testit_loader(WeightIter)
AllCorr = 0;
AllCount = 0;

for BlockID = 1:1:100
  try
    eval(['load Test/Temp_Iter' num2str(WeightIter) '_BlockID' num2str(BlockID) '.mat Correct Count']);
    AllCorr = AllCorr + Correct;
    AllCount = AllCount + Count;

  catch
    BlockID
  end
end
[AllCorr AllCount]
100 * AllCorr/AllCount

if AllCount == 10000
  Result = 100 * AllCorr/AllCount;

  eval(['save test_Iter' num2str(WeightIter) '.mat Result']);
end
```
