## Supplementary material for "Back-propagation learning in deep Spike-By-Spike networks": Source Code: Update_Weights.m

```
function Update_Weights(WeightIter)
%clear
%WeightIter = 0;

%if WeightIter < 150
  Gamma = 0.05;
%else
%  UUU = fix(WeightIter/10)-13;
%  Gamma = (0.05)*UUU.^(-1/2);
%end


NumberOfH_1 = 32;
NumberOfH_2 = 32;
NumberOfH_3 = 64;
NumberOfH_4 = 64;
NumberOfH_5 = 1024;
NumberOfOutput = 10;

if WeightIter == 0
  Para_Tau_M = 10;
else
  eval(['load Stat_Iter' num2str(WeightIter) '.mat L L_Max L_New L_M Para_Tau_M dW_X_H1_M dW_H1_H2_M dW_H2_H3_M dW_H3_H4_M dW_H4_H5_M dW_H5_HY_M']);
  L_Old = L_New;
end


% Weights
eval(['load W_X_H1_Iter' num2str(WeightIter) '.mat W_X_H1']);
W = double(W_X_H1);
W(find(W < 0.0001)) = 0.0001;
W = W ./ repmat(sum(W,1),[size(W,1) 1 ]);
W_X_H1 = W;
TdW_X_H1 = zeros(size(W));

eval(['load W_H1_H2.mat W_H1_H2']);
W = double(W_H1_H2);
W(find(W < 0.0001)) = 0.0001;
W = W ./ repmat(sum(W,1),[size(W,1) 1 ]);
W_H1_H2 = W;
TdW_H1_H2 = zeros(size(W));

eval(['load W_H2_H3_Iter' num2str(WeightIter) '.mat W_H2_H3']);
W = double(W_H2_H3);
W(find(W < 0.0001)) = 0.0001;
W = W ./ repmat(sum(W,1),[size(W,1) 1 ]);
W_H2_H3 = W;
TdW_H2_H3 = zeros(size(W));


eval(['load W_H3_H4.mat W_H3_H4']);
W = double(W_H3_H4);
W(find(W < 0.0001)) = 0.0001;
W = W ./ repmat(sum(W,1),[size(W,1) 1 ]);
W_H3_H4 = W;
TdW_H3_H4 = zeros(size(W));


eval(['load W_H4_H5_Iter' num2str(WeightIter) '.mat W_H4_H5']);
W = double(W_H4_H5);
W(find(W < 0.0001)) = 0.0001;
W = W ./ repmat(sum(W,1),[size(W,1) 1 ]);
W_H4_H5 = W;
TdW_H4_H5 = zeros(size(W));

    
eval(['load W_H5_HY_Iter' num2str(WeightIter) '.mat W_H5_HY']);
W = double(W_H5_HY);
W(find(W < 0.0001)) = 0.0001;
W = W ./ repmat(sum(W,1),[size(W,1) 1 ]);
W_H5_HY = W;
TdW_H5_HY = zeros(size(W));

Sp_Offset_H2 = zeros(5,5);
Order_H2 = zeros(25,2);
for x = 1:1:5,
  for y = 1:1:5,
    Sp_Offset_H2(x,y) = ((x-1) + (y-1) * 5) * NumberOfH_2;
  end
end

Temp = (Sp_Offset_H2/NumberOfH_2)+1;
for i = 1:1:25
  [A,B] = find(Temp == i);
  Order_H2(i,1) = A;
  Order_H2(i,2) = B;
end

Sp_Offset_H4 = zeros(4,4);
w = 0;
for x = 1:1:4,
  for y = 1:1:4,
    Sp_Offset_H4(x,y) = w * NumberOfH_4;
    w = w + 1;
  end
end
  
Order_H4 = zeros(16,2);
Temp = (Sp_Offset_H4/NumberOfH_4)+1;
for i = 1:1:16
  [A,B] = find(Temp == i);
  Order_H4(i,1) = A;
  Order_H4(i,2) = B;
end

TL = 0;
TL_Counter = 0;

for BlockID = 1:1:600,
  BlockID 
  eval(['load  WSub/Temp_Iter' num2str(WeightIter) '_BlockID' num2str(BlockID) '.mat ']);
  TL = TL + L;
  TL_Counter = TL_Counter + 1;

  TdW_X_H1  = TdW_X_H1  + dW_X_H1;
  TdW_H1_H2 = TdW_H1_H2 + dW_H1_H2;
  TdW_H2_H3 = TdW_H2_H3 + dW_H2_H3;
  TdW_H3_H4 = TdW_H3_H4 + dW_H3_H4;
  TdW_H4_H5 = TdW_H4_H5 + dW_H4_H5;
  TdW_H5_HY = TdW_H5_HY + dW_H5_HY;

end
  L_New = TL / TL_Counter;

  if isfinite(L_New) ~= 1
    L_New = L_Old;
    WarningFlag = 1;
  else
    WarningFlag = 0;
  end

  NdW_X_H1  = TdW_X_H1  .* W_X_H1  * (1.0 / Para_Tau_M);
  NdW_H1_H2 = TdW_H1_H2 .* W_H1_H2 * (1.0 / Para_Tau_M);
  NdW_H2_H3 = TdW_H2_H3 .* W_H2_H3 * (1.0 / Para_Tau_M);
  NdW_H3_H4 = TdW_H3_H4 .* W_H3_H4 * (1.0 / Para_Tau_M);
  NdW_H4_H5 = TdW_H4_H5 .* W_H4_H5 * (1.0 / Para_Tau_M);
  NdW_H5_HY = TdW_H5_HY .* W_H5_HY * (1.0 / Para_Tau_M);

  if WeightIter == 0
    L_M = L_New * (1.0 / Para_Tau_M);

    dW_X_H1_M  = NdW_X_H1;
    dW_H1_H2_M = NdW_H1_H2;
    dW_H2_H3_M = NdW_H2_H3;
    dW_H3_H4_M = NdW_H3_H4;
    dW_H4_H5_M = NdW_H4_H5;
    dW_H5_HY_M = NdW_H5_HY;
  else

    L_M = L_M * (1.0 - (1.0 / Para_Tau_M)) + L_New * (1.0 / Para_Tau_M);

    dW_X_H1_M  = dW_X_H1_M  * (1.0 - (1.0 / Para_Tau_M)) + NdW_X_H1;
    dW_H1_H2_M = dW_H1_H2_M * (1.0 - (1.0 / Para_Tau_M)) + NdW_H1_H2;
    dW_H2_H3_M = dW_H2_H3_M * (1.0 - (1.0 / Para_Tau_M)) + NdW_H2_H3;
    dW_H3_H4_M = dW_H3_H4_M * (1.0 - (1.0 / Para_Tau_M)) + NdW_H3_H4;
    dW_H4_H5_M = dW_H4_H5_M * (1.0 - (1.0 / Para_Tau_M)) + NdW_H4_H5;
    dW_H5_HY_M = dW_H5_HY_M * (1.0 - (1.0 / Para_Tau_M)) + NdW_H5_HY;
  end

  FactorX = 1.0 - (1 - (1.0 / Para_Tau_M))^(WeightIter+1) ;
  dW_X_H1  = dW_X_H1_M  / FactorX;
  dW_H1_H2 = dW_H1_H2_M / FactorX;
  dW_H2_H3 = dW_H2_H3_M / FactorX;
  dW_H3_H4 = dW_H3_H4_M / FactorX;
  dW_H4_H5 = dW_H4_H5_M / FactorX;
  dW_H5_HY = dW_H5_HY_M / FactorX;
  L = L_M  / FactorX;

  dW_X_H1  = dW_X_H1  / min([1.0 max(abs(dW_X_H1(:)))]);
  dW_H1_H2 = dW_H1_H2 / min([1.0 max(abs(dW_H1_H2(:)))]);
  dW_H2_H3 = dW_H2_H3 / min([1.0 max(abs(dW_H2_H3(:)))]);
  dW_H3_H4 = dW_H3_H4 / min([1.0 max(abs(dW_H3_H4(:)))]);
  dW_H4_H5 = dW_H4_H5 / min([1.0 max(abs(dW_H4_H5(:)))]);
  dW_H5_HY = dW_H5_HY / min([1.0 max(abs(dW_H5_HY(:)))]);

  if WeightIter == 0
    L_Max = L;
  end
  L_Max = max([L_Max L]);  

  LearnRate = Gamma * sqrt(L / L_Max);

  Name = '_X_H1';
  eval(['W = W' Name ';']);
  eval(['dW = dW' Name ';']);
  W_OLD = W;
  W = W + LearnRate *  dW;
  Idx = find((W_OLD/2) > W);
  W(Idx) = W_OLD(Idx)/2;
  W(find(W < 0.0001)) = 0.0001;
  W = W ./ repmat(sum(W,1), [size(W,1) 1]);

  if sum(isnan(W(:))) == 0
    eval(['W' Name ' = single(W);']);
  end

  %%%%%%%%%%%%%%%%%%%%%%%%%%%%%%%%%%%%%%%%%%%%%%%%%%%%

  Name = '_H1_H2';
  eval(['W = W' Name ';']);
  eval(['dW = dW' Name ';']);
  W_OLD = W;
  W = W + LearnRate *  dW;
  Idx = find((W_OLD/2) > W);
  W(Idx) = W_OLD(Idx)/2;
  W(find(W < 0.0001)) = 0.0001;
  W = W ./ repmat(sum(W,1), [size(W,1) 1]);

  if sum(isnan(W(:))) == 0
    eval(['W' Name ' = single(W);']);
  end

  %%%%%%%%%%%%%%%%%%%%%%%%%%%%%%%%%%%%%%%%%%%%%%%%%%%%

  Name = '_H2_H3';
  eval(['W = W' Name ';']);
  eval(['dW = dW' Name ';']);
  W_OLD = W;
  W = W + LearnRate *  dW;
  Idx = find((W_OLD/2) > W);
  W(Idx) = W_OLD(Idx)/2;
  W(find(W < 0.0001)) = 0.0001;
  W = W ./ repmat(sum(W,1), [size(W,1) 1]);

  if sum(isnan(W(:))) == 0
    eval(['W' Name ' = single(W);']);
  end

  %%%%%%%%%%%%%%%%%%%%%%%%%%%%%%%%%%%%%%%%%%%%%%%%%%%%

  Name = '_H3_H4';
  eval(['W = W' Name ';']);
  eval(['dW = dW' Name ';']);
  W_OLD = W;
  W = W + LearnRate *  dW;
  Idx = find((W_OLD/2) > W);
  W(Idx) = W_OLD(Idx)/2;
  W(find(W < 0.0001)) = 0.0001;
  W = W ./ repmat(sum(W,1), [size(W,1) 1]);

  if sum(isnan(W(:))) == 0
    eval(['W' Name ' = single(W);']);
  end

  %%%%%%%%%%%%%%%%%%%%%%%%%%%%%%%%%%%%%%%%%%%%%%%%%%%%

  Name = '_H4_H5';
  eval(['W = W' Name ';']);
  eval(['dW = dW' Name ';']);
  W_OLD = W;
  W = W + LearnRate *  dW;
  Idx = find((W_OLD/2) > W);
  W(Idx) = W_OLD(Idx)/2;
  W(find(W < 0.0001)) = 0.0001;
  W = W ./ repmat(sum(W,1), [size(W,1) 1]);

  if sum(isnan(W(:))) == 0
    eval(['W' Name ' = single(W);']);
  end

  %%%%%%%%%%%%%%%%%%%%%%%%%%%%%%%%%%%%%%%%%%%%%%%%%%%%

  Name = '_H5_HY';
  eval(['W = W' Name ';']);
  eval(['dW = dW' Name ';']);
  W_OLD = W;
  W = W + LearnRate *  dW;
  Idx = find((W_OLD/2) > W);
  W(Idx) = W_OLD(Idx)/2;
  W(find(W < 0.0001)) = 0.0001;
  W = W ./ repmat(sum(W,1), [size(W,1) 1]);

  if sum(isnan(W(:))) == 0
    eval(['W' Name ' = single(W);']);
  end

  %%%%%%%%%%%%%%%%%%%%%%%%%%%%%%%%%%%%%%%%%%%%%%%%%%%%

  eval(['save W_X_H1_Iter' num2str(WeightIter+1) '.mat W_X_H1']);
%  eval(['save W_H1_H2_Iter' num2str(WeightIter+1) '.mat W_H1_H2']);
  eval(['save W_H2_H3_Iter' num2str(WeightIter+1) '.mat W_H2_H3']);
%  eval(['save W_H3_H4_Iter' num2str(WeightIter+1) '.mat W_H3_H4']);
  eval(['save W_H4_H5_Iter' num2str(WeightIter+1) '.mat W_H4_H5']);
  eval(['save W_H5_HY_Iter' num2str(WeightIter+1) '.mat W_H5_HY']);
  eval(['save Stat_Iter' num2str(WeightIter+1) '.mat L L_Max L_New Para_Tau_M L_M dW_X_H1_M dW_H1_H2_M dW_H2_H3_M dW_H3_H4_M dW_H4_H5_M dW_H5_HY_M WarningFlag LearnRate']);
```
