## Supplementary material for "Back-propagation learning in deep Spike-By-Spike networks": Source Code: Update_Weights_Sub.m

```
function Update_Weights_Sub(BlockID ,WeightIter)
%clear
%WeightIter = 0;
Gamma = 0.025;

Epsilon = 0.1;
Epsilon =  Epsilon / (1.0 - Epsilon);

NumberOfH_1 = 32;
NumberOfH_2 = 32;
NumberOfH_3 = 64;
NumberOfH_4 = 64;
NumberOfH_5 = 1024;
NumberOfOutput = 10;

  ExistTest = 0;
  try
    eval(['load  WSub/Temp_Iter' num2str(WeightIter) '_BlockID' num2str(BlockID) '.mat dW_X_H1 ']);
  catch
    ExistTest = 1;
  end
    
  if ExistTest == 0
    fprintf(['Already done\n']);
    return;
  end


% Weights
eval(['load W_X_H1_Iter' num2str(WeightIter) '.mat W_X_H1']);
W = double(W_X_H1);
W(find(W < 0.0001)) = 0.0001;
W = W ./ repmat(sum(W,1),[size(W,1) 1 ]);
W_X_H1 = W;
dW_X_H1 = zeros(size(W));

eval(['load W_H1_H2.mat W_H1_H2']);
W = double(W_H1_H2);
W(find(W < 0.0001)) = 0.0001;
W = W ./ repmat(sum(W,1),[size(W,1) 1 ]);
W_H1_H2 = W;
dW_H1_H2 = zeros(size(W));

eval(['load W_H2_H3_Iter' num2str(WeightIter) '.mat W_H2_H3']);
W = double(W_H2_H3);
W(find(W < 0.0001)) = 0.0001;
W = W ./ repmat(sum(W,1),[size(W,1) 1 ]);
W_H2_H3 = W;
dW_H2_H3 = zeros(size(W));


eval(['load W_H3_H4.mat W_H3_H4']);
W = double(W_H3_H4);
W(find(W < 0.0001)) = 0.0001;
W = W ./ repmat(sum(W,1),[size(W,1) 1 ]);
W_H3_H4 = W;
dW_H3_H4 = zeros(size(W));


eval(['load W_H4_H5_Iter' num2str(WeightIter) '.mat W_H4_H5']);
W = double(W_H4_H5);
W(find(W < 0.0001)) = 0.0001;
W = W ./ repmat(sum(W,1),[size(W,1) 1 ]);
W_H4_H5 = W;
dW_H4_H5 = zeros(size(W));

    
eval(['load W_H5_HY_Iter' num2str(WeightIter) '.mat W_H5_HY']);
W = double(W_H5_HY);
W(find(W < 0.0001)) = 0.0001;
W = W ./ repmat(sum(W,1),[size(W,1) 1 ]);
W_H5_HY = W;
dW_H5_HY = zeros(size(W));

Sp_Offset_H2 = zeros(5,5);
Order_H2 = zeros(25,2);
for x = 1:1:5,
  for y = 1:1:5,
    Sp_Offset_H2(x,y) = ((x-1) + (y-1) * 5) * NumberOfH_2;
  end
end

Temp = (Sp_Offset_H2/NumberOfH_2)+1;
for i = 1:1:25
  [A,B] = find(Temp == i);
  Order_H2(i,1) = A;
  Order_H2(i,2) = B;
end

Sp_Offset_H4 = zeros(4,4);
w = 0;
for x = 1:1:4,
  for y = 1:1:4,
    Sp_Offset_H4(x,y) = w * NumberOfH_4;
    w = w + 1;
  end
end
  
Order_H4 = zeros(16,2);
Temp = (Sp_Offset_H4/NumberOfH_4)+1;
for i = 1:1:16
  [A,B] = find(Temp == i);
  Order_H4(i,1) = A;
  Order_H4(i,2) = B;
end


%for BlockID = 1:1:600,
  BlockID 

  eval(['load TempData/Temp_Train_Iter' num2str(WeightIter) '_BlockID' num2str(BlockID) '.mat  Pattern_Input H1_All H2_All H3_All H4_All H5_All HY_All Pattern_Output ']);

  U = Pattern_Output .* log(Pattern_Output./HY_All);
  U(find(isnan(U))) = 0;
  U(find(isinf(U))) = 0;

  L = mean(sum(U,1));

  for TID = 1:1:size(Pattern_Output,2)

    P_X   = double(squeeze(Pattern_Input(:,:,:,TID)));
    H1    = double(squeeze(H1_All(:,:,:,TID)));
    H2    = double(squeeze(H2_All(:,:,:,TID)));
    H3    = double(squeeze(H3_All(:,:,:,TID)));
    H4    = double(squeeze(H4_All(:,:,:,TID)));
    H5    = double(squeeze(H5_All(:,TID)));
    HY    = double(squeeze(HY_All(:,TID)));
    P_Y   = double(squeeze(Pattern_Output(:,TID)));

    % HY
    [ES]                              = ErrorFactor(HY, P_Y);
    [F_HY, T_HY, SUM_T_HY]            = FrontFactor(H5, HY, W_H5_HY, Epsilon);

    PropFactor = ES;
    dW_H5_HY = dW_H5_HY + DiffFactor(PropFactor, T_HY, SUM_T_HY, F_HY);

    % H5
    H4_Comby = double([]);
    for a = 1:1:16
      H4_Comby = [H4_Comby ; H4(:,Order_H4(a,1),Order_H4(a,2))];
    end
    H4_Comby = H4_Comby / sum(H4_Comby);
    [F_H5, T_H5, SUM_T_H5]            = FrontFactor(H4_Comby, H5, W_H4_H5, Epsilon);

    [PropFactor] = PropagateFactor(T_HY, SUM_T_HY, PropFactor);

    dW_H4_H5  = dW_H4_H5  + DiffFactor(PropFactor, T_H5, SUM_T_H5, F_H5);

    [PropFactor] = PropagateFactor(T_H5, SUM_T_H5, PropFactor);

    Temp = cell(size(H4_All,2),size(H4_All,3));
    for x = 1:1:size(H4_All,2),
      for y = 1:1:size(H4_All,3),    
        Temp{x,y} = PropFactor(Sp_Offset_H4(x,y)+1:Sp_Offset_H4(x,y)+size(H4_All,1)) / numel(Sp_Offset_H4);
      end
    end
    PropFactor = Temp;

    % H4
    F_H4 = cell(size(H4,2),size(H4,3));
    T_H4 = cell(size(H4,2),size(H4,3));
    SUM_T_H4 = cell(size(H4,2),size(H4,3));       

    PropFactor_New = cell(size(H4,2),size(H4,3));   

    for x = 1:1:size(H4,2)
      for y = 1:1:size(H4,3)
        H3_Sub = H3(:,(x-1)*2+[1:2],(y-1)*2+[1:2]);
        H3_Flat = zeros(numel(H3_Sub),1);
        for x_s = 1:1:2,
          for y_s = 1:1:2,
            H3_Flat(((x_s-1)*2+(y_s-1))*NumberOfH_3+[1:NumberOfH_3]) = H3_Sub(:,x_s,y_s);
          end
        end
        [F_H4{x,y}, T_H4{x,y}, SUM_T_H4{x,y}]  = FrontFactor(H3_Flat, H4(:,x,y), W_H3_H4, Epsilon);

        dW_H3_H4  = dW_H3_H4  + DiffFactor(PropFactor{x,y}, T_H4{x,y}, SUM_T_H4{x,y}, F_H4{x,y});

        [PropFactor_New{x,y}] = PropagateFactor(T_H4{x,y}, SUM_T_H4{x,y}, PropFactor{x,y});
      end
    end
    PropFactor = PropFactor_New;

    PropFactor_New = cell(size(H3,2),size(H3,3));   

    for x = 1:1:size(H4,2)
      for y = 1:1:size(H4,3)
        for x_s = 1:1:2,
          for y_s = 1:1:2,
            Temp = PropFactor{x,y};
            PropFactor_New{(x-1)*2+x_s, (y-1)*2+y_s} = Temp(((x_s-1)*2+(y_s-1))*NumberOfH_4+[1:NumberOfH_4]);
          end
        end
      end
    end
    PropFactor = PropFactor_New;    

    % H3 
    F_H3 = cell(size(H3,2),size(H3,3));
    T_H3 = cell(size(H3,2),size(H3,3));
    SUM_T_H3 = cell(size(H3,2),size(H3,3));       

    PropFactor_New = cell(size(H3,2),size(H3,3));   

    for x = 1:1:size(H3,2)
      for y = 1:1:size(H3,3)

        H2_Comby = double([]);
        for a = 1:1:25
          H2_Comby = [H2_Comby ; H2(:,x+Order_H2(a,1)-1,y+Order_H2(a,2)-1)];
        end
        [F_H3{x,y}, T_H3{x,y}, SUM_T_H3{x,y}]  = FrontFactor(H2_Comby, H3(:,x,y), W_H2_H3, Epsilon);

        dW_H2_H3  = dW_H2_H3  + DiffFactor(PropFactor{x,y}, T_H3{x,y}, SUM_T_H3{x,y}, F_H3{x,y});

        [PropFactor_New{x,y}] = PropagateFactor(T_H3{x,y}, SUM_T_H3{x,y}, PropFactor{x,y});

      end   
    end
    PropFactor = PropFactor_New;
    PropFactor_New = cell(size(H2,2),size(H2,3));   

    Temp = cell(size(H3,2),size(H3,3),5,5);
    for x0 = 1:1:size(H3,2)
      for y0 = 1:1:size(H3,3)

        Temp_APropFactor = PropFactor{x0,y0};

        for x = 1:1:5,
          for y = 1:1:5,
            Temp{x0,y0,x,y} = Temp_APropFactor(Sp_Offset_H2(x,y)+1:Sp_Offset_H2(x,y)+size(H2,1)) / numel(Sp_Offset_H2);
          end
        end
      end
    end
    PropFactor = Temp;

    X_Coord = zeros(size(PropFactor));
    Y_Coord = zeros(size(PropFactor));

    X_Vector = 1:1:size(H2,2);
    Y_Vector = 1:1:size(H2,3);

    XMatrix = repmat(X_Vector',[1 length(Y_Vector)]);
    YMatrix = repmat(Y_Vector,[length(X_Vector) 1]);

    for x0 = 1:1:size(H3,2)
      for y0 = 1:1:size(H3,3)
        XMatrix_Sub = XMatrix(x0+[0:1:4],y0+[0:1:4]);
        YMatrix_Sub = YMatrix(x0+[0:1:4],y0+[0:1:4]);

        for x = 1:1:5,
          for y = 1:1:5,
            X_Coord(x0,y0,x,y) = XMatrix_Sub(x,y);
            Y_Coord(x0,y0,x,y) = YMatrix_Sub(x,y);
          end
        end
      end
    end

  % H2

    PropFactor_New = cell(size(H3,2),size(H3,3),5,5);   

    for x0 = 1:1:size(H3,2)
      for y0 = 1:1:size(H3,3)
        for x = 1:1:5,
          for y = 1:1:5,
            X_Int = X_Coord(x0,y0,x,y);
            Y_Int = Y_Coord(x0,y0,x,y);
            H2_Select = H2(:,X_Int,Y_Int);

            H1_Sub = H1(:,(X_Int-1)*2+[1:2],(Y_Int-1)*2+[1:2]);

            H1_Flat = zeros(numel(H1_Sub),1);
            for x_s = 1:1:2,
              for y_s = 1:1:2,
                H1_Flat(((x_s-1)*2+(y_s-1))*NumberOfH_1+[1:NumberOfH_1]) = H1_Sub(:,x_s,y_s);
              end
            end

            [F_H2, T_H2, SUM_T_H2]  = FrontFactor(H1_Flat, H2_Select , W_H1_H2, Epsilon);
            dW_H1_H2  = dW_H1_H2  + DiffFactor(PropFactor{x0,y0,x,y}, T_H2, SUM_T_H2, F_H2);

            [PropFactor_New{x0,y0,x,y}] = PropagateFactor(T_H2, SUM_T_H2, PropFactor{x0,y0,x,y});
          end
        end
      end
    end
    PropFactor = PropFactor_New;

    PropFactor_New = cell(4*numel(PropFactor),3);   
    Counter = 0;
    for x0 = 1:1:size(H3,2)
      for y0 = 1:1:size(H3,3)
        for x = 1:1:5,
          for y = 1:1:5,
            Temp = PropFactor{x0,y0,x,y};
            X_Int = X_Coord(x0,y0,x,y);
            Y_Int = Y_Coord(x0,y0,x,y);

            for x_s = 1:1:2,
              for y_s = 1:1:2,
                Counter = Counter + 1;

                PropFactor_New{Counter,1} = Temp(((x_s-1)*2+(y_s-1))*NumberOfH_1+[1:NumberOfH_1]);
                PropFactor_New{Counter,2} = (X_Int-1)*2 + x_s;
                PropFactor_New{Counter,3} = (Y_Int-1)*2 + y_s;

              end
            end
          end
        end
      end
    end
    PropFactor = PropFactor_New;

    % H1
    for i = 1:1:size(PropFactor,1)
      [F_H1, T_H1, SUM_T_H1]  = FrontFactor(P_X(:,PropFactor{i,2},PropFactor{i,3}), H1(:,PropFactor{i,2},PropFactor{i,3}), W_X_H1, Epsilon);
      dW_X_H1  = dW_X_H1  + DiffFactor(PropFactor{i,1}, T_H1, SUM_T_H1, F_H1);
    end

  end
%end


eval(['save WSub/Temp_Iter' num2str(WeightIter) '_BlockID' num2str(BlockID) '.mat dW_X_H1 dW_H1_H2 dW_H2_H3 dW_H3_H4 dW_H4_H5 dW_H5_HY L']);
fprintf(['Done\n']);
   
```
