## Supplementary material for "Back-propagation learning in deep Spike-By-Spike networks": Source Code: README.txt

0.) Perform the stage A

1.) Copy the last weight over from stage A (e.g. W_X_H1_Iter150.mat) and rename it to W_X_H1_Iter0.mat. Create the other weight matrices:
Create_W_H1_H2
CreateInitW_H2_H3
Create_W_H3_H4
CreateInitW_H4_H5
CreateInitW_H5_HY

2.) Do the rest like in stage A.
