## Supplementary material for "Back-propagation learning in deep Spike-By-Spike networks": Source Code: README.txt

0.) Perform the stage B

1.) Copy the last weight over from stage A (e.g. W_X_H1_Iter150.mat) and rename it to W_X_H1_Iter0.mat.  
Copy the last weight over from stage B (e.g. W_H2_H3_Iter150.mat) and rename it to W_H2_H3_Iter0.mat.  

Create the other weight matrices:
Create_W_H1_H2
Create_W_H3_H4
CreateInitW_H4_H5
CreateInitW_H5_HY

2.) Do the rest like in stage B but with more learning steps (e.g. 200), due to the drop out procedure needs them.
