## Supplementary material for "Back-propagation learning in deep Spike-By-Spike networks": Source Code: mnist_deep_ourstruct.py

```
# Copyright 2015 The TensorFlow Authors. All Rights Reserved.
#
# Licensed under the Apache License, Version 2.0 (the "License");
# you may not use this file except in compliance with the License.
# You may obtain a copy of the License at
#
#     http://www.apache.org/licenses/LICENSE-2.0
#
# Unless required by applicable law or agreed to in writing, software
# distributed under the License is distributed on an "AS IS" BASIS,
# WITHOUT WARRANTIES OR CONDITIONS OF ANY KIND, either express or implied.
# See the License for the specific language governing permissions and
# limitations under the License.
# ==============================================================================

"""A deep MNIST classifier using convolutional layers.

See extensive documentation at
https://www.tensorflow.org/get_started/mnist/pros
"""
# Disable linter warnings to maintain consistency with tutorial.
# pylint: disable=invalid-name
# pylint: disable=g-bad-import-order

from __future__ import absolute_import
from __future__ import division
from __future__ import print_function

import argparse
import sys
import tempfile

from tensorflow.examples.tutorials.mnist import input_data

import tensorflow as tf

FLAGS = None


def deepnn(x):
  """deepnn builds the graph for a deep net for classifying digits.

  Args:
    x: an input tensor with the dimensions (N_examples, 784), where 784 is the
    number of pixels in a standard MNIST image.

  Returns:
    A tuple (y, keep_prob). y is a tensor of shape (N_examples, 10), with values
    equal to the logits of classifying the digit into one of 10 classes (the
    digits 0-9). keep_prob is a scalar placeholder for the probability of
    dropout.
  """
  # Reshape to use within a convolutional neural net.
  # Last dimension is for "features" - there is only one here, since images are
  # grayscale -- it would be 3 for an RGB image, 4 for RGBA, etc.
  with tf.name_scope('reshape'):
    x_image = tf.reshape(x, [-1, 28, 28, 1])

  # First convolutional layer - maps one grayscale image to 32 feature maps.
  with tf.name_scope('conv1'):
    W_conv1 = weight_variable([5, 5, 1, 32])
    b_conv1 = bias_variable([32])
    h_conv1 = tf.nn.relu(conv2d(x_image, W_conv1) + b_conv1)

  # Pooling layer - downsamples by 2X.
  with tf.name_scope('pool1'):
    h_pool1 = max_pool_2x2(h_conv1)

  # Second convolutional layer -- maps 32 feature maps to 64.
  with tf.name_scope('conv2'):
    W_conv2 = weight_variable([5, 5, 32, 64])
    b_conv2 = bias_variable([64])
    h_conv2 = tf.nn.relu(conv2d(h_pool1, W_conv2) + b_conv2)

  # Second pooling layer.
  with tf.name_scope('pool2'):
    h_pool2 = max_pool_2x2(h_conv2)

  # Fully connected layer 1 -- after 2 round of downsampling, our 28x28 image
  # is down to 7x7x64 feature maps -- maps this to 1024 features.
  with tf.name_scope('fc1'):
    W_fc1 = weight_variable([4 * 4 * 64, 1024])
    b_fc1 = bias_variable([1024])

    h_pool2_flat = tf.reshape(h_pool2, [-1, 4*4*64])
    h_fc1 = tf.nn.relu(tf.matmul(h_pool2_flat, W_fc1) + b_fc1)

  # Dropout - controls the complexity of the model, prevents co-adaptation of
  # features.
  with tf.name_scope('dropout'):
    keep_prob = tf.placeholder(tf.float32)
    h_fc1_drop = tf.nn.dropout(h_fc1, keep_prob)

  # Map the 1024 features to 10 classes, one for each digit
  with tf.name_scope('fc2'):
    W_fc2 = weight_variable([1024, 10])
    b_fc2 = bias_variable([10])

    y_conv = tf.matmul(h_fc1_drop, W_fc2) + b_fc2
  return y_conv, keep_prob


def conv2d(x, W):
  """conv2d returns a 2d convolution layer with full stride."""
  return tf.nn.conv2d(x, W, strides=[1, 1, 1, 1], padding='VALID')


def max_pool_2x2(x):
  """max_pool_2x2 downsamples a feature map by 2X."""
  return tf.nn.max_pool(x, ksize=[1, 2, 2, 1],
                        strides=[1, 2, 2, 1], padding='VALID')


def weight_variable(shape):
  """weight_variable generates a weight variable of a given shape."""
  initial = tf.truncated_normal(shape, stddev=0.1)
  return tf.Variable(initial)


def bias_variable(shape):
  """bias_variable generates a bias variable of a given shape."""
  initial = tf.constant(0.1, shape=shape)
  return tf.Variable(initial)


def main(_):
  # Import data
  mnist = input_data.read_data_sets(FLAGS.data_dir, one_hot=True)

  # Create the model
  x = tf.placeholder(tf.float32, [None, 784])

  # Define loss and optimizer
  y_ = tf.placeholder(tf.float32, [None, 10])

  # Build the graph for the deep net
  y_conv, keep_prob = deepnn(x)

  with tf.name_scope('loss'):
    cross_entropy = tf.nn.softmax_cross_entropy_with_logits(labels=y_,
                                                            logits=y_conv)
  cross_entropy = tf.reduce_mean(cross_entropy)

  with tf.name_scope('adam_optimizer'):
    train_step = tf.train.AdamOptimizer(1e-4).minimize(cross_entropy)

  with tf.name_scope('accuracy'):
    correct_prediction = tf.equal(tf.argmax(y_conv, 1), tf.argmax(y_, 1))
    correct_prediction = tf.cast(correct_prediction, tf.float32)
  accuracy = tf.reduce_mean(correct_prediction)

  graph_location = tempfile.mkdtemp()
  print('Saving graph to: %s' % graph_location)
  train_writer = tf.summary.FileWriter(graph_location)
  train_writer.add_graph(tf.get_default_graph())

  with tf.Session() as sess:
    sess.run(tf.global_variables_initializer())
    for i in range(20000):
      batch = mnist.train.next_batch(50)
      if i % 100 == 0:
        train_accuracy = accuracy.eval(feed_dict={
            x: batch[0], y_: batch[1], keep_prob: 1.0})
        print('step %d, training accuracy %g' % (i, train_accuracy))
      train_step.run(feed_dict={x: batch[0], y_: batch[1], keep_prob: 0.5})

    print('test accuracy %g' % accuracy.eval(feed_dict={
        x: mnist.test.images, y_: mnist.test.labels, keep_prob: 1.0}))

if __name__ == '__main__':
  parser = argparse.ArgumentParser()
  parser.add_argument('--data_dir', type=str,
                      default='./mnist/input_data',
                      help='Directory for storing input data')
  FLAGS, unparsed = parser.parse_known_args()
  tf.app.run(main=main, argv=[sys.argv[0]] + unparsed)
```
