## Supplementary material for "Back-propagation learning in deep Spike-By-Spike networks": Source Code: README.txt

We took the original source code, modfied it, ran it and got
test accuracy 0.9919

Modification are:

82c82
<     W_fc1 = weight_variable([7 * 7 * 64, 1024])
---
>     W_fc1 = weight_variable([4 * 4 * 64, 1024])
85c85
<     h_pool2_flat = tf.reshape(h_pool2, [-1, 7*7*64])
---
>     h_pool2_flat = tf.reshape(h_pool2, [-1, 4*4*64])
105c105
<   return tf.nn.conv2d(x, W, strides=[1, 1, 1, 1], padding='SAME')
---
>   return tf.nn.conv2d(x, W, strides=[1, 1, 1, 1], padding='VALID')
111c111
<                         strides=[1, 2, 2, 1], padding='SAME')
---
>                         strides=[1, 2, 2, 1], padding='VALID')
