## Supplementary material for "Back-propagation learning in deep Spike-By-Spike networks": Source Code: README.txt

We downloaded it from
https://raw.githubusercontent.com/tensorflow/tensorflow/r1.4/tensorflow/examples/tutorials/mnist/mnist_deep.py
, we ran it (Python 3.5.4, TensorFlow 1.4.1) and got 
test accuracy 0.9923.
