## Supplementary material for "Back-propagation learning in deep Spike-By-Spike networks": Source Code: cnn_mnist_nopad.py

```
#  Copyright 2016 The TensorFlow Authors. All Rights Reserved.
#
#  Licensed under the Apache License, Version 2.0 (the "License");
#  you may not use this file except in compliance with the License.
#  You may obtain a copy of the License at
#
#   http://www.apache.org/licenses/LICENSE-2.0
#
#  Unless required by applicable law or agreed to in writing, software
#  distributed under the License is distributed on an "AS IS" BASIS,
#  WITHOUT WARRANTIES OR CONDITIONS OF ANY KIND, either express or implied.
#  See the License for the specific language governing permissions and
#  limitations under the License.
"""Convolutional Neural Network Estimator for MNIST, built with tf.layers."""

from __future__ import absolute_import
from __future__ import division
from __future__ import print_function

import numpy as np
import tensorflow as tf

tf.logging.set_verbosity(tf.logging.INFO)


def cnn_model_fn(features, labels, mode):
  """Model function for CNN."""
  # Input Layer
  # Reshape X to 4-D tensor: [batch_size, width, height, channels]
  # MNIST images are 28x28 pixels, and have one color channel
  input_layer = tf.reshape(features["x"], [-1, 28, 28, 1])

  # Convolutional Layer #1
  # Computes 32 features using a 5x5 filter with ReLU activation.
  # Padding is added to preserve width and height.
  # Input Tensor Shape: [batch_size, 28, 28, 1]
  # Output Tensor Shape: [batch_size, 28, 28, 32]
  conv1 = tf.layers.conv2d(
      inputs=input_layer,
      filters=32,
      kernel_size=[5, 5],
      padding="valid",
      activation=tf.nn.relu)

  # Pooling Layer #1
  # First max pooling layer with a 2x2 filter and stride of 2
  # Input Tensor Shape: [batch_size, 28, 28, 32]
  # Output Tensor Shape: [batch_size, 14, 14, 32]
  pool1 = tf.layers.max_pooling2d(inputs=conv1, pool_size=[2, 2], strides=2)

  # Convolutional Layer #2
  # Computes 64 features using a 5x5 filter.
  # Padding is added to preserve width and height.
  # Input Tensor Shape: [batch_size, 14, 14, 32]
  # Output Tensor Shape: [batch_size, 14, 14, 64]
  conv2 = tf.layers.conv2d(
      inputs=pool1,
      filters=64,
      kernel_size=[5, 5],
      padding="valid",
      activation=tf.nn.relu)

  # Pooling Layer #2
  # Second max pooling layer with a 2x2 filter and stride of 2
  # Input Tensor Shape: [batch_size, 14, 14, 64]
  # Output Tensor Shape: [batch_size, 7, 7, 64]
  pool2 = tf.layers.max_pooling2d(inputs=conv2, pool_size=[2, 2], strides=2)

  # Flatten tensor into a batch of vectors
  # Input Tensor Shape: [batch_size, 7, 7, 64]
  # Output Tensor Shape: [batch_size, 7 * 7 * 64]
  pool2_flat = tf.reshape(pool2, [-1, 4 * 4 * 64])

  # Dense Layer
  # Densely connected layer with 1024 neurons
  # Input Tensor Shape: [batch_size, 7 * 7 * 64]
  # Output Tensor Shape: [batch_size, 1024]
  dense = tf.layers.dense(inputs=pool2_flat, units=1024, activation=tf.nn.relu)

  # Add dropout operation; 0.6 probability that element will be kept
  #dropout = tf.layers.dropout(
  #    inputs=dense, rate=0.4, training=mode == tf.estimator.ModeKeys.TRAIN)

  # Logits layer
  # Input Tensor Shape: [batch_size, 1024]
  # Output Tensor Shape: [batch_size, 10]
  #logits = tf.layers.dense(inputs=dropout, units=10)
  logits = tf.layers.dense(inputs=dense, units=10)

  predictions = {
      # Generate predictions (for PREDICT and EVAL mode)
      "classes": tf.argmax(input=logits, axis=1),
      # Add `softmax_tensor` to the graph. It is used for PREDICT and by the
      # `logging_hook`.
      "probabilities": tf.nn.softmax(logits, name="softmax_tensor")
  }
  if mode == tf.estimator.ModeKeys.PREDICT:
    return tf.estimator.EstimatorSpec(mode=mode, predictions=predictions)

  # Calculate Loss (for both TRAIN and EVAL modes)
  onehot_labels = tf.one_hot(indices=tf.cast(labels, tf.int32), depth=10)
  loss = tf.losses.softmax_cross_entropy(
      onehot_labels=onehot_labels, logits=logits)

  # Configure the Training Op (for TRAIN mode)
  if mode == tf.estimator.ModeKeys.TRAIN:
    optimizer = tf.train.GradientDescentOptimizer(learning_rate=0.001)
    train_op = optimizer.minimize(
        loss=loss,
        global_step=tf.train.get_global_step())
    return tf.estimator.EstimatorSpec(mode=mode, loss=loss, train_op=train_op)

  # Add evaluation metrics (for EVAL mode)
  eval_metric_ops = {
      "accuracy": tf.metrics.accuracy(
          labels=labels, predictions=predictions["classes"])}
  return tf.estimator.EstimatorSpec(
      mode=mode, loss=loss, eval_metric_ops=eval_metric_ops)


def main(unused_argv):
  # Load training and eval data
  mnist = tf.contrib.learn.datasets.load_dataset("mnist")
  train_data = mnist.train.images  # Returns np.array
  train_labels = np.asarray(mnist.train.labels, dtype=np.int32)
  eval_data = mnist.test.images  # Returns np.array
  eval_labels = np.asarray(mnist.test.labels, dtype=np.int32)

  # Create the Estimator
  mnist_classifier = tf.estimator.Estimator(
      model_fn=cnn_model_fn, model_dir="/tmp/mnist_convnet_model")

  # Set up logging for predictions
  # Log the values in the "Softmax" tensor with label "probabilities"
  tensors_to_log = {"probabilities": "softmax_tensor"}
  logging_hook = tf.train.LoggingTensorHook(
      tensors=tensors_to_log, every_n_iter=50)

  # Train the model
  train_input_fn = tf.estimator.inputs.numpy_input_fn(
      x={"x": train_data},
      y=train_labels,
      batch_size=100,
      num_epochs=None,
      shuffle=True)
  mnist_classifier.train(
      input_fn=train_input_fn,
      steps=20000,
      hooks=[logging_hook])

  # Evaluate the model and print results
  eval_input_fn = tf.estimator.inputs.numpy_input_fn(
      x={"x": eval_data},
      y=eval_labels,
      num_epochs=1,
      shuffle=False)
  eval_results = mnist_classifier.evaluate(input_fn=eval_input_fn)
  print(eval_results)


if __name__ == "__main__":
  tf.app.run()
```
