## Supplementary material for "Back-propagation learning in deep Spike-By-Spike networks": Source Code: README.txt

We took the original source code and modified it such that it represented our network structure. We ran it (Python 3.5.4, TensorFlow 1.4.1) and it resulted in an accuracy of 0.971.

The changes we made:
42c42
<       padding="same",
---
>       padding="valid",
60c60
<       padding="same",
---
>       padding="valid",
72c72
<   pool2_flat = tf.reshape(pool2, [-1, 7 * 7 * 64])
---
>   pool2_flat = tf.reshape(pool2, [-1, 4 * 4 * 64])
81,82c81,82
<   dropout = tf.layers.dropout(
<       inputs=dense, rate=0.4, training=mode == tf.estimator.ModeKeys.TRAIN)
---
>   #dropout = tf.layers.dropout(
>   #    inputs=dense, rate=0.4, training=mode == tf.estimator.ModeKeys.TRAIN)
87c87,88
<   logits = tf.layers.dense(inputs=dropout, units=10)
---
>   #logits = tf.layers.dense(inputs=dropout, units=10)
>   logits = tf.layers.dense(inputs=dense, units=10)


Well, there is also this difference between the original r1.12 source code and the code we used but we couldn't determine the origin anymore:
<   loss = tf.losses.sparse_softmax_cross_entropy(labels=labels, logits=logits)
---
>   onehot_labels = tf.one_hot(indices=tf.cast(labels, tf.int32), depth=10)
>   loss = tf.losses.softmax_cross_entropy(
>       onehot_labels=onehot_labels, logits=logits)
