## Supplemental information and pictures for "Back-propagation learning in deep Spike-By-Spike networks"

### Supplementary Material

#### 1 DERIVATION OF THE SBS BACKPROP LEARNING RULE

For derivation of the learning rule, the error function is defined by the cross entropy

$$E = -\sum_{\mu}^{M}\sum_{d}^{D} t_\mu(d)\log(h^t_{y,\mu}(d)) \tag{S1}$$

with $t_\mu(d)$ as the desired output for neuron $d$ in the output layer of the network and $h^t_{y,\mu}(d)$ as the actual output of the network at time $t$. It is assumed that the output layer has $D$ output neurons as well as that the error is optimized over an ensemble of $M$ patterns. The individual pattern is denoted by $\mu$. Furthermore, $\sum_d^D t_\mu(d) = 1$ and $\sum_d^D h^t_{y,\mu}(d) = 1$ is enforced.

In addition, the update rule for $h^t_{y,\mu}(d)$ (Ernst et al. (2007)) is given by

$$h^t_{y,\mu}(d) = \left(\frac{1}{1+\epsilon}\right)\cdot\left(h^{t-1}_{y,\mu}(d) + \epsilon\sum_s^S \frac{h^{t-1}_{y,\mu}(d)W^{y-1\to y}(s|d)}{\sum_j^D h^{t-1}_{y,\mu}(j)W^{y-1\to y}(s|j)}h^{t-1}_{y-1,\mu}(s)\right) \tag{S2}$$

with $t-1$ denoting latent variables one update step back in the past and $y-1$ for the layer directly before the output layer, which contains $S$ neurons. $W^{y-1\to y}(s|d)$ describe the weights between the layer before the output layer ($y-1$) and the output layer ($y$) itself. These weights, like all weights in a Spike-By-Spike (SbS) network, are normalized via $\sum_s W^{y-1\to y}(s|d) = 1$.

Using equation S2 in equation S1 leads to

$$E = -\sum_{\mu}^{M}\sum_{d}^{D} t_\mu(d)\log\left(\left(\frac{1}{1+\epsilon}\right)\cdot\left(h^{t-1}_{y,\mu}(d) + \epsilon\sum_s^S \frac{h^{t-1}_{y,\mu}(d)W^{y-1\to y}(s|d)}{\sum_j^D h^{t-1}_{y,\mu}(j)W^{y-1\to y}(s|j)}h^{t-1}_{y-1,\mu}(s)\right)\right) \tag{S3}$$

For optimizing the weights, the gradient $-\frac{\partial E}{\partial W^{y-1\to y}(u|v)}$ is of special interest. Part of this gradient is

$$\frac{\partial}{\partial W^{y-1\to y}(u|v)}\sum_s^S \frac{h^{t-1}_{y,\mu}(d)W^{y-1\to y}(s|d)}{\sum_j^D h^{t-1}_{y,\mu}(j)W^{y-1\to y}(s|j)}h^{t-1}_{y-1,\mu}(s) = \tag{S4}$$

$$\frac{h^{t-1}_{y,\mu}(v)h^{t-1}_{y-1,\mu}(u)}{\left(\sum_j^D h^{t-1}_{y,\mu}(j)W^{y-1\to y}(u|j)\right)^2}\left(\left(\delta_{d,v}\sum_{j'}^{D} h^{t-1}_{y,\mu}(j')W^{y-1\to y}(u|j')\right) - \left(h^{t-1}_{y,\mu}(d)W^{y-1\to y}(u|d)\right)\right)$$

with $\delta_{d,v}$ as Kronecker delta.

For keeping the further nomenclature shorter,

$$F_{b,\mu}(u,v) = \frac{h^t_{a,\mu}(u)h^t_{b,\mu}(v)}{\left(\sum_j h^t_{b,\mu}(j)W^{a\to b}(u|j)\right)^2} \tag{S5}$$

$$T_\mu(d) = \frac{t_\mu(d)}{h^t_{y,\mu}(d)} \tag{S6}$$

$$r_{y,\mu}(u,d) = h^t_{y,\mu}(d)W^{y-1\to y}(u|d) \tag{S7}$$

$$R_{y,\mu}(u) = \sum_j^D r_{y,\mu}(u,j) \tag{S8}$$

are introduced.

This leads to the following gradient for optimizing the weights $W^{y-1\to y}$:

$$-\frac{\partial E}{\partial W^{y-1\to y}(u|v)} = \frac{\epsilon}{1+\epsilon}\sum_\mu^M F_{y,\mu}(u,v)\cdot\sum_d^D T_\mu(d)\left(R_{y,\mu}(u)\delta_{d,v} - r_{y,\mu}(u,d)\right) \tag{S9}$$

For updating the weights one layer down $W^{y-2\to y-1}(u|v)$, in equation S3 $h^{t-1}_{y-1,\mu}(s)$ will be replaced by its update rule

$$h^{t-1}_{y-1,\mu}(s) = \left(\frac{1}{1+\epsilon}\right)\cdot\left(h^{t-2}_{y-1,\mu}(s) + \epsilon\sum_k^K \frac{h^{t-2}_{y-1,\mu}(s)W^{y-2\to y-1}(k|s)}{\sum_j^S h^{t-2}_{y-1,\mu}(j)W^{y-2\to y-1}(k|j)}h^{t-2}_{y-2,\mu}(k)\right) . \tag{S10}$$

This results in

$$\begin{aligned} E &= -\sum_\mu^M\sum_d^D t_\mu(d)\log\left(\left(\frac{1}{1+\epsilon}\right)\cdot\left(h^{t-1}_{y,\mu}(d) + \epsilon\sum_s^S\frac{h^{t-1}_{y,\mu}(d)W^{y-1\to y}(s|d)}{\sum_j^D h^{t-1}_{y,\mu}(j)W^{y-1\to y}(s|j)}\cdot\right.\right. \\ &\left.\left.\left(\frac{1}{1+\epsilon}\right)\cdot\left(h^{t-2}_{y-1,\mu}(s) + \epsilon\sum_k^K\frac{h^{t-2}_{y-1,\mu}(s)W^{y-2\to y-1}(k|s)}{\sum_j^S h^{t-2}_{y-1,\mu}(j)W^{y-2\to y-1}(k|j)}h^{t-2}_{y-2,\mu}(k)\right)\right)\right) . \end{aligned} \tag{S11}$$

This delivers the basis for calculating the gradient for the weight values $W^{y-2\to y-1}(u|v)$ as follows:

$$-\frac{\partial E}{\partial W^{y-2\to y-1}(u|v)} = \sum_\mu^M\sum_s\left(\frac{\partial h^{t-1}_{y-1,\mu}(s)}{\partial W^{y-2\to y-1}(u|v)}\right)\left(\sum_d \Psi_{y,\mu}(s,d)T_\mu(d)\right) \tag{S12}$$

with

$$\Psi_{y,\mu}(s,d) = \left(\frac{\epsilon}{1+\epsilon}\right)\frac{h^{t-1}_{y,\mu}(d)W^{y-1\to y}(s|d)}{\sum_j h^{t-1}_{y,\mu}(j)W^{y-1\to y}(s|j)} \tag{S13}$$

and

$$\frac{\partial h^{t-1}_{y-1,\mu}(s)}{\partial W^{y-2\to y-1}(u|v)} = \left(\frac{\epsilon}{1+\epsilon}\right) \frac{\partial}{\partial W^{y-2\to y-1}(u|v)} \sum_k^K \frac{h^{t-2}_{y-1,\mu}(s) W^{y-2\to y-1}(k|s)}{\sum_j^S h^{t-2}_{y-1,\mu}(j) W^{y-2\to y-1}(k|j)} h^{t-2}_{y-2,\mu}(k)$$

$$= \left(\frac{\epsilon}{1+\epsilon}\right) F_{y-1,\mu}(u,v) \left(R_{y-1,\mu}(u)\delta_{s,v} - r_{y-1,\mu}(u,s)\right) . \tag{S14}$$

This procedure can be iterated until the input layer is reached. This delivers the gradient for the weights

$$-\frac{\partial E}{\partial W^{y-m-1\to y-m}(u|v)} = \sum_\mu^M F_{y-m,\mu}(u,v) \cdot$$

$$\sum_{q_{y-m}}^{Q_{y-m}} \left(R_{y-m,\mu}(u)\delta_{q_{y-m},v} - r_{y-m,\mu}(u,q_{y-m})\right) \Phi_{y-m,\mu}(q_{y-m}) \tag{S15}$$

with for $m > 0$

$$\Phi_{y-m,\mu}(q_{y-m}) = \left(\frac{\epsilon}{1+\epsilon}\right) \sum_{q_{y-m+1}}^{Q_{y-m+1}} \Phi_{y-m+1,\mu}(q_{y-m+1}) \frac{h^{t-m}_{y-m+1,\mu}(q_{y-m+1}) W^{y-m\to y-m+1}(q_{y-m}|q_{y-m+1})}{\sum_j h^{t-m}_{y-m+1,\mu}(j) W^{y-m\to y-m+1}(q_{y-m}|j)} \tag{S16}$$

and for $m = 0$

$$\Phi_{y,\mu}(q_y) = \left(\frac{\epsilon}{1+\epsilon}\right) \frac{t_\mu(q_y)}{h^t_{y,\mu}(q_y)} \tag{S17}$$

as well as

$$\begin{aligned} F_{y-m,\mu}(u,v) &= \frac{h^{t-m-1}_{y-m-1,\mu}(u) h^{t-m-1}_{y-m,\mu}(v)}{\left(\sum_j h^{t-m-1}_{y-m,\mu}(j) W^{y-m-1\to y-m}(u|j)\right)^2} \\ r_{y-m,\mu}(u,q_{y-m}) &= h^{t-m-1}_{y-m,\mu}(q_{y-m}) W^{y-m-1\to y-m}(u|q_{y-m}) \\ R_{y-m,\mu}(u) &= \sum_j^{Q_{y-m}} r_{y-m,\mu}(u,j) . \end{aligned}$$

Like in other backprop rules, the error is back-propagated from the output layer to the input layer but here also in time (one iteration back in time for each layer closer to the input).

Using this gradient, the weights in all the examples in this paper are updated as follows

$$\begin{aligned} V^{y-m-1\to y-m}(u|v) &= W^{y-m-1\to y-m}(u|v) \left(1 - \frac{\gamma}{S} \frac{\partial E}{\partial W^{y-m-1\to y-m}(u|v)}\right) \\ W^{new,y-m-1\to y-m}(u|v) &= \frac{V^{y-m-1\to y-m}(u|v)}{\sum_j V^{y-m-1\to y-m}(j|v)} \end{aligned} \tag{S18}$$

with $0 < \gamma < 1$ as learning rate and

$$S = \max\left(\left|\frac{\partial E}{\partial W^{y-m-1 \to y-m}(u|v)}\right|\right) \tag{S19}$$

as scaling factor for ensuring the non-negativity of the weight values.

#### 2 XOR: LEARNING THE WEIGHTS

In the main paper, we showed that it is possible to use the SbS error back-propagation rule to successfully learn the weights for a XOR network. However, the weights learned by the network (see figure S1a) are far away from ideal binary weights (see figure 3a in the main paper). The network compensates these difference with its in-build competition within an IP. In the following we wanted to know if the weights would approach the ideal weights more if the task requires the network to be more robust against noise in its information processing. Thus we reduced the number of spikes from $N_{Spikes} = 1024$ (as well as $\gamma = 0.025$) by half after 100 ($N_{Spikes} = 512$, $\gamma = \frac{0.025}{2}$), 350 ($N_{Spikes} = 256$, $\gamma = \frac{0.025}{4}$) and 600 ($N_{Spikes} = 128$, $\gamma = \frac{0.025}{8}$) learning steps (denoted by the vertical dashed lines in figure S1b, c, and d). Figure S1b shows the quality of the output during learning. As measure the Kullback-Leibler (KL) divergence between the expected output and the real output is calculated:

$$KL = \sum_{\mu}^{M} \sum_{d}^{D} t_{\mu}(d) \log\left(\frac{t_{\mu}(d)}{h_{HY,\mu}(d)}\right) \tag{S20}$$

Compared to the cross entropy measure, which was used for developing the SbS backprop rule, the KL adds a reference which results in zero if and only if both distributions are the same. However, due to limiting the smallest weight value to $\Theta = 0.0001$, it is not possible to reach a KL divergence of 0 by design. Here the smallest possible KL value is $\approx 4 \cdot 10^{-15}$. Similar behavior of the KL can be seen for the other examples with Boolean functions. Due to the increased amount of noise induced by reducing the number of spikes, the KL doesn't decrease as much as before. However, the weights values start to spread over a larger range of values. In the end the weights are approaching the ideal weights. Figure S1a shows the development of an example of weights over the learning procedure.

A note on measuring the range over which the weights spread: We selected to calculate the maximum and minimum of the weight matrix and then average these values over the 250 initial conditions. The averaging has to be done as second step because the assignment of the hidden neurons and the four input pattern is randomly permutated for every initial condition. Furthermore, this assignment can switch during learning. This is especially true until the correct set of weights was found by the network. In the end, this prevented us from just averaging the weight matrices over the 250 initial seeds without calculating the maximum and minimum first.

#### 3 XOR: IGNORING IRRELEVANT INFORMATION

In a second test with the XOR function we investigated if the SbS backprop rule can be used to learn a task correctly even if part of the input is not contributing to solving the task. Instead this part of the input introduces random unrelated information into the system that needs to be ignored. Thus we revisited the XOR network and added one random bit $J$ to the input pattern. Every time the XOR input bit pattern is changed also this bit is set to a new random value (equal random chance for zero or one). Figure S2a shows the modified network structure. The same learning procedure as well as the same analysis from the XOR example was performed.

The result is that the weights between the two neurons encoding bit $J$ and the hidden neurons show roughly the same values (see figure S2b and c for example weights). In terms of the winner-take-all competition inside the hidden layer, a uniform input is ignored due to the normalization of the latent variables. The development of KL divergence and the used weight value range (see figure S2d and e) shows similarities to figure S1, albeit slower ($1/3$ of all input spikes are lost to the meaningless bit $J$) and noisier. After 28 learning steps the output of the network shows no error (see figure S2f).

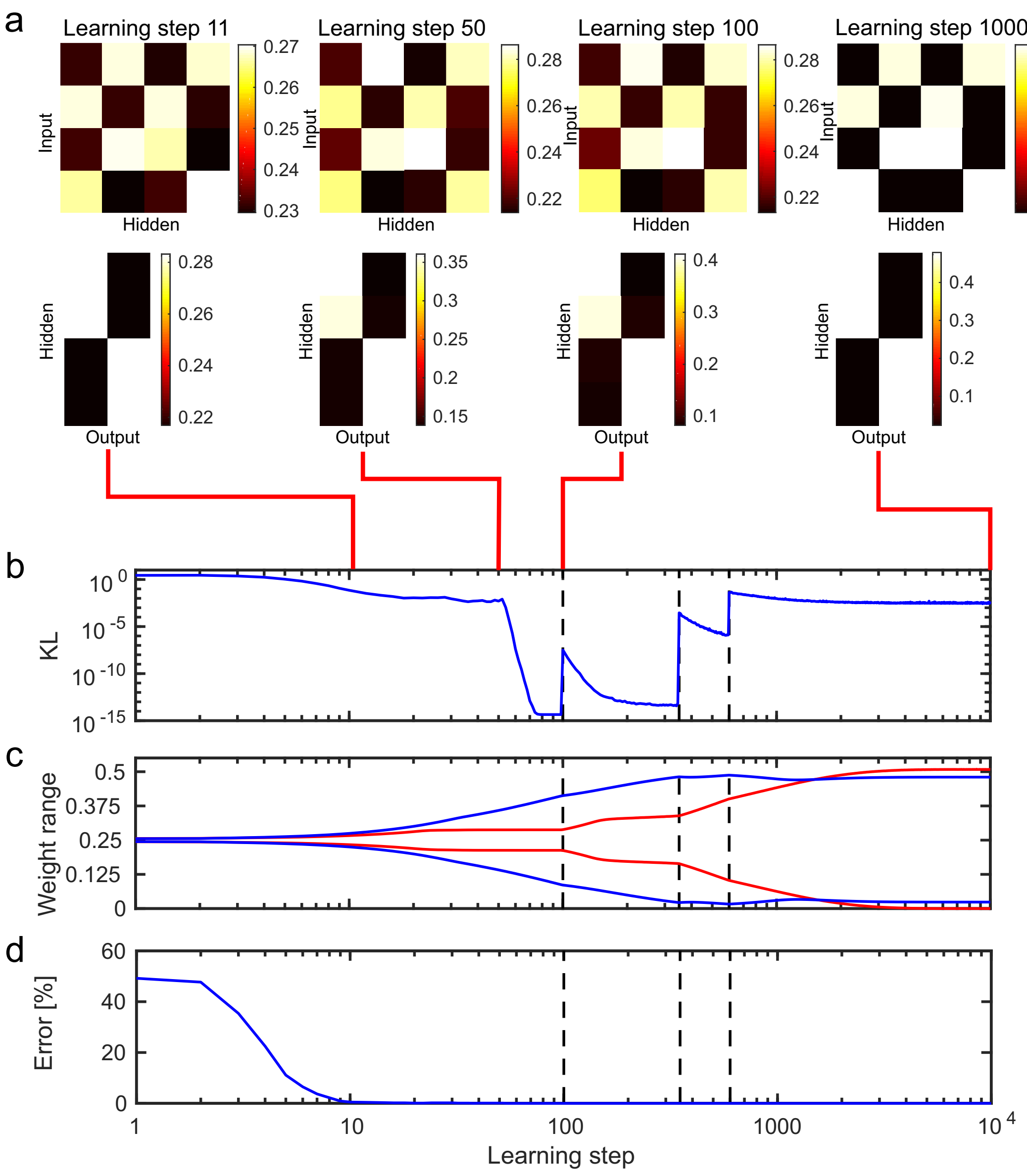


**Figure S1. Learning the weights for the SbS XOR network** a.) One example for the weights after increasing number of learning steps (i.e. after 11, 50, 100, and 10000 learning steps). At learning step zero, the weights are random. The upper row shows the weights between the input layer and hidden layer, while the lower row shows the weights between the hidden and output layer. The difference in the weight sets between learning steps condenses mainly in the range over which the weight values are spread. b.) The average distance between the target pattern and output of the hidden layer is shown. As measure the Kullback-Leibler divergence is used and it was averaged over 250 initial seeds. The vertical dashed lines show when parameters (number of spikes $N_{Spikes}$ and learning rate $\gamma$) of the network are changed. These parameter changes are done to increase the range of used weight values. c.) shows the weight range (maximum and minimum taken from the individual weight matrices and then averaged over the 250 initial conditions; red lines: weights between the input and the hidden layer, blue lines: weights between the hidden and output layer) in dependence of the learning step. d.) Error of the output of the XOR network averaged over the 250 initial conditions. The amount of spikes used for measuring the performance is the same as used for learning.

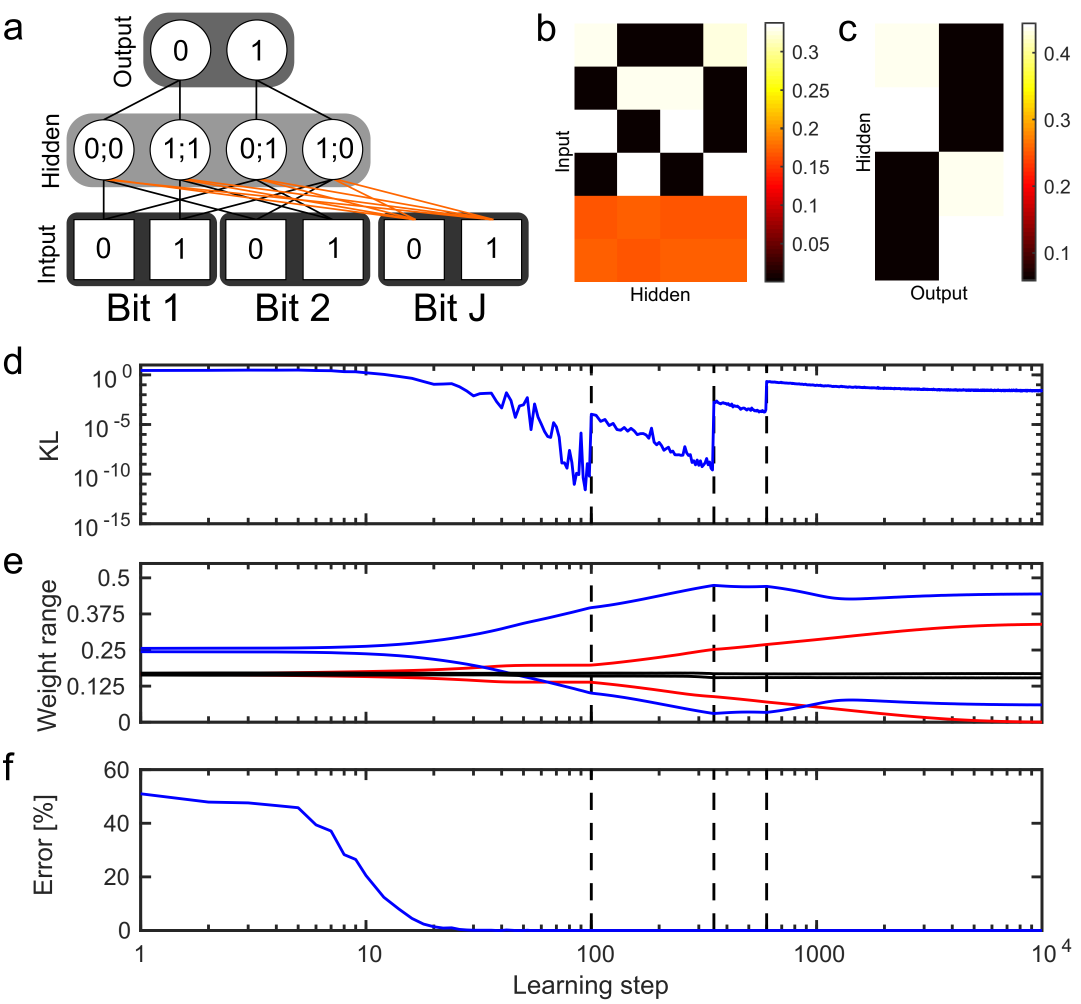


**Figure S2. Learning the XOR function with an additional uncorrelated input bit.** a.) Similar to figure S1. However, in this instance, two additional input neurons are present which represent an input bit $J$ that is uncorrelated to the XOR task. Applying the same learning procedure shown in figure S1, the learning process results in example weights that are shown in b.) and c.). The main difference is that the weight values, corresponding to the bit $J$, deliver the same input to all the neurons in the hidden layer. Thus the competition between the hidden neurons ignores this kind of irrelevant input. d.) shows the KL averages over 250 initial conditions. e.) Range of the weights over the learning steps: The additional black lines represent the development of the minimum and the maximum of the weight values corresponding to bit $J$ and the red lines represent the minimum and maximum of the weight values for bit 1 and bit 2. The blue line shows the range of the weights between the hidden and output layer. All these lines are the average over the 250 initial conditions. f.) Error of the output of the network averaged over the 250 initial seeds.

### 4 LEARNING THE 4 BIT PARITY FUNCTION

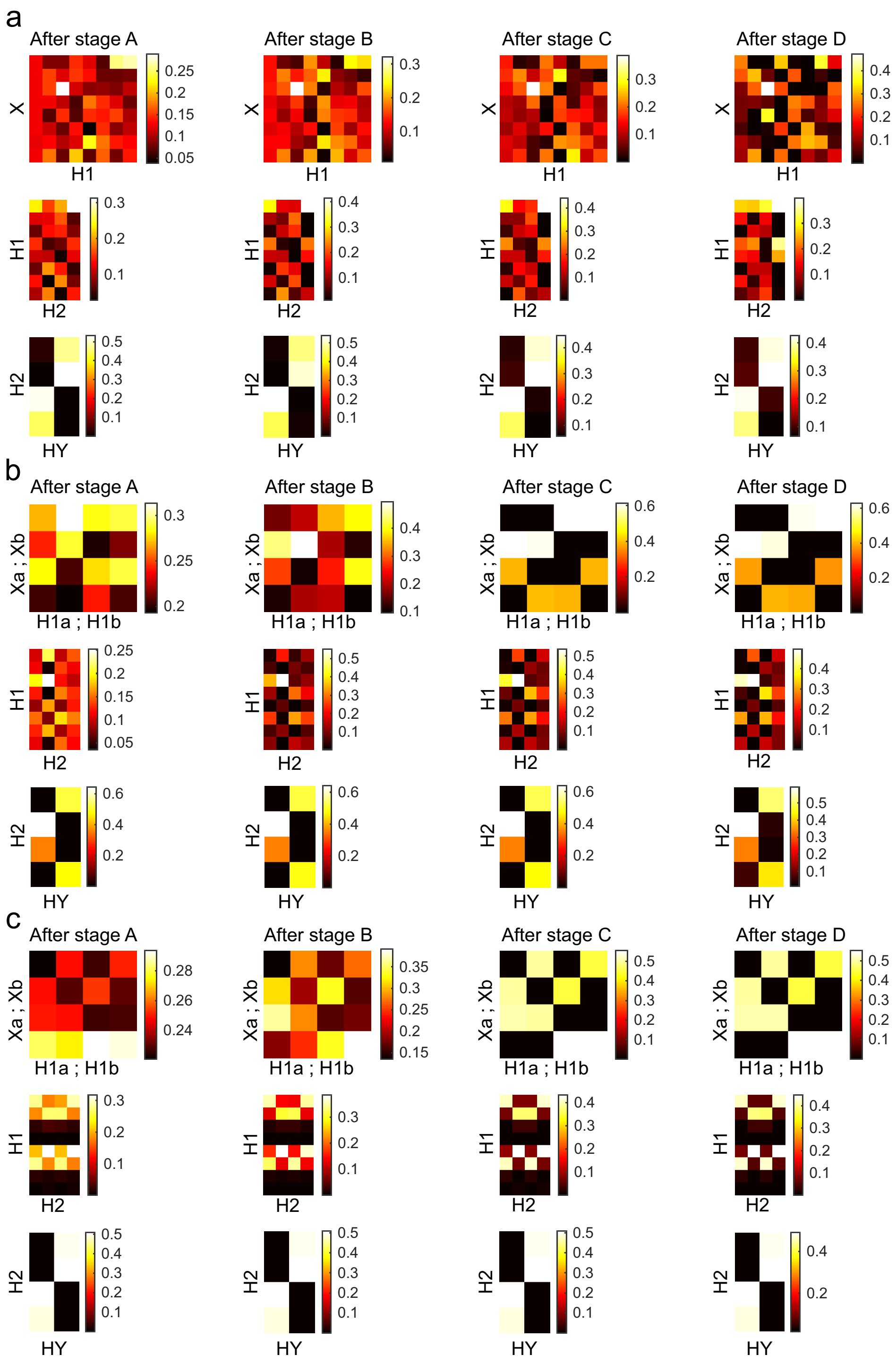


**Figure S3. Example weights for the 4 bit parity function after the four stages during learning.** a.) used the network shown in figure 4a of the main paper. b.) & c.) use the network shown in figure 4b, while c.) is only using information from the present state of the latent variables and not the past like a.) & b.).

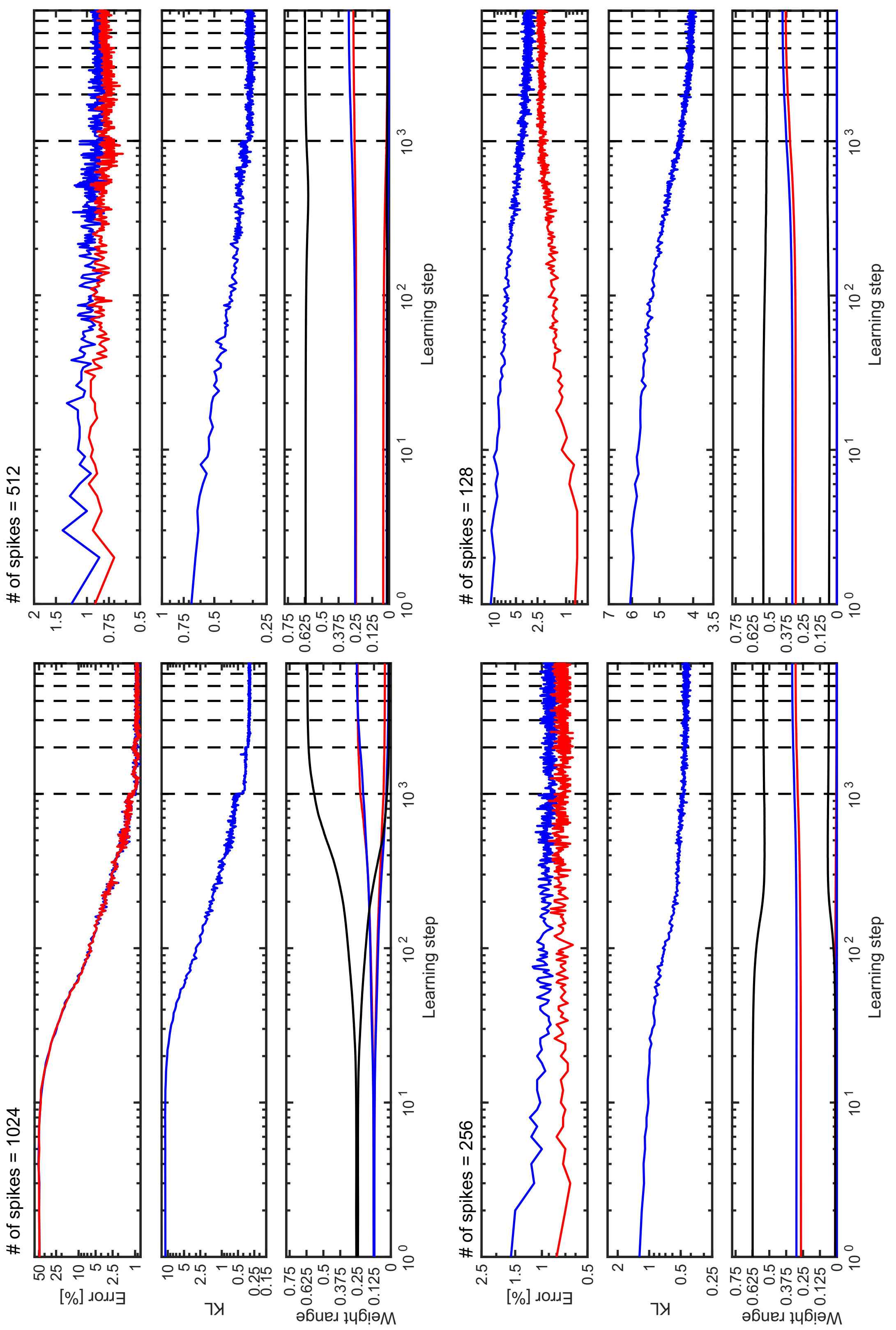


**Figure S4. Learning the 4 bit parity function (see figure 4a in the main paper).** For the four stages of the learning process a.) the average error of the output (like in S1d), b.) the Kullback-Leibler- divergence between the output of the output layer and the desired output (like in S1b), and c.) the range of the weights (like in S1c) are shown. The curves are averaged over 250 initial seeds and the black dashed lines symbolize the parameter changes. a.) red: performance measured with 1024 spikes. blue: performance tested with the number of spikes used for learning. c.) red: $W^{X\to H1}$, blue: $W^{H1\to H2}$, black: $W^{H2\to HY}$

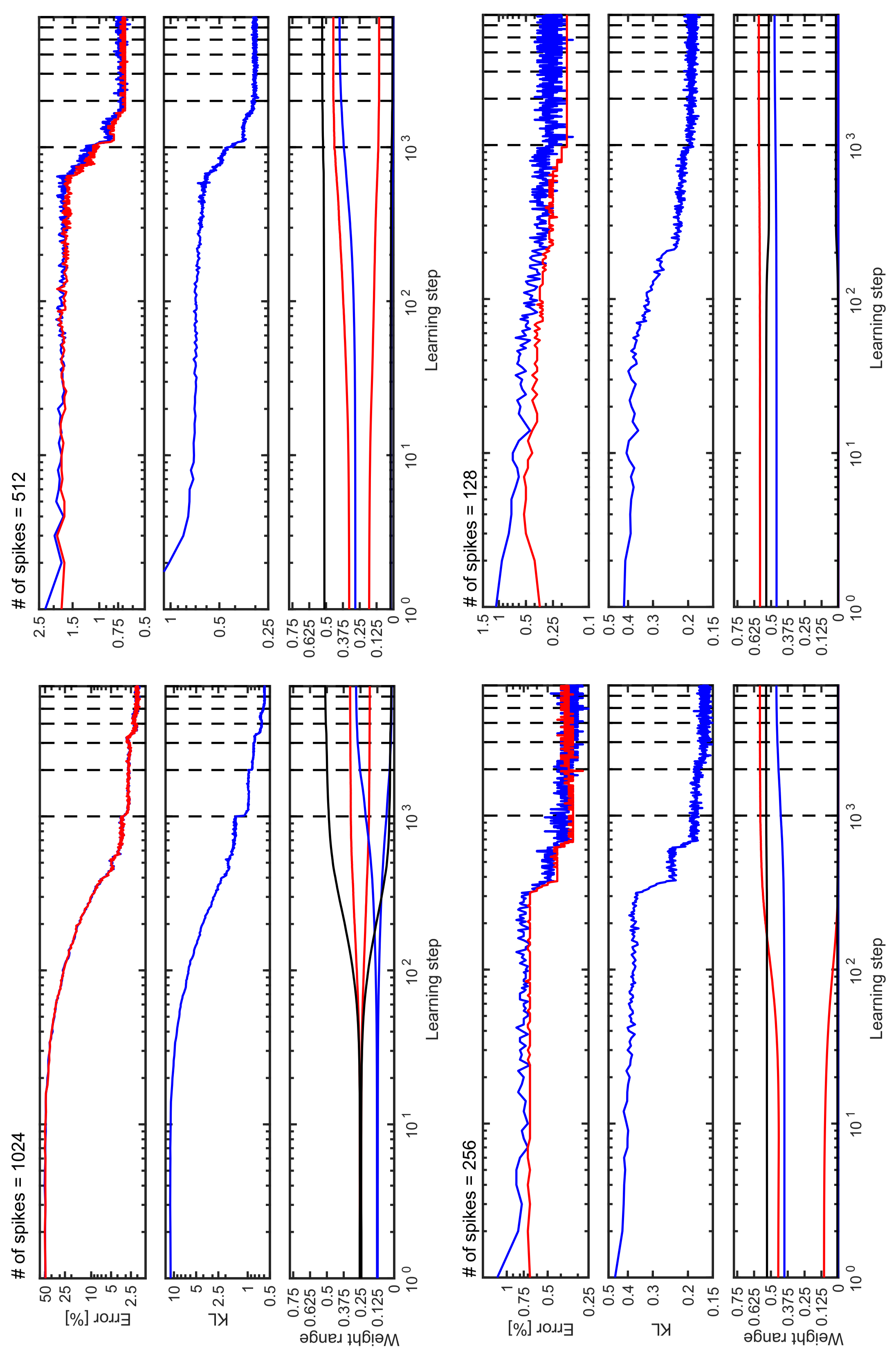


**Figure S5. Learning the 4 bit parity function with a split in $H1$ and in the input layer (see figure 4b in the main paper).** See figure S4 for details.

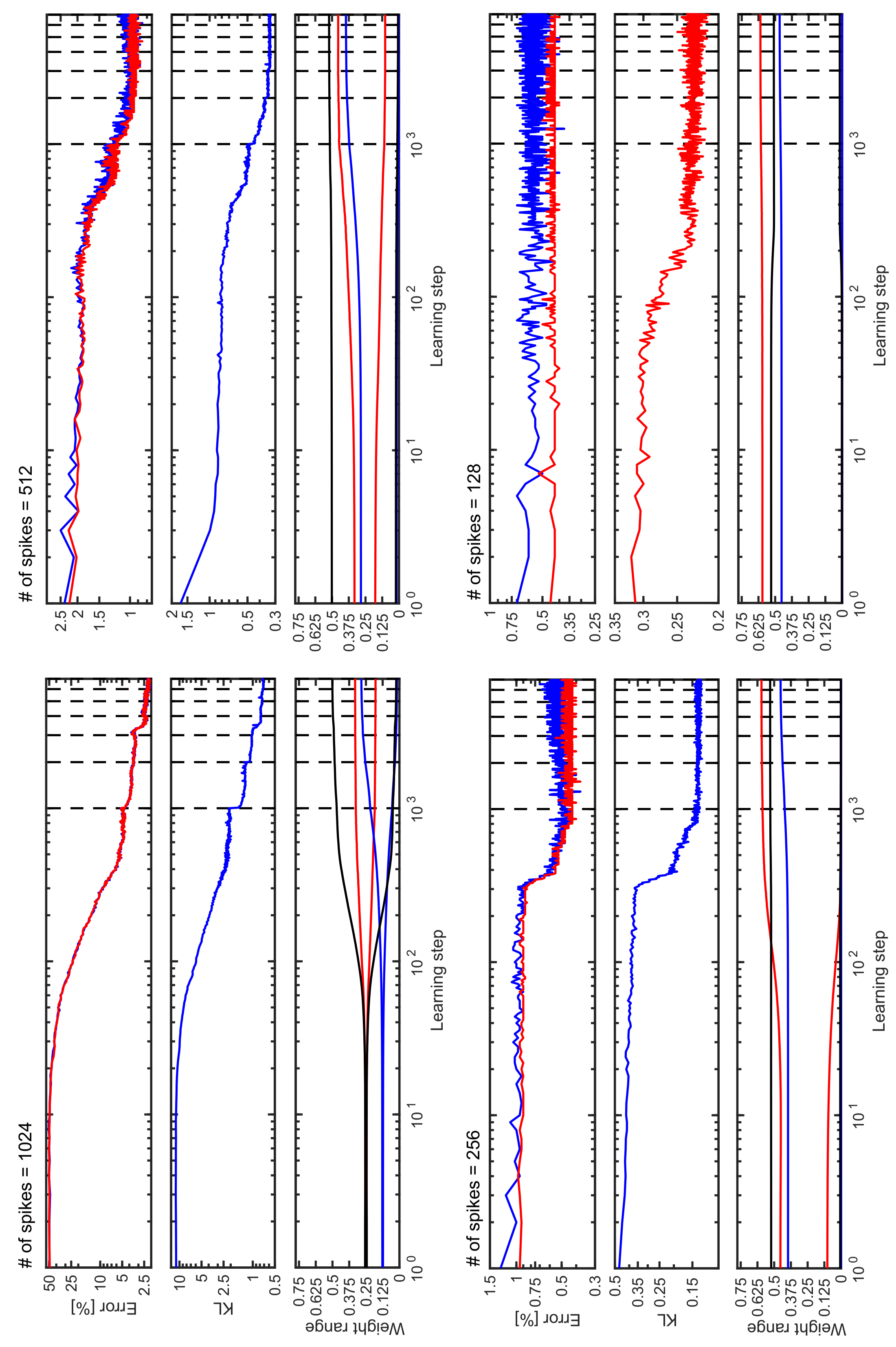


**Figure S6. Learning the 4 bit parity function with a split in $H1$ and in the input layer (see figure 4b in the main paper), using only the latent variables from the last spike.** See figure S4 for details.

#### 5 DEEP CONVOLUTIONAL NETWORK (MNIST)

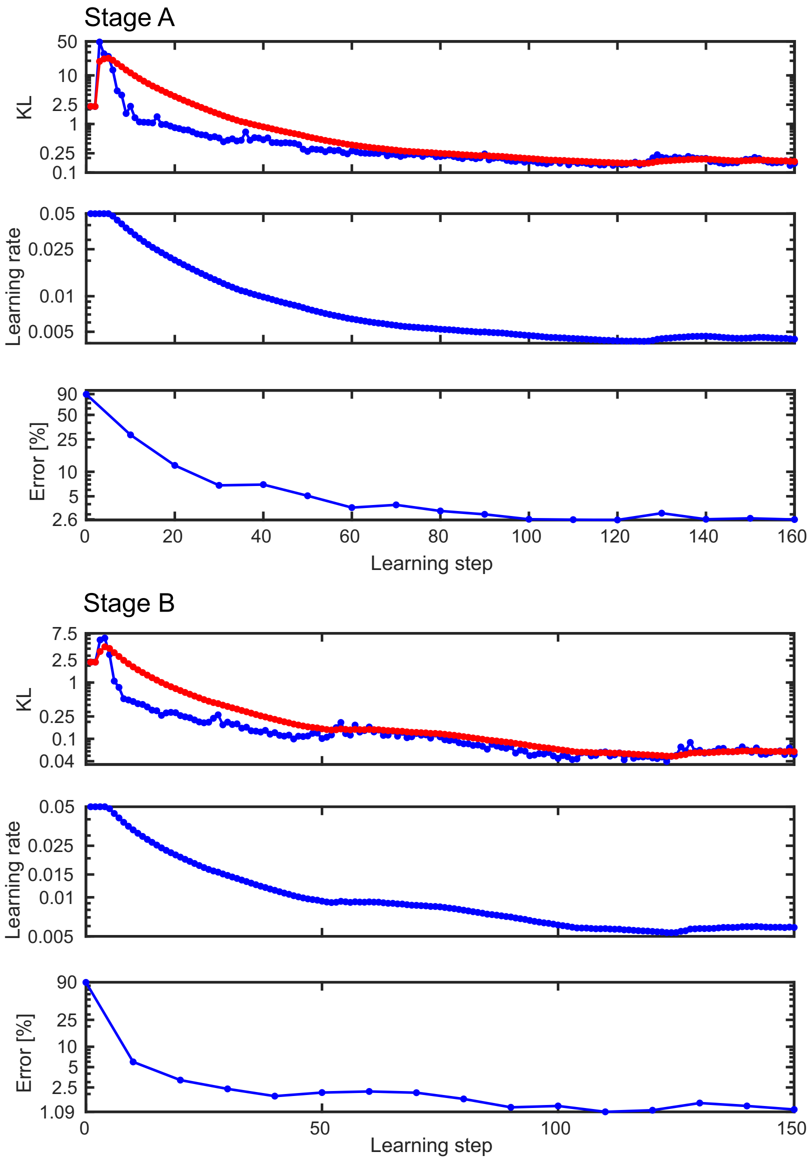


**Figure S7. Learning stage A & B for the MNIST SbS network.** Shown is the training error over the learning steps (measured as Kullback- Leibler divergence (KL) between the latent variables of the output layer and the desired output), where the blue curve shows the KL for the actual mini-batch and the red curve shows the low-pass filtered version of the KL. Furthermore, the learning rate – which is derived from the low-pass filtered KL – and the classification error on the test data set are shown over the learning steps.

Details concerning the network structure for the MNIST SbS network:

**Input layer $X$:** For the input a so called on/off split was made (Ernst et al. (2007)), which results in two channels per pixel. This is very similar to the representation of a bit by two neurons in the XOR example. This transformation can be described by $I_{ON}(x, y) = f(2P(x, y) - 1)$ and $I_{Off}(x, y) = f(1 - 2P(x, y))$ with $f(\cdot)$ as a threshold linear function which sets all negative values to 0 and passes on all positive values without change. $P(x, y)$ is the pixel values at the position $x$ and $y$ with $P(x, y) \in [0, 1]$. Next we used a 5 x 5 selection window and moved it with stride 1 in $x$ and $y$ over $I_{ON}(x, y)$ and $I_{Off}(x, y)$. This resulted in

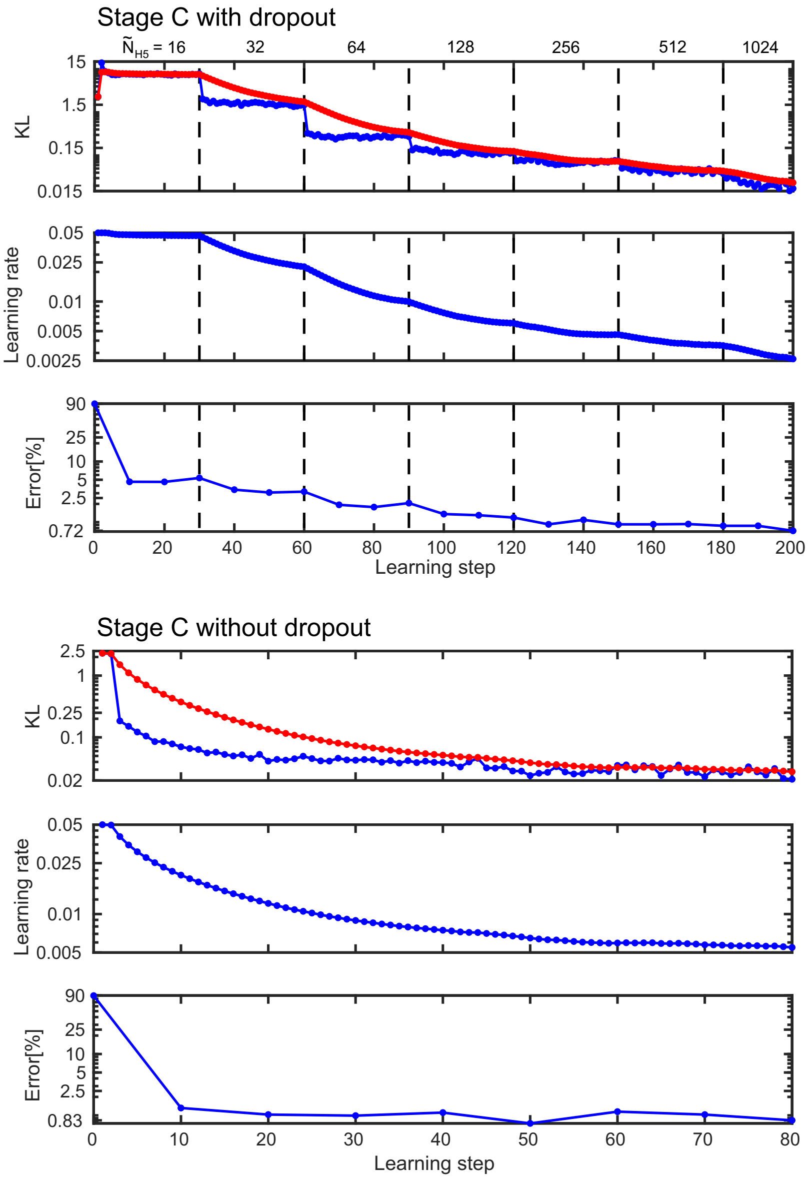


**Figure S8. Learning stage C for the MNIST SbS network.** Shown is the training error over the learning steps (measured as Kullback- Leibler divergence (KL) between the latent variables of the output layer and the desired output), where the blue curve shows the KL for the actual mini-batch and the red curve shows the low-pass filtered version of the KL. Furthermore, the learning rate – which is derived from the low-pass filtered KL – and the classification error on the test data set are shown over the learning steps. The upper set of three plots results from using a drop out procedure on $H5$ while the lower three plots are the result from training without dropout.

24 x 24 groups of with 50 values each. In a last step we normalized the values in every one of these groups individually to get 24 x 24 input probability distributions $p_X(s, x, y)$ (with $s \in [1, 50]$ and $\sum_s p_X(s, x, y)$ = 1). In every time step, one spike is drawn from each of these input probability distributions generating 24 x 24 input spikes per time step which are sent to layer $H1$.

**Convolution layer $H1$:** The hidden layer $H1$ consists of 24 x 24 independent normalization groups with 32 neurons per group each ($h_1(i, x, y)$ with $i \in [1, 32]$ and $\sum_i h_1(i, x, y, ) = 1$). These 24 x 24 normalization groups share the same weights. Every one of these $H1$ groups receives spikes from its direct counterpart from the input layer at the same $x$ and $y$ coordinates (The 5 x 5 convolution with stride one shown in figure 5 of the main paper is executed by building 24 x 24 groups from the original picture in input layer $X1$.). In every time step, one spike each is drawn from every one of the 24 x 24 groups, delivering 24 x 24 spikes to layer $H2$.

**Pooling layer $H2$:** Instead of using max functions, the pooling layer uses only the inherent competition implemented by the SbS update rule. This is done via the kernel's 2 x 2 weight matrix $W^{H1\rightarrow H2}(s,x,y|i) = \frac{\delta_{s,j}}{4}$ (with $x \in [1,2]$ and $y \in [1,2]$) using the h-update rule.

Or in other words: The weight matrix for the pooling layer has three dimensions on the input side ($s$,$x$,$y$) and one dimension on the output side ($i$). $i$ is the neuron index of the 'spike-receiving' neuron in the SbS inference population. $s$ is the neuron index of the 'spike-emitting' neuron in the 'input' SbS inference population. $x$ and $y$ refer to the spatial positioning of the 'input' SbS inference population in the 2 x 2 spatial kernel of the weights for the pooling layer. The equations for the weights describes that the weights are the same for all four $x$ & $y$ combinations. Furthermore, everywhere where $s$ is not equal to $i$ the weight value is zero. When $s$ is equal to $i$ then the weight value is set to $1/4$.

These weights are fixed and not learned. A 2 x 2 spatial patch from $H1$ with its 2 x 2 x 32 neurons delivers input to one normalization group in $H2$ which has also 32 neurons. The structure of the weights ensures that the input from different features (i.e. the 32 features that are represented by the 32 neurons in the normalization groups) do not mix. Thus the combined spatial inputs from the 32 features compete against each other. Only the features with strong inputs are represented in the corresponding $H2$ normalization group. This pooling kernel is applied with stride 2, which results in 12 x 12 spatially non-overlapping normalization groups with 32 neurons each. The next layer $H3$ gets 12 x 12 spikes per time step as input.

**Convolution layer $H3$:** While the convolution in $H1$ was simply done by reorganizing the input, in layer $H3$ the convolution is realized by using a 5 x 5 spatial region of layer $H2$ as input. This funnels the spikes generated from these 5 x 5 x 32 neurons from $H2$ into one normalization group of $H4$ with 64 neurons. Due to a stride of one, this results in a total of 8 x 8 normalization groups in $H3$. Thus layer $H3$ delivers 8 x 8 spikes to layer $H4$. The weight matrix is shared among all the normalization groups in this layer.

For making the difference between the convolutional layers $H1$ and $H3$ more clear: $H3$ is handled like a usual convolutional layer in a convolutional network: One SbS IP gets spikes from a spatial patch of 5 x 5 SbS IPs as input. These weights are used at all valid spatial positions.

Due to computational restrictions, we had to do the following trick for $H1$: Instead of using the spikes from a spatial 5 x 5 patch from the input populations – which are 25 spikes for every time step in the simulation –, we re-arranged the input instead. We created new input populations out of these 5 x 5 patches (we used the 5 x 5 spatial window, slid it over the input populations, and made new 25x bigger input populations out of it) from which we draw one spike per simulation time step. By using this input re-organization, we reduce the computational effort by a factor of 25.

On in short: In $H3$ we did the convolution on the incoming spikes. In $H1$ we did the convolution on the input pattern distribution before drawing the spikes.

**Pooling layer $H4$:** Layer $H4$ is similar to layer $H2$. The difference is that in $H4$ there are 64 neurons in each normalization group. $H4$ consists of 4 x 4 normalization groups, delivering 4 x 4 spikes per time step to layer $H5$.

**Fully connected layer $H5$:** Layer $H5$ is one large normalization group with e.g. 1024 neurons, which generates one spike per time step. It processes all spikes produced by the whole layer $H4$.

**Output layer $HY$:** The output layer is one normalization group with 10 neurons, where each neuron represents one class of the handwritten digits. This group takes the spikes from layer $H5$ as its input and updates its h-values according the h update rule. For decoding the results of the classification, the neuron with the highest h value is selected. The corresponding digits connected to this neuron is used as result of the classification task.
